# Physical Network Constraints Define the Multiplicative Architecture of the Brain’s Connectome

**DOI:** 10.1101/2025.02.27.640551

**Authors:** Ben Piazza, Dániel L. Barabási, Giulia Menichetti, André Ferreira Castro, Albert-László Barabási

## Abstract

The brain has long been conceptualized as a network of neurons connected by synapses. However, attempts to describe the connectome using established network science models have yielded conflicting outcomes, leaving the architecture of neural networks unresolved. Here, by performing a comparative analysis of eight experimentally mapped connectomes, we find that their degree distributions cannot be captured by the well-established random or scale-free models. Instead, the node degrees and strengths are well approximated by lognormal distributions, although these lack a mechanistic explanation in the context of the brain. By acknowledging the physical network nature of the brain, we show that neuron size is governed by a stochastic multiplicative process, which allows us to analytically derive the lognormal nature of the neuron length distribution. Our framework not only predicts the degree and strength distributions across each of the eight connectomes, but also yields a series of empirically falsifiable relationships between different neuron characteristics. The resulting multiplicative network represents a novel architecture for network science, whose distinctive quantitative features bridge critical gaps between neural structure and function, with implications for brain dynamics, robustness, and synchronization.

---

Since Ramón y Cajal revealed the first images of neurons in 1888, the brain has been conceptualized as a network, whose nodes represent individual neurons and links encode the synapses connecting them (Fig. 1a). However, it took more than a century to map this network, starting with the map of the *C. elegans* nematode four decades ago^1^, and culminating with the recent reconstruction of the full brain of the fruit fly^2–4^. Yet these maps represent only the first step of the journey — to uncover the alterations underlying brain disorders, from autism to schizophrenia, they must be complemented by a quantitative understanding of the connectome’s wiring principles. Indeed, advances in network science have demonstrated how the structural features define multiple functional characteristics of a network, from its robustness to node and link failures^5,6^, to its resilience^7^, information spread^8^, controllability^9,10^, and synchronization^11^. These advances have inspired multiple attempts to elucidate the structural characteristics of neural networks^12^, with conflicting outcomes: while the wiring of the *C. elegans* connectome was reported to be consistent with a randomly growing network^13^, co-activity networks extracted from human fMRI data have repeatedly demonstrated a scale-free organization^14–17^, two irreconcilable architectures. The recent proliferation of connectome data has also yielded puzzling outcomes: although the degree distributions do not exhibit a clear power-law scaling^18–20^, the observed prevalence of highly connected neurons^19–22^ remains a hallmark typically associated with scale-free networks^23^.

**Figure 1.**
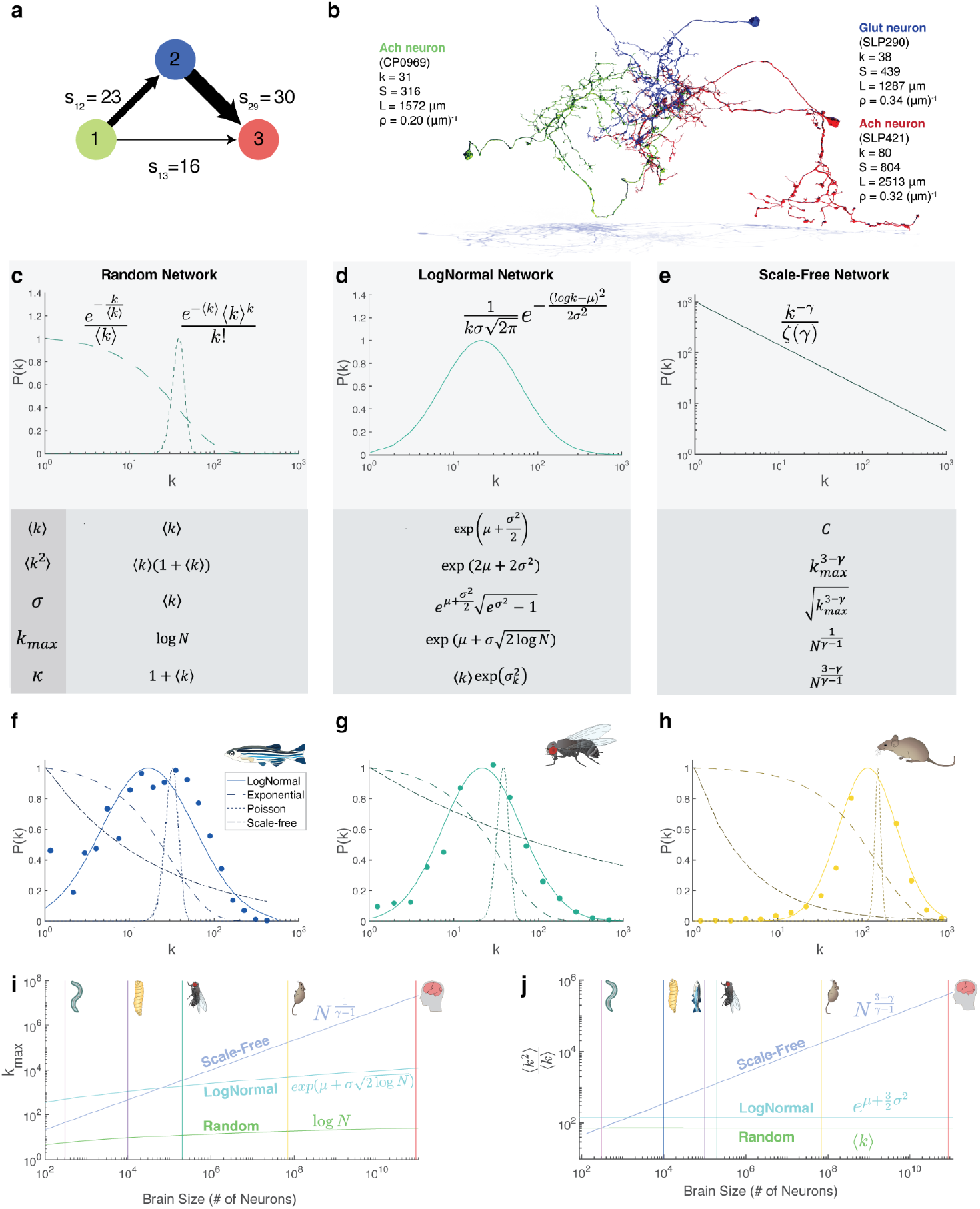
The Connectome and Network Science. **(a)** The traditional network representation of the connectome views the neurons as separate nodes (circles), connected by links whose weights correspond to the number of synapses between them. **(b)** The connectome is a physical network, as each neuron has a three dimensional spatial structure, as illustrated here for three neurons from the fruit fly (Flyware dataset). Each neuron is characterized by the length of its skeleton (*L*_*i*_), degree (*k*_*i*_), total number of synapses (*S*_*i*_), and synapse density (ρ_*i*_). **(c)** On log-linear scale a static random network has a narrow Poisson degree distribution, whose *<k>, <k*^2^*>*, σ_*k*_, expected maximum degree *k*_*max*_ and degree heterogeneity κ are analytically known, and listed below. The plot also shows *P(k)* for a growing network with random attachment (Model A), that has an exponential *P(k)*^*27*^. **(d)** A lognormal distribution shown on a log-linear axis, along with its analytically calculated parameters. **(e)** The degree distribution of a scale-free network shown on a log-log plot, along with its analytically calculated parameters. **(f-h)** Compared to the exponential, Poisson and scale-free distributions, the lognormal form offers the best approximation for the degree distribution of the zebrafish **(f)**, the adult fly **(g)** and mouse **(h). (i)** The scaling of the maximum degree, *k*_*max*_ with *N* (brain size) for the three basic network models, a random network (green), a scale-free network with degree exponent γ= 2.5, and a lognormal network. Here, and in panel (j), the value γ = 2.5 for the degree exponent was chosen only for illustrative purposes, to demonstrate the fundamentally different classes scaling predictions (logarithmic, polynomial, and exponential) the different models offer. For comparison, we show as vertical lines the expected number of neurons *N* in the five brains explored here. **(j)** The scaling of the relative heterogeneity parameter κ with neuron count for the three basic network models.

To resolve these conflicting outcomes, here we compare data from eight connectomes spanning five organisms, arriving at the conclusion that the traditional models previously explored by network science are unable to account for the observed structural features of the connectome. Specifically, we show that the degree and the strength distributions of the connectome are inconsistent with both random^24^ and scale-free^23^ models, but both quantities are well approximated by a lognormal distribution. However, while lognormal distributions typically arise from multiplicative processes, networks are generally built through incremental node addition—an additive mechanism. Here we show that the lognormal architecture of the brain’s connectome can be derived once we recognize the physical nature of neurons: unlike social or technological networks, where nodes and links are distinct entities (Fig. 1a), the brain is composed of spatially extended neurites that simultaneously act as both nodes and links (Fig. 1b). We find that the large-scale architecture of the connectome is driven primarily by the length of its individual neurons^25,26^, which determine both a neuron’s synapse number (node strength) and its degree. We then use a Galton-Watson branching process to analytically derive the neuronal length distribution, not only predicting a lognormal degree and strength distributions, but obtaining falsifiable predictions that can account for the empirical data across all eight connectomes, along with the unique features of individual neural classes. The observed multiplicative network constitutes a previously unrecognized architectural class in network science^27–29^, blending characteristics of both random and scale-free topologies and reshaping canonical notions of robustness and synchronization.

## Datasets

We integrated data from eight connectome mapping efforts that offer synaptic connectivity at the resolution of single neurons for five organisms (SI 1): (1) the complete *C. elegans* connectome^1,30^ (*C. elegans*), (2) a neural-level map of the entire brain of the *Drosophila melanogaster* larva’s first instar phase^21^ (Fly Larva), and two efforts to map the adult *Drosophila*’s brain, (3) one reconstructing a single hemisphere (Hemi-Brain)^31^ and (4) another capturing neurons from both hemispheres^2–4^ (FlyWire), (5) a map of the *Drosophila* adult’s ventral nerve cord^32^ (MANC), (6) the brainstem of the larval zebrafish^33^ (Zebrafish), (7) a millimeter-scale volume of mouse primary visual cortex spanning from layer 1 to white matter^34,35^ (Mouse), and (8) a human surgical sample from the temporal lobe of the cerebral cortex^36^ (Human). While the size of the profiled brains spans eight orders of magnitude, from 302 neurons in the roundworm to an estimated 10^10^ human neurons^37^, most of the available connectomes represent incomplete samples: although they capture all 302 neurons in *C. elegans*^*1,30*^ and all 129,278 neurons in the adult Fly brain^2–4^, they map only 2,587 neurons of an estimated total of 100,000 in the larval zebrafish^33^, 52,059 out of 70 million neurons in the mouse brain^35^, and 46,378 out of 86 billion neurons in the human brain^36^ (Supplemental Table 1).

We harmonize all eight connectome maps by correcting formatting differences and rescaling spatial measurements based on imaging resolution and sample thickness (SI 1), enabling direct comparisons of neuron size and connectivity across datasets. For incomplete connectomes we distinguish completely mapped neurons from those that are only partially mapped (SI 1.3-1.5). We limit our study to complete neurons, while utilizing all reported synapses that complete neurons make (SI 1).

### The Degree Distribution of the Connectome is Neither Random nor Scale-Free

The degree distribution, *P(k)*, representing the probability that a neuron synapses with *k* other neurons, is a fundamental network property^27–29^, as we must know (and control for) *P*(*k*) to interpret any higher-order features, from network robustness^5^, to community structure^38,39^ or motifs^40^. While the functional form of *P(k)* is determined by multiple diverse local processes^27,28^, real networks tend to belong to one of two broad classes (SI 2.10): (1) Networks with a bounded *P(k)*, such as the Poisson degree distribution observed for random networks generated by the Erdős-Rényi model^24^ or a simple exponential emerging in randomly growing networks (Fig. 1c). The exponentially decaying tail^27^ of such bounded networks limits the variance in node sizes and excludes hubs; (2) Networks observed in many real systems, from protein interactions to the WWW, are scale-free, with a *P(k)* whose tail is often approximated by a power law^23^ (Fig. 1e), supporting the presence of multiple high degree nodes, known as hubs.

To elucidate the connectome’s architecture, we began by measuring the degree distribution for each dataset. We find that each connectome is characterized by a unique *P(k)* distribution whose average (peak position) and width is dataset dependent (Fig. 2a). Yet, all reconstructions share a unimodal shape on a log-linear plot (*P(k)* vs *log k*), leading to the hypothesis that apart from dataset-dependent logarithmic average μ_*k*_ *= <log k>* and logarithmic width σ_*k*_ (standard deviation of *P(log k)*) (Fig. 2a), the degrees may be drawn from the same distribution. To test this hypothesis, we employed a standard procedure of statistical physics, plotting 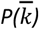, where 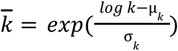, a linear transformation in the log space that corrects for differences in μ_*k*_ and σ_*k*_, but leaves the functional form of *P(k)* unchanged. We find that the rescaled degrees from each organism collapse onto a single 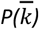 distribution (Fig. 2b, Supplemental Figure 16), supporting the hypothesis that, irrespective of the organism, the degrees are extracted from a single universal distribution (see Supplemental Figure 16b for statistical tests).

**Figure 2.**
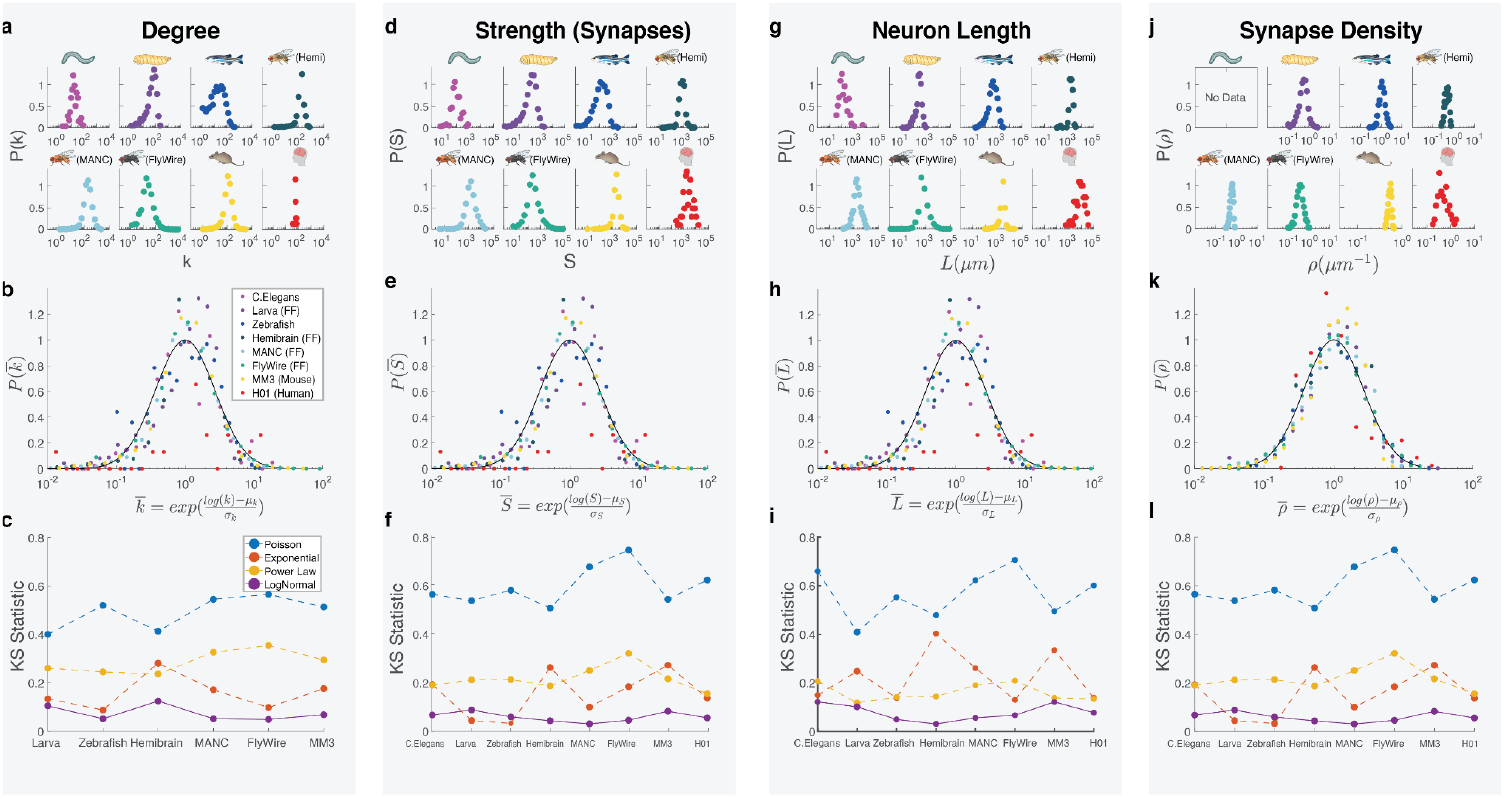
Neuron Degree and Strengths Follow Stereotypic Forms. **(a)** The degree distribution for the eight datasets shown on a log-linear plot. Each data point corresponds to the log-binned average value over multiple neurons (see SI 2.9). **(b)** The eight degree distributions in (a) plotted on the same figure after normalizing with their respective average (μ _*k*_) and variance (σ_*k*_). The fact that the distributions collapse into a single curve suggests that the values are sampled from the same distribution (see Supplemental Figure 16 for statistical validation). As a guide to the eye, we also show as a continuous line a lognormal distribution with μ=1 and σ=1. **(c)** The KS statistics for fitting the individual datasets shown in (a) to a Poisson, exponential, power law and lognormal distributions. We find that for each dataset, the lognormal has the lowest KS statistics, indicating that it offers the best fit to the data. **(d-f)** The strength distribution for the neurons, representing the total number of synapses each neuron has, for the eight datasets shown on a log-linear plot (d), along with their rescaled values (e), and KS parameters (f). **(g-i)** The log-binned neuron skeleton length *L* distribution for the eight datasets shown on a log-linear plot, along with their rescaled values (h) and KS parameters (i). **(j-l)** The log-binned local synapse density distribution for neurons (see SI 3.2 for definition) for the eight datasets shown on a log-linear plot, along with their rescaled values (k), and KS statistics (l). Curiously, even before rescaling, μ_ρ_ ∼ 1 across organisms, an observation consistent with previous comparative studies^55^. See SI 3.2 for the lacking information on *C. elegans* in panel j. Note that in panels **i** and **j** we fit a discrete Poisson to continuous variables *L* and ρ, justified given that for all studied datasets we have λ ≥ 20, hence the Poisson closely approximates a normal distribution, making the KS test appropriate.

The connectome is best described as a weighted network, as we can assign to each link between two neurons *i* and *j* the number of distinct synapses *s*_*ij*_ between them. In weighted networks, each node *i* is characterized by the node strength, 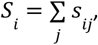, representing the total number of synapses neuron *i* has. We find *P(S)* to be again characterized by a symmetric and unimodal peak in a log-linear scale with different average and width (Fig. 2d). Plotting 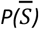, where 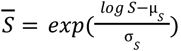, we again find that the rescaled distributions collapse onto a common 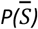 (Fig. 2e, Supplemental Figure 16).

Taken together, the existence of a common 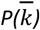 and 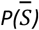 is unexpected, considering that the eight connectome datasets encompass five different species with varying levels of mapping completeness, diverse cell types, distinct synaptic connections, and a range of neurotransmitters. On one hand, our findings suggest the existence of stereotypical developmental mechanisms that shape the connectome’s structural features. However, the observed *P(k)* and *P(S)* are incompatible with the canonical degree distributions predicted by the existing network science models (Fig. 1f,h). Indeed, the empirically observed *P(k)* cannot be fit by a Poisson distribution (Erdős-Rényi model), a simple exponential *e*^*-αk*^ (randomly growing network model, Figure 1c), or by a power law *P*(*k*)∼*k*^−γ^ (scale-free network) (Fig. 1e). Kolmogorov-Smirnoff tests (Fig. 2c,f) indicate that all these canonical distributions have large KS statistics (see SI 2.5-2.8 for additional statistical measures). Instead, the clear unimodal and symmetric nature of the data in a log-linear plot (Fig. 2b,e) suggests that both *P(k)* and *P(S)* follow a lognormal distribution. Consistent with this hypothesis, we find that a fit to a lognormal has systematically low KS statistics across most datasets (Fig. 2c,f, see also SI 2.5-2.8 for additional statistical evidence). As we discuss in SI 2.5, for some datasets distributions like Weibull or Gamma also offer low KS values — however, we lack network-based generative models for these distributions.

While lognormal distributions provide reasonable fits to correlations in neural and brain activity^41^, as well as synapse sizes^42,43^, their significance to network structure remains less understood. The puzzle arises from how networks assemble: the incremental addition of nodes usually gives rise to either a power law or an exponential *P(k)*. While certain models of socio-economic systems allow for a lognormal *P(k)*, they require rather specific preferential attachment functions^44,45^, additional parameter families^46^ (SI 2.10), or narrowly defined fitness distributions as inputs^47–49^. In general, lognormal distributions naturally arise from multiplicative processes^50^ and yield exponential dynamics — fundamentally different from the incremental, node-by-node growth observed in real networks, including the brain. Consequently, the origin of lognormal distributions in network science, and the connectome in particular, remains unknown. At the same time, while the empirical evidence presented in Figs. 1-2 is evocative, it is insufficient to conclusively establish the lognormal nature of the connectome^51^. A definitive assessment also requires us to unveil the nature of the multiplicative process that can mechanistically explain the observed lognormal architecture.

### The Connectome is a Physical Network

The brain is often conceptualized as a hub-and-spoke network, where nodes represent the somas of each neuron, connected by directed and weighted links that represent the number of synaptic connections between dendrite and axon pairs (Fig. 1a). However, in the connectome we lack a separation between the nodes and the links, as links (synapses) require spatial proximity between the axons and the dendrites (Fig. 1b). In other words, the brain is a physical network^52^, hence its description demands a modeling framework that acknowledges the spatially extended physical nature of each neuron^25,26,53,54^. In physical networks, the longer a neuron is, the more chances it has to reach other neurons (proximity)^2,54,55^, and to synapse with them. Formally, this suggests that

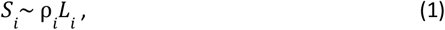

where *L*_*i*_ is the neuron’s total length and ρ is the *local synapse density*^*56*^ *of neuron i*, obtained by averaging the number of synapses encountered within 10 µms from any synapse (see SI 3.2). To test the validity of hypothesis (1), we plotted *S*_*i*_ vs *L*_*i*_ for each neuron (Fig. 3a), finding that while for small *L*_*i*_ we observe deviations from (1), in each of the eight datasets the linear scaling holds for large *L*_*i*_ neurons.

**Figure 3.**
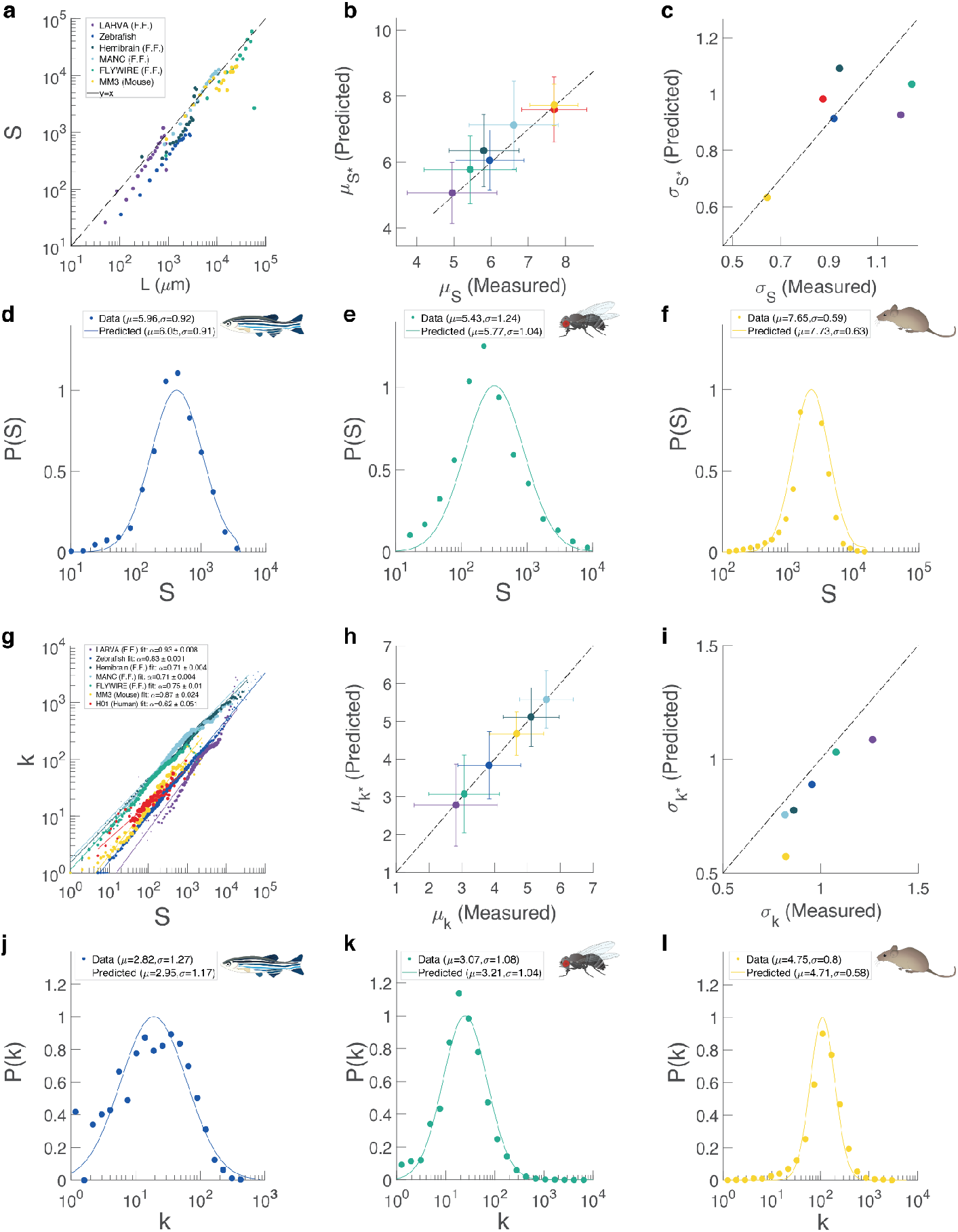
Predicting Dependencies between Neuron Lengths, Synapse Densities, and Degrees. **(a)** Scatter plot showing the strength S_i_ for each neuron *i* in function of its length *L*_*i*_, on a log-log plot. We show data for seven of the eight datasets in different colors, offering evidence for the scaling of Eq. (1). We excluded the human dataset as the low cell count left us an insufficient diversity for fitting *S* vs *k*. Each datapoint corresponds to log-binned average across multiple neurons. **(b)** A test of Eq. (3), that connects the empirically observed scale parameter μ_*S*_ for the neuron strength (horizontal axis) with the scale parameter predicted by Eq. 3 (vertical axis). **(c)** A direct test of Eq. (4), connecting the empirical observed shape parameter σ_*S*_ for the strength (horizontal axis) with the shape parameter predicted by Eq. 4 (vertical axis). **(d**,**e**,**f)** The observed strength distributions P(S) plotted along with a lognormal distribution with the analytically predicted (μ _*S*_, σ _*S*_) parameters. **(g)** Scatter plot showing the degree *k*_*i*_ for each neuron *i* in function of its length *S*_*i*_, on a log-log plot. We show data for each of the eight datasets in different colors, offering evidence for the sublinear scaling of Eq. (5). **(h)** A direct test of Eq. (6), that connects the empirically observed scale parameter μ_*k*_ for the neuron degrees (horizontal axis) with the scale parameter predicted by Eq. 6 (vertical axis). **(i)** A direct test of Eq. (7), connecting the empirical observed shape parameter σ_*k*_ for the neuronal (horizontal axis) with the shape parameter predicted by Eq. 7 (vertical axis). **(j**,**k**,**l)** The empirically observed degree distributions *P(k)* (symbols) plotted along with a lognormal distribution with the analytically predicted (μ_*k*_, σ_*k*_) parameters.

Given (1), a lognormal form for *P(S)* is only possible if the neuronal lengths also follow a lognormal distribution. This prompted us to measure the total length, *L*_*i*_, of each fully reconstructed neuron’s skeleton (Fig. 1b, SI 3.1), finding a unimodal and symmetric *P(L)* distribution on a log-linear plot (Fig. 2g). Plotting 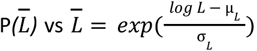, we find that each organism’s length distribution collapses onto the same functional form (Fig. 2h, Supplemental Figure 16).

Equation (1) also requires us to account for variations in synaptic densities, ρ_*i*._ We therefore determined the *local synapse density*, ρ_*i*_ of each fully reconstructed neuron, by limiting the measurement only to the regions of the neuron that contain synapses (a procedure that also avoids circular reasoning in the definition of ρ_*i*_, see SI 3.2). We find that *P(*ρ) for the different connectomes to be again unimodal (Fig. 2j) and under 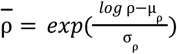 rescaling collapses onto a single 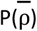 curve (Fig. 2k, Supplemental Figure 16). Finally, we fitted multiple candidate distributions to *P(L)* and *P*(ρ), finding that the lognormal form offers the lowest KS parameter across all datasets (Fig. 2i,l and SI 2.7 for additional statistical tests).

### Falsifying the Lognormal Connectome

Equation (1) helps us derive a series of falsifiable predictions that go beyond the scaling law of Fig. 3a. Indeed, let us assume that consistent with empirical data, *S* and *L* follow the lognormal distribution

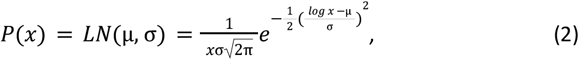

where μ = <*log*(*x*)> is the scale parameter, and σ > 0 is the shape parameter^57^, corresponding to the standard deviation of *P*(*log*(*x*)). However, if *L*_i_ and ρ_i_ both follow lognormal distributions with parameters *P*(*L*) = *LN*(μ_*L*_, σ_*L*_) and *P*(ρ) = *LN*(μ_ρ_, σ_ρ_), then their product (1) must also follow a lognormal distribution *P*(*S*) = *LN*(μ_*S*_, σ_*S*_), with modified parameters (SI 2.1),

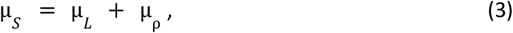

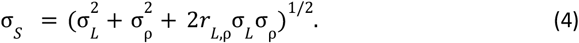

While formally (3) and (4) hold for any positive variables, under the additional assumption that *log L* and *log* ρ are approximately Gaussian, Eqs. (3)–(4) represent closure under multiplication, stating that *S* itself is lognormal with parameters (μ_S_,σ_S_) given by these identities (SI §2.2). To test the validity of (3) and (4), we measured the (μ, σ) parameters for each connectome (see SI Table 2-3), finding a remarkable agreement between the empirically observed (μ_*S*_,σ _*S*_), and the values predicted by (3) (Fig. 3b) and a reasonable agreement with (4) (Fig. 3c).

The final test of Eqs. (1)-(3) is offered by Fig. 3d-f, where we plot for three connectomes the empirically observed *P(S)* distribution (symbols), along with a lognormal function whose parameters (μ_*S*_, σ _*S*_) are predicted by Eqs. (3) and (4) (Fig. 3d-f, continuous line), finding that the predicted *P(S)* offers a good approximation for the empirically observed *P(S)* (Fig. 3d-f, symbols). Finally, we note that Eqs. (3)–(4) may be of practical relevance when data are incomplete. In microscopic studies, where accurate synapse counts are only partially available, neuronal lengths can often be measured with high accuracy and local synapse densities can be estimated. Combining these quantities through Eqs. (3)–(4) therefore enables quantitative predictions for both the expected total synapse number and its variability.

### Degree Distribution

As the number of synapses between neurons *i* and *j* is driven by the number of total synapses each neuron has (*S*_*i*_ and *S*_*j*_), it is tempting to write *k*_*i*_ ∼*S*_*i*_ ∼ρ _*i*_ *L*_*i*_ . However, as we have multiple synapses between two neurons (*s*_*ij*_ ≥1), we need to formalize the connectome as a weighted network, where *k*_*i*_ can have a nonlinear dependence on *S* _*i*_. Indeed, we find that the degrees are well approximated by

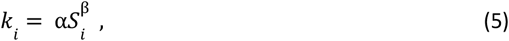

where β < 1 (Fig. 3g), a sublinear dependence frequently observed in weighted networks^58,59^. While (5) here represents an empirical observation, in Sect 4.6 we introduce a stochastic model that analytically predicts the observed nonlinear form. Yet, if *S* follows a lognormal distribution with parameters *P*(*S*) = *LN*(μ_*S*_, σ _*S*_), then *P(k)* must also follow a lognormal distribution with the modified parameters

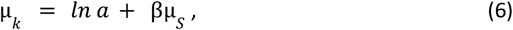

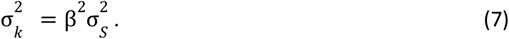

Once again, (6) and (7) represent closure under multiplication, that work with the evidence that *P(S)* and *P(L)* follow lognormals. To falsify the predictions (6) and (7), we plot in Fig. 3h,i the empirically observed (μ_*k*_, σ _*k*_) against the values predicted by (6) and (7), finding an agreement for most datasets. As a final test, we plot the empirically observed *P(k)* (symbols), together with the lognormal function whose parameters (μ_*k*_, σ_*k*_) are predicted by (7) and (8), confirming that the analytically predicted *P(k)* offers an excellent approximation to the empirically observed *P(k)* (Fig. 3j-l).

In summary, Figs. 2 and 3 indicate that *P(S), P(k), P(*ρ), and *P(L)* are stereotyped measures, displaying a remarkable consistency across the different connectomes. Further, Eqs. (3), (4), (6), and (7), establish relationships between the parameters of independently measured neural characteristics of *S, k*, ρ and *L*. So far we lack, however, a theoretical argument for the mechanistic origin of the lognormal distribution. Next we show that we can identify the origin of the observed distributions, if we focus on the origin of neuronal length distribution *P(L)*, given that according to Eqs. (1) and (5), *P(L)* determines both *P(S)* and *P(k)*.

### The Mechanistic Origin of the Lognormal Connectome

Neuronal growth is driven by complex genetically encoded developmental mechanisms^60^, guided by multiple families of signaling and adhesion molecules^61–65^. Yet, the stereotypical form of *P(L)* suggests that the mechanisms governing neuronal length are robust to genetic variability, hinting at the existence of a common mechanistic origin. Morphologically, every neuron is an incomplete tree (Fig. 1b and 4c, see also SI 4.1), and under rather general conditions the growth of such trees is formally described by a Galton-Watson (GW) branching process, introduced in 1874 to describe the extinction of social groups^66^ (SI 4.2) The process generates a random binary tree with *2*^*n*^ possible branches after *n* generations (Fig. 4a), with a total length of *L*_*n*_*=2*^*n*^*<ℓ>*, where *<ℓ>* is the average length of the individual branches. Although the growth of each branch is governed by cell-type-specific chemical signals, and partner feedback^67^, a mathematically robust description of neuronal branching can encapsulate these details into a layer *j* dependent survival probability 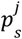 that represents the likelihood of a branches’ continued growth (Fig. 4a). In this case, the total length of a neuron after *n* generations follows (SI 4.4)

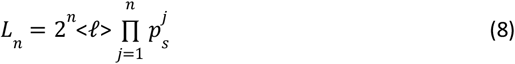

where we allow the survival probability to vary as the neuron grows, by assigning a different family of distributions 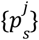, with parameter families 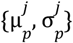 to each layer *j* (Fig. 4a). As (8) describes a multiplicative process (Fig. 4b), the multiplicative central limit theorem allows us to analytically derive the tree size distribution *P*(*L*_*n*_ /*ℓ*). Note that the standard GW dynamics asymptotically yields a Gaussian behavior, rather than the empirically observed lognormals. To make progress, we acknowledge the stochastic nature of the branching process, and treat 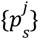 as stochastic variables, allowing 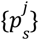 sets to vary from neuron to neuron, reflecting intrinsic developmental variability (SI 4.4). Now the variability compounds multiplicatively across generations, and, as we derive in SI 4.4, the resulting distribution becomes a lognormal (2) with parameters,

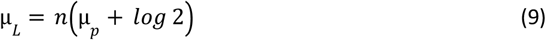

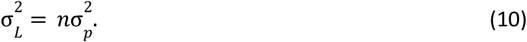

where *n* represents the layer number at which the tree tress stops expanding (i.e., when 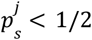 the numerical value of *n* will be determined later). In other words, the size distribution of the neural trees is determined by the survival probability distribution defined by 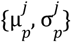, which govern the growth of the individual branches.

**Figure 4.**
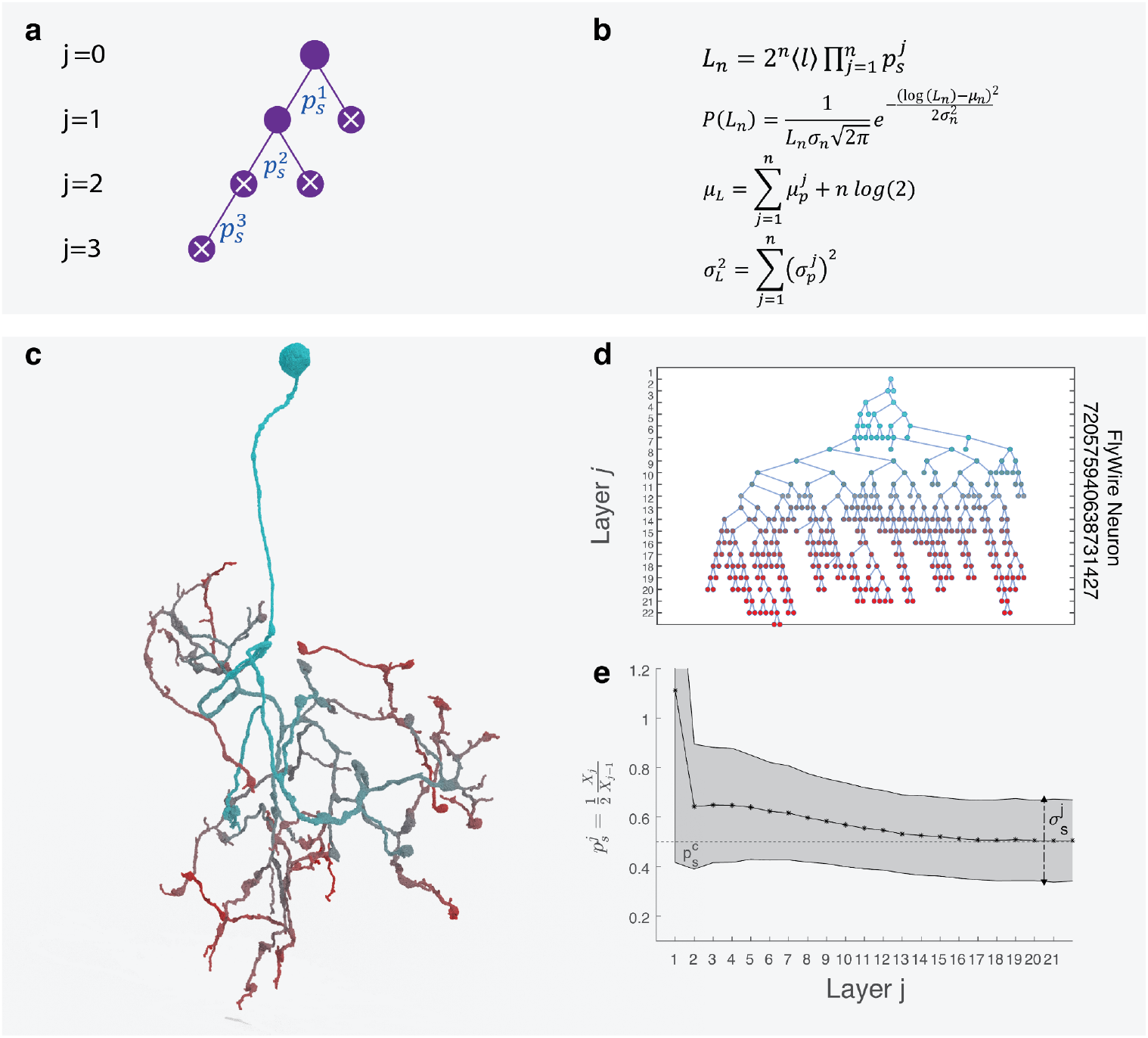
Multiplicative Process Governs Neuron Lengths and Predicts the Lognormal Architecture. **(a)** Each neuron is an incomplete tree whole growth we model via a Galton-Watson branching process. At each branching point at layer *j* the branch can either create two new nodes with survival probability 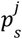, or cease permanently its growth, with probability 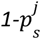. The figure illustrates this process for a small tree that survives for *n = j*_*max*_ *= 3* layers, creating a tree of depth *L* _3_. **(b)** The model predicts that a neuron’s total length *L*_*n*_ at any larger *n* follows a multiplicative process that, thanks to the central limit theorem, converges to a lognormal distribution whose parameters are governed by the layer dependent survival probability 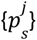, specifically the parameters 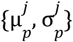. **(c)** The skeleton of FlyWire ALLN neuron 720575940638731427, each segment being colored according to its layer number *j* in the tree representation shown in panel d. **(d)** The incomplete tree behind the neuron shown in (c), extracted from its skeleton. **(e)** The empirically measured layer dependent average branching probability 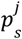 over all FlyWire neurons (black line) along with its variability across neurons (gray), shown in function of the layer number *j*. We find the average branching probability to be high near the soma (small *j*), but it rapidly decreases and stabilizes at a probability just above 0.5, i.e. always staying in the *p*_*s*_ *> 0*.*5* regime where the analytically derived lognormal distribution, shown in (b), is valid.

Predictions (9) and (10) are directly falsifiable, as for each neuron we have access to its detailed 3D reconstruction, allowing us to identify each individual branch from the spatial layout. We therfore measure the layer-dependent survival probability 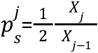, where *X*_*j*_ is the observed number of branches in layer *j* (Fig. 4c,d), allowing us to use maximum-likelihood estimation to extract 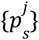 for each neuron *i* from the geometry of its skeleton. We performed this procedure for all FlyWire neurons, confirming that 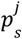 indeed depends on the layer number, being high near the soma (small *j*), but rapidly converging to 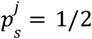 for large *j* (Fig. 4e). Ultimately Eqs. (8), (9) and (10) close the theoretical framework necessary to understand the neural-level architecture of the connectome, predicting that the multiplicative process governing the growth of the individual neuronal trees is responsible for the emergence of the lognormal *P(L)*. Further, in Supplemental Section 4.6 we introduced a stochastic process that treats the neuron length as a fitness variable that captures its ability to synapse with other neurons. This framework not only predicts the full stochastic lognormal form of P(S) and P(k) from P(L), but it also predicts that Eq. (5) must be always sublinear, in agreement with the empirical data of Fig 3g.

### Predicting the characteristic of individual neural classes

The more than 8,000 subtypes of neurons identified in the FlyWire dataset^3^ illustrate the exceptional diversity of individual neurons (SI 1.3), prompting us to ask if the theoretical framework developed above is capable of capturing these inherent morphological differences. Differences between classes are well described by each class’s unique *P(S)* and *P(L)* distributions, characterizing each of the 29 distinct neural classes identified for the adult fly^3^ (Fig. 5a,d, see also Supplemental Fig. 4 for *P(k)* and *P(*ρ)). First we verified that upon rescaling, the *P(S)* and *P(L)* distributions extracted for the different neural classes collapse into a single stereotyped distribution (Fig. 5b,e, Supplemental Figure 16), best approximated by a lognormal (Fig. 5c,f). At the same time, we find that each neuronal class is characterized by a different survival probability 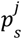 function, underpinning the different developmental processes governing them (Fig. 5g). To test the applicability of our theory to specific classes, we next used the empirically measured class-dependent survival probability functions 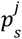 (Fig. 5g) as input to Eqs. (9) and (10), allowing us to predict the μ_*L*_ and σ _*L*_ parameters that govern the tree size distribution *P(L*/*ℓ)* for each neuron class. In Fig. 5h, we plot μ_*L*_ predicted by Eq. (9) against the empirically observed neuronal length μ _*L*_ = *< log* (*L*/*ℓ*) *>* for each of the 29 classes. The strong correlations between the two measurements show the validity of Eq (1). Further, the slope of the observed linear dependence allows us to empirically determine the characteristic layer number *n* in Eq. (8) at which p_s_ < 0.5, i.e. the neurons cease their expansion (but may still continue to grow branches for a few generations), finding to be *n* ≃ *20* for all classes (Fig. 5h). We also tested the validity of Eq (10), finding that the best fit provides n=22.31±3.68 (Fig, 5i), in agreement with Fig. 5h. Most importantly, the agreement between the predicted and measured μ_*L*_ = *< log* (*L*/*ℓ*) *>* not only confirms the predictive power of Eqs. (9) and (10), but also indicates that our theoretical framework, developed for the whole connectome, captures the characteristics of individual neuronal classes as well.

**Figure 5.**
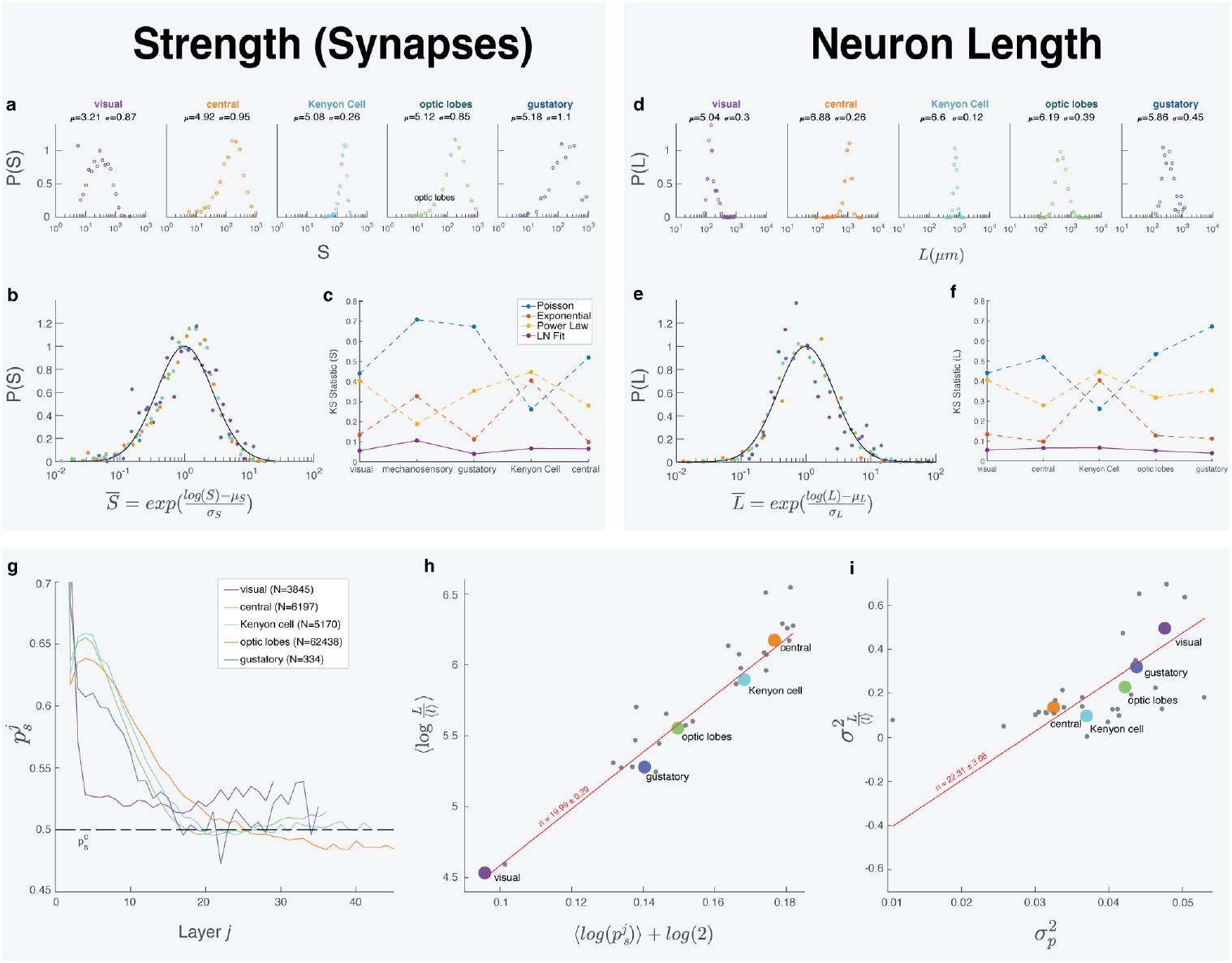
FlyWire cell classes follow lognormal distributions. To test whether the developed analytical framework applies to individual neuron classes, we focused on the 29 neuronal classes defined by the FlyWire project^3^. **(a)** Strength distribution for neurons from five FlyWire cell classes. The difference between these classes is captured by the different average synapse numbers, spanning from median *S = e*^*µ*^ = 24.8 synapses for visual neurons to median *S = e*^*µ*^ = 177.7 synapses for gustatory neurons. **(b)** The five distributions in (a) collapsed after normalizing with their respective μ_*S*_ and σ _*S*_ value, indicating that despite the significant difference in μ_*S*_ *= <log S>* and σ_*S*_, the values are stereotypical, being sampled from a single distribution. As a guide to the eye, we also show as a continuous line a lognormal distribution with μ=1 and σ=1. **(c)** KS statistics obtained by fitting the individual datasets shown in (a) to a Poisson, exponential, power law and log-normal distributions, indicating that for each neuronal class, the lognormal has the lowest KS statistics. **(d)** The neuron skeleton length distribution for the five representative FlyWire cell types shown in (a). The differences between these classes is illustrated by the significant differences in the average neuron lengths *L*∼ *e*^*µ*^, spanning from median *e*^*µ*^ = 144.5 microns for visual neurons to median *e*^*µ*^ *=* 972.6 microns for central neurons. **(e)** The five distributions in (e) after normalizing with their respective μ and σ value, along with a lognormal distribution with μ=1 and σ=1. **(f)** The KS statistics for fitting the individual datasets shown in (e) to a Poisson, exponential, power-law and log-normal distributions. For each dataset the lognormal has the lowest KS statistics. **(g)** The empirically measured branching probability 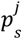 for five FlyWire cell classes versus the layer number *j*. Each cell type has its unique and different 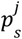 function, some approaching 0.5 faster, hence expecting a shorter layer number, and smaller neuronal length *L*. **(h)** Mean measured tree size *log* (*L*/*ℓ*) (vertical axis) vs the tree size predicted by Eq. (9) (horizontal axis), where we used as input to Eq. (9) the empirically measured survival function 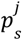 shown in (g). The symbols cover all 29 classes, highlighting the 5 classes shown in (a) and (e). Points are weighted linearly according to the relative frequency of each neuron class. The linear dependence between the predicted (horizontal axis) and the measured (vertical axis) tree size confirms that the neuron length *L* is primarily driven by survival branching probability 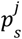, as predicted by Eqs. (8) and (9). The slope of the fit (*n*, indicated in red) reveals the characteristic layer number *n* ≃ 20 in Eq. (8) at which, on average, the neurons cease their branching expansion. This does not mean that the neuron stops growing completely at this depth, it simply means that in this regime p < 0.5, i.e. the individual branches are more likely to terminate than continue past this depth. **(i)** Standard deviation of measured tree size *log* (*L*/*ℓ*) (vertical axis) vs the tree size predicted by Eq. (10) (horizontal axis). While the σ prediction is noisier than that of the mean in (h), the slope is close to the n ≈ 20 observed for µ in (h), indicating that our theory is consistent between the two plots.

### Multiplicative Networks and Network Science

The observed lognormal degree distribution points to a network architecture in which degrees emerge through multiplicative rather than additive processes. Such *multiplicative networks* are previously unexplored in network science, and as we show next, they exhibit properties distinct from both random and scale-free networks. Consider, for example, the expected maximum degree, *k*_*max*_, which in a random network described by the Erdős-Rényi (ER) model, depends on the network size *N* as *log N*. Hence, for both *C. elegans* (*N* ≃ 300) and the human brain (*N* ≃ 10 ^11^), the *log N* dependence predicts the absence of hubs, captured by an implausibly modest maximum degree (Fig. 1i). By contrast, in a hypothetical scale-free brain the largest hub scales polynomially (e.g. 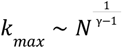), which can exceed *k* _*max*_ ∼10^7^ for humans, an unrealistically large number of distinct connections per neuron, predicting exceptionally large hubs. In a lognormal network, however, the maximum degree follows (Fig 1d,i)

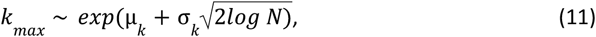

predicting a reasonable *k*_*max*_ ≃ 10 for *N* ≃ 10^11^. In other words, a multiplicative architecture can accommodate the existence of hub neurons while keeping their degree within biologically realistic limits, in line with previous empirical observations that hubs are less prominent in the fruit fly than in the worm^20^.

The novel features of multiplicative networks are reflected by the relative degree heterogeneity κ = *<k*^*2*^*>/<k>*, which governs many network-based processes, from robustness^6^ to resilience^7^ and synchronization^11,68,69^. In a random ER network κ = *<k> + 1*, independent of the network size. By contrast, for a scale-free network κ grows polynomially with N, following κ ∼ *N*^*(3*−*γ)/(γ*−*1)*^. In a lognormal network, however,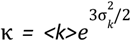(Figure 1d). Notably, κ does not depend on *N*, but its exponential dependence on σ_*k*_ can make it arbitrarily large. In other words, while a scale-free network’s robustness and synchronizability vary significantly with the network size, the corresponding properties of a multiplicative network depend only on changes in σ _*k*_, which depends on the developmental processes that shape the variability of degrees.

Finally, numerous empirical studies report a “rich-club” organization in brain networks—both in structural connectomes^20,70,71^ and fMRI-based functional maps^72,73^. We not only confirm these empirical results in our datasets, but also find that rich-clubs in the connectome are not ad hoc features but are direct consequences of the lognormal architecture. Specifically, the rich-clubs behavior is rooted in the sublinear degree–strength scaling (5), which, once inverted, predicts that 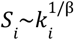, implying that high-degree nodes carry disproportionately large weights (see SI 4.8).

## Discussion and Conclusions

The connectome has long posed a quantitative challenge for network science, often yielding conflicting conclusions about its underlying architecture. Here, we reconcile these discrepancies by showing that the physical nature of neurons imposes features absent in previously studied non-physical social and technological networks. Accounting for this physicality enables us to analytically derive the empirically observed *P(k), P(S)*, and *P(L)* distributions, finding that each follows a lognormal distribution. Previous studies have noted the fat tailed nature of the weighted degree distribution P(S)^74–76^, also observing that the weighted degrees of source and sink neurons occasionally deviate from power laws, and are better fitted with a lognormal distribution^74,76^. Here, however, we show that lognormality is not an exception but the dominant feature — and we offer a directly testable mechanism for its emergence. Finally, we show that the lognormal connectome leads to a multiplicative network architecture, whose robustness and synchronizability are shaped not by the network’s size but by the intrinsic heterogeneity of neurons rooted in developmental processes.

We expect the theoretical framework introduced here to be enriched by the inclusion of additional developmental processes that govern neuron and synapse formation, leading to accurate generative models^27–29,77^. As an initial step in this direction, in SI 2.11 we discuss the Connectome Configuration Model, which tests our central hypothesis that the brain’s architecture is driven by the physical properties of individual neurons. While the model intentionally simplifies many aspects of neuronal structure and connectivity, it reproduces several key empirical observations, helping identify features that require more precise treatment of physicality.

We find that not only entire connectomes but also individual cell types are characterized by lognormal distributions, each defined by distinct, cell-line–dependent parameters. This observation suggests that future studies can exploit this variability—and its link to multiplicative branching-survival mechanisms—to identify the molecular determinants of neuronal growth, revealing how specific guidance cues (e.g., Netrins, Semaphorins, Ephrins) and their receptors influence branching probabilities in a cell-type–specific manner^61,62,78^. Although synapse formation requires spatial proximity between neurons, it also depends on the appropriate combination of membrane surface proteins to trigger synaptogenesis^79,80^. Accordingly, we need concurrent advances in understanding the packing of physical networks^19,81^, and the combinatorial expression of transcription factors^64,65^, to construct a biologically more detailed generative model.

Our study has several limitations that will likely be resolved by more complete future datasets. First, for mammalian brains, we currently have access to only a small fraction of the connectome—0.07% of estimated neurons for mice and 0.00005% for humans. While we mitigated this incompleteness by focusing on fully reconstructed neurons, the available maps systematically omit neurons whose size exceeds the mapped region.

To circumvent these data limitations, we examined length distributions obtained from injection-based tracing techniques (SI 1.5), confirming that all four tracer datasets follow a lognormal pattern. The elevated μ _*L*_ values observed suggest that current incomplete maps likely underestimate the (μ_*k*_, σ_*k*_) parameters. This data incompleteness also accounts for the observed log-skewness in *P(L), P(S)* and *P(k)* (SI 3.5). A further limitation arises from the incomplete annotation and reconstruction of axons in the zebrafish, mouse, and human datasets, raising the question of whether the lognormal architecture extends to them. To address this, we again turned to tracer studies and found that both axons and dendrites not only follow lognormal distributions but also display matching branching patterns (Supplementary Fig. 6). As larger vertebrate connectomics datasets become available, providing a more comprehensive view of axonal arborization, our growth models can be expanded to incorporate explicit axon–dendrite geometry and synapse directionality, thereby linking this statistical foundation to the extensive literature on spatial wiring optimization^82,83^ and compartment-specific growth.

Finally, understanding the connectome’s statistical properties is fundamental not only for elucidating its structural organization, but also for advancing models of brain dynamics. For instance, the slower scaling of the maximum degree in lognormal networks implies that loss of hubs may be less damaging compared to their known impact in scale-free networks. Further, the log-normal degree heterogeneity implies a finite spreading threshold and different conditions for global oscillations—features of direct relevance to seizure propagation and optogenetic stimulations. Lognormals may also provide an optimal architecture under competing design pressures: a log-normal *P(L)* distribution yields a small set of long, high-strength “backbone” neurons alongside a vast pool of short, hence may simultaneously be compatible with wiring economy^84^, developmental simplicity^85^, and computational capacity—echoing theories that lognormals in the functional properties of neurons support a fast-decision/slow-precision trade-off^41^. Understanding the role of optimization^82^, and whether further spatial or simplicity constraints are necessary, remains an important future direction, especially if we wish to understand how higher level design principles interact with the role of specific symmetries^86^ and cluster-synchronisation motifs^87^ that modulate function at the local circuit scale. Insights into these architectural principles are also crucial for linking genetic variations and developmental constraints to alterations in neural wiring that underlie specific brain disorders, like Fetal Alcohol Syndrome. This knowledge will foster a deeper integration of structural and functional neuroscience, with broad implications for both basic research and clinical practice.

## Data Availability

All datasets were acquired from public repositories. Instructions for downloading and cleaning data, as well as summary datasets for producing figures are available at https://github.com/bdanubius/ComparativeConnectomics. Intermediate files and larger datasets can be found at https://storage.googleapis.com/connectome-additional-data/Archive.zip.

## Code Availability

The code used for this manuscript is available at https://github.com/bdanubius/ComparativeConnectomics.

## Acknowledgements

D.L.B. is supported by funding from the Eric and Wendy Schmidt Center at the Broad Institute of MIT and Harvard. A.-L.B. is supported by the NSF award No 2243104 – COMPASS and by the European Union’s Horizon 2020 research and innovation program No 810115 – DYNASNET. We thank Daria Koshkina for designing figures.

## Author Contributions

B.P., D.L.B. and A.F.C. collected and cleaned the data. B.P. and D.L.B. analyzed the data. B.P. and G.M. conducted theoretical analyses. B.P., D.L.B., G.M. and A.-L.B. wrote, reviewed and edited the manuscript.

## Supplementary Information

Supplementary Information is available for this paper.

## References

1. White, J. G., Southgate, E., Thomson, J. N. & Brenner, S. The structure of the nervous system of the nematode Caenorhabditis elegans. Philos. Trans. R. Soc. Lond. B Biol. Sci. 314, 1–340 (1986).

2. Dorkenwald, S., Matsliah, A., Sterling, A. R., Schlegel, P., Yu, S.-C., McKellar, C. E., Lin, A., Costa, M., Eichler, K., Yin, Y., Silversmith, W., Schneider-Mizell, C., Jordan, C. S., Brittain, D., Halageri, A., Kuehner, K., Ogedengbe, O., Morey, R., Gager, J., Kruk, K., Perlman, E., Yang, R., Deutsch, D., Bland, D., Sorek, M., Lu, R., Macrina, T., Lee, K., Alexander Bae, J., Mu, S., Nehoran, B., Mitchell, E., Popovych, S., Wu, J., Jia, Z., Castro, M., Kemnitz, N., Ih, D., Bates, A. S., Eckstein, N., Funke, J., Collman, F., Bock, D. D., Jefferis, G. S. X., Sebastian Seung, H., Murthy, M. & the FlyWire Consortium. Neuronal wiring diagram of an adult brain. Nature 634, 124–138 (2024).

3. Schlegel, P., Yin, Y., Bates, A. S., Dorkenwald, S., Eichler, K., Brooks, P., Han, D. S., Gkantia, M., dos Santos, M., Munnelly, E. J., Badalamente, G., Capdevila, L. S., Sane, V. A., Pleijzier, M. W., Tamimi, I. F. M., Dunne, C. R., Salgarella, I., Javier, A., Fang, S., Perlman, E., Kazimiers, T., Jagannathan, S. R., Matsliah, A., Sterling, A. R., Yu, S.-C., McKellar, C. E., FlyWire Consortium, Costa, M., Sebastian Seung, H., Murthy, M., Hartenstein, V., Bock, D. D. & Jefferis, G. S. X. Whole-brain annotation and multi-connectome cell typing quantifies circuit stereotypy in Drosophila. Nature 634, 139–152 (2024).

4. Zheng, Z., Scott Lauritzen, J., Perlman, E., Robinson, C. G., Nichols, M., Milkie, D., Torrens, O., Price, J., Fisher, C. B., Sharifi, N., Calle-Schuler, S. A., Kmecova, L., Ali, I. J., Karsh, B., Trautman, E. T., Bogovic, J. A., Hanslovsky, P., Jefferis, G. S. X., Kazhdan, M., Khairy, K., Saalfeld, S., Fetter, R. D. & Bock, D. D. A Complete Electron Microscopy Volume of the Brain of Adult Drosophila melanogaster. Cell 174, 730–743.e22 (2018).

5. Albert, R., Jeong, H. & Barabási, A. L. Error and attack tolerance of complex networks. Nature 406, 378–382 (2000).

6. Cohen, R., Erez, K., ben-Avraham, D. & Havlin, S. Resilience of the internet to random breakdowns. Phys Rev Lett 85, 4626–4628 (2000).

7. Gao, J., Barzel, B. & Barabási, A.-L. Universal resilience patterns in complex networks. Nature 530, 307–312 (2016).

8. Pastor-Satorras, R. & Vespignani, A. Epidemic spreading in scale-free networks. Phys Rev Lett 86, 3200–3203 (2001).

9. Liu, Y.-Y. & Barabási, A.-L. Control principles of complex systems. Rev. Mod. Phys. 88, (2016).

10. Gu, S., Pasqualetti, F., Cieslak, M., Telesford, Q. K., Yu, A. B., Kahn, A. E., Medaglia, J. D., Vettel, J. M., Miller, M. B., Grafton, S. T. & Bassett, D. S. Controllability of structural brain networks. Nat Commun 6, 8414 (2015).

11. Nishikawa, T., Motter, A. E., Lai, Y.-C. & Hoppensteadt, F. C. Heterogeneity in oscillator networks: are smaller worlds easier to synchronize? Phys. Rev. Lett. 91, 014101 (2003).

12. Barabási, D. L., Bianconi, G., Bullmore, E., Burgess, M., Chung, S., Eliassi-Rad, T., George, D., Kovács, I. A., Makse, H., Nichols, T. E., Papadimitriou, C., Sporns, O., Stachenfeld, K., Toroczkai, Z., Towlson, E. K., Zador, A. M., Zeng, H., Barabási, A.-L., Bernard, A. & Buzsáki, G. Neuroscience Needs Network Science. J Neurosci 43, 5989–5995 (2023).

13. Amaral, L. A., Scala, A., Barthelemy, M. & Stanley, H. E. Classes of small-world networks. Proc. Natl. Acad. Sci. U. S. A. 97, 11149–11152 (2000).

14. Sporns, O. & Zwi, J. D. The small world of the cerebral cortex. Neuroinformatics 2, 145–162 (2004).

15. Ed Bullmore, Anna Barnes, Danielle S Bassett, Alex Fornito, Manfred Kitzbichler, David Meunier, John Suckling. Generic aspects of complexity in brain imaging data and other biological systems. Neuroimage 47, 1125–1134 (2009).

16. van den Heuvel, M. P., Stam, C. J., Boersma, M. & Hulshoff Pol, H. E. Small-world and scale-free organization of voxel-based resting-state functional connectivity in the human brain. Neuroimage 43, 528–539 (2008).

17. Eguíluz, V. M., Chialvo, D. R., Cecchi, G. A., Baliki, M. & Apkarian, A. V. Scale-free brain functional networks. Phys Rev Lett 94, 018102 (2005).

18. Yang, R., Vishwanathan, A., Wu, J., Kemnitz, N., Ih, D., Turner, N., Lee, K., Tartavull, I., Silversmith, W. M., Jordan, C. S., David, C., Bland, D., Sterling, A., Goldman, M. S., Aksay, E. R. F., Seung, H. S. & Eyewirers. Cyclic structure with cellular precision in a vertebrate sensorimotor neural circuit. Curr Biol 33, 2340–2349.e3 (2023).

19. Salova, A. & Kovács, I. A. Combined topological and spatial constraints are required to capture the structure of neural connectomes. Network Neuroscience (2025). At <http://arxiv.org/abs/2405.06110>

20. Lin, A., Yang, R., Dorkenwald, S., Matsliah, A., Sterling, A. R., Schlegel, P., Yu, S.-C., McKellar, C. E., Costa, M., Eichler, K., Bates, A. S., Eckstein, N., Funke, J., Jefferis, G. S. X. E. & Murthy, M. Network statistics of the whole-brain connectome of Drosophila. Nature 634, 153–165 (2024).

21. Winding, M., Pedigo, B. D., Barnes, C. L., Patsolic, H. G., Park, Y., Kazimiers, T., Fushiki, A., Andrade, I. V., Khandelwal, A., Valdes-Aleman, J., Li, F., Randel, N., Barsotti, E., Correia, A., Fetter, R. D., Hartenstein, V., Priebe, C. E., Vogelstein, J. T., Cardona, A. & Zlatic, M. The connectome of an insect brain. Science 379, eadd9330 (2023).

22. Betzel, R., Puxeddu, M. G. & Seguin, C. Hierarchical communities in the larval Drosophila connectome: Links to cellular annotations and network topology. bioRxiv (2023). doi:10.1101/2023.10.25.562730

23. Barabási, A. L. & Albert, R. Emergence of scaling in random networks. Science 286, 509–512 (1999).

24. Erdős, P. & Rényi, A. On the evolution of random graphs. Publications of the Mathematical Institute of the Hungarian Academy of Sciences 5, 17–61 (1960).

25. Horvát, S., Gămănuț, R., Ercsey-Ravasz, M., Magrou, L., Gămănuț, B., Van Essen, D. C., Burkhalter, A., Knoblauch, K., Toroczkai, Z. & Kennedy, H. Spatial Embedding and Wiring Cost Constrain the Functional Layout of the Cortical Network of Rodents and Primates. PLOS Biology 14, e1002512 (2016).

26. Ercsey-Ravasz, M., Markov, N. T., Lamy, C., Van Essen, D. C., Knoblauch, K., Toroczkai, Z. & Kennedy, H. A predictive network model of cerebral cortical connectivity based on a distance rule. Neuron 80, 184–197 (2013).

27. Barabási, A.-L. Network Science. (Cambridge University Press, 2016).

28. Dorogovtsev, S. N. & Mendes, J. F. F. The Nature of Complex Networks. (2025).

29. Caldarelli, G. Scale-Free Networks: Complex Webs in Nature and Technology. (OUP Oxford, 2007).

30. Cook, S. J., Jarrell, T. A., Brittin, C. A., Wang, Y., Bloniarz, A. E., Yakovlev, M. A., Nguyen, K. C. Q., Tang, L. T.-H., Bayer, E. A., Duerr, J. S., Bülow, H. E., Hobert, O., Hall, D. H. & Emmons, S. W. Whole-animal connectomes of both Caenorhabditis elegans sexes. Nature 571, 63–71 (2019).

31. Scheffer, L. K., Xu, C. S., Januszewski, M., Lu, Z., Takemura, S.-Y., Hayworth, K. J., Huang, G. B., Shinomiya, K., Maitlin-Shepard, J., Berg, S., Clements, J., Hubbard, P. M., Katz, W. T., Umayam, L., Zhao, T., Ackerman, D., Blakely, T., Bogovic, J., Dolafi, T., Kainmueller, D., Kawase, T., Khairy, K. A., Leavitt, L., Li, P. H., Lindsey, L., Neubarth, N., Olbris, D. J., Otsuna, H., Trautman, E. T., Ito, M., Bates, A. S., Goldammer, J., Wolff, T., Svirskas, R., Schlegel, P., Neace, E., Knecht, C. J., Alvarado, C. X., Bailey, D. A., Ballinger, S., Borycz, J. A., Canino, B. S., Cheatham, N., Cook, M., Dreher, M., Duclos, O., Eubanks, B., Fairbanks, K., Finley, S., Forknall, N., Francis, A., Hopkins, G. P., Joyce, E. M., Kim, S., Kirk, N. A., Kovalyak, J., Lauchie, S. A., Lohff, A., Maldonado, C., Manley, E. A., McLin, S., Mooney, C., Ndama, M., Ogundeyi, O., Okeoma, N., Ordish, C., Padilla, N., Patrick, C. M., Paterson, T., Phillips, E. E., Phillips, E. M., Rampally, N., Ribeiro, C., Robertson, M. K., Rymer, J. T., Ryan, S. M., Sammons, M., Scott, A. K., Scott, A. L., Shinomiya, A., Smith, C., Smith, K., Smith, N. L., Sobeski, M. A., Suleiman, A., Swift, J., Takemura, S., Talebi, I., Tarnogorska, D., Tenshaw, E., Tokhi, T., Walsh, J. J., Yang, T., Horne, J. A., Li, F., Parekh, R., Rivlin, P. K., Jayaraman, V., Costa, M., Jefferis, G. S., Ito, K., Saalfeld, S., George, R., Meinertzhagen, I. A., Rubin, G. M., Hess, H. F., Jain, V. & Plaza, S. M. A connectome and analysis of the adult central brain. Elife 9, (2020).

32. Azevedo, A., Lesser, E., Phelps, J. S., Mark, B., Elabbady, L., Kuroda, S., Sustar, A., Moussa, A., Khandelwal, A., Dallmann, C. J., Agrawal, S., Lee, S.-Y. J., Pratt, B., Cook, A., Skutt-Kakaria, K., Gerhard, S., Lu, R., Kemnitz, N., Lee, K., Halageri, A., Castro, M., Ih, D., Gager, J., Tammam, M., Dorkenwald, S., Collman, F., Schneider-Mizell, C., Brittain, D., Jordan, C. S., Dickinson, M., Pacureanu, A., Seung, H. S., Macrina, T., Lee, W.-C. A. & Tuthill, J. C. Connectomic reconstruction of a female Drosophila ventral nerve cord. Nature 631, 360–368 (2024).

33. Vishwanathan, A., Ramirez, A. D., Wu, J., Sood, A., Yang, R., Kemnitz, N., Ih, D., Turner, N., Lee, K., Tartavull, I., Silversmith, W. M., Jordan, C. S., David, C., Bland, D., Goldman, M. S., Aksay, E. R. F., Seung, H. S. & the Eyewirers. Predicting modular functions and neural coding of behavior from a synaptic wiring diagram. bioRxiv (2020). doi:10.1101/2020.10.28.359620

34. MICrONS Consortium, Bae, J. A., Baptiste, M., Bodor, A. L., Brittain, D., Buchanan, J., Bumbarger, D. J., Castro, M. A., Celii, B., Cobos, E., Collman, F., da Costa, N. M., Dorkenwald, S., Elabbady, L., Fahey, P. G., Fliss, T., Froudarakis, E., Gager, J., Gamlin, C., Halageri, A., Hebditch, J., Jia, Z., Jordan, C., Kapner, D., Kemnitz, N., Kinn, S., Koolman, S., Kuehner, K., Lee, K., Li, K., Lu, R., Macrina, T., Mahalingam, G., McReynolds, S., Miranda, E., Mitchell, E., Mondal, S. S., Moore, M., Mu, S., Muhammad, T., Nehoran, B., Ogedengbe, O., Papadopoulos, C., Papadopoulos, S., Patel, S., Pitkow, X., Popovych, S., Ramos, A., Reid, R. C., Reimer, J., Schneider-Mizell, C. M., Seung, H. S., Silverman, B., Silversmith, W., Sterling, A., Sinz, F. H., Smith, C. L., Suckow, S., Takeno, M., Tan, Z. H., Tolias, A. S., Torres, R., Turner, N. L., Walker, E. Y., Wang, T., Williams, G., Williams, S., Willie, K., Willie, R., Wong, W., Wu, J., Xu, C., Yang, R., Yatsenko, D., Ye, F., Yin, W. & Yu, S.-C. Functional connectomics spanning multiple areas of mouse visual cortex. bioRxiv (2021). doi:10.1101/2021.07.28.454025

35. Schneider-Mizell, C. M., Bodor, A. L., Brittain, D., Buchanan, J., Bumbarger, D. J., Elabbady, L., Gamlin, C., Kapner, D., Kinn, S., Mahalingam, G., Seshamani, S., Suckow, S., Takeno, M., Torres, R., Yin, W., Dorkenwald, S., Bae, J. A., Castro, M. A., Halageri, A., Jia, Z., Jordan, C., Kemnitz, N., Lee, K., Li, K., Lu, R., Macrina, T., Mitchell, E., Mondal, S. S., Mu, S., Nehoran, B., Popovych, S., Silversmith, W., Turner, N. L., Wong, W., Wu, J., MICrONS Consortium, Reimer, J., Tolias, A. S., Seung, H. S., Reid, R. C., Collman, F. & Maçarico da Costa, N. Cell-type-specific inhibitory circuitry from a connectomic census of mouse visual cortex. bioRxiv (2024). doi:10.1101/2023.01.23.525290

36. Shapson-Coe, A., Januszewski, M., Berger, D. R., Pope, A., Wu, Y., Blakely, T., Schalek, R. L., Li, P. H., Wang, S., Maitin-Shepard, J., Karlupia, N., Dorkenwald, S., Sjostedt, E., Leavitt, L., Lee, D., Troidl, J., Collman, F., Bailey, L., Fitzmaurice, A., Kar, R., Field, B., Wu, H., Wagner-Carena, J., Aley, D., Lau, J., Lin, Z., Wei, D., Pfister, H., Peleg, A., Jain, V. & Lichtman, J. W. A petavoxel fragment of human cerebral cortex reconstructed at nanoscale resolution. Science 384, eadk4858 (2024).

37. Herculano-Houzel, S. & Lent, R. Isotropic fractionator: a simple, rapid method for the quantification of total cell and neuron numbers in the brain. J. Neurosci. 25, 2518–2521 (2005).

38. Newman, M. E. J. & Girvan, M. Finding and evaluating community structure in networks. Phys Rev E 69, 026113 (2004).

39. Fortunato, S. Community detection in graphs. Phys. Rep. 486, 75–174 (2010).

40. Milo, R., Shen-Orr, S., Itzkovitz, S., Kashtan, N., Chklovskii, D. & Alon, U. Network motifs: simple building blocks of complex networks. Science 298, 824–827 (2002).

41. Buzsáki, G. & Mizuseki, K. The log-dynamic brain: how skewed distributions affect network operations. Nat. Rev. Neurosci. 15, 264–278 (2014).

42. Loewenstein, Y., Kuras, A. & Rumpel, S. Multiplicative dynamics underlie the emergence of the log-normal distribution of spine sizes in the neocortex in vivo. J Neurosci 31, 9481–9488 (2011).

43. Song, S., Sjöström, P. J., Reigl, M., Nelson, S. & Chklovskii, D. B. Highly Nonrandom Features of Synaptic Connectivity in Local Cortical Circuits. PLOS Biology 3, e68 (2005).

44. Sheridan, P. & Onodera, T. A Preferential Attachment Paradox: How Preferential Attachment Combines with Growth to Produce Networks with Log-normal In-degree Distributions. Sci Rep 8, 2811 (2018).

45. Redner, S. Citation statistics from 110 years of physical review. Phys. Today 58, 49–54 (2005).

46. Kim, M. & Leskovec, J. Multiplicative Attribute Graph Model of Real-World Networks. Internet Mathematics 8, 113–160 (2010).

47. Buzsáki, G. The Brain from Inside Out. (Oxford University Press, 2019).

48. Liu, Y., Seguin, C., Betzel, R. F., Han, D., Akarca, D., Di Biase, M. A. & Zalesky, A. A generative model of the connectome with dynamic axon growth. Netw Neurosci 8, 1192–1211 (2024).

49. Smith, K. M. Explaining the emergence of complex networks through log-normal fitness in a Euclidean node similarity space. Sci Rep 11, 1976 (2021).

50. Mitzenmacher, M. A brief history of generative models for power law and lognormal distributions. Internet Math. 1, 226–251 (2004).

51. Serafino, M., Cimini, G., Maritan, A., Rinaldo, A., Suweis, S., Banavar, J. R. & Caldarelli, G. True scale-free networks hidden by finite size effects. Proc Natl Acad Sci U S A 118, (2021).

52. Dehmamy, N., Milanlouei, S. & Barabási, A.-L. A structural transition in physical networks. Nature 563, 676–680 (2018).

53. Pósfai, M., Szegedy, B., Bačić, I., Blagojević, L., Abért, M., Kertész, J., Lovász, L. & Barabási, A.-L. Impact of physicality on network structure. Nature Physics (2023). doi:10.1038/s41567-023-02267-1

54. Pete, G., Timár, Á., Stefánsson, S. Ö., Bonamassa, I. & Pósfai, M. Physical networks as network-of-networks. Nat Commun 15, 4882 (2024).

55. Ferreira Castro, A. & Cardona, A. Invariant synaptic density links neuronal activity stability and wiring optimisation principles across species. bioRxiv (2024). doi:10.1101/2024.07.18.604056

56. Wittner, L., Henze, D. A., Záborszky, L. & Buzsáki, G. Three-dimensional reconstruction of the axon arbor of a CA3 pyramidal cell recorded and filled in vivo. Brain Struct Funct 212, 75–83 (2007).

57. Menichetti, G. & Barabási, A.-L. Nutrient concentrations in food display universal behaviour. Nature Food 3, 375–382 (2022).

58. Goh, K. I., Kahng, B. & Kim, D. Universal behavior of load distribution in scale-free networks. Phys Rev Lett 87, 278701 (2001).

59. Barrat, A., Barthélemy, M., Pastor-Satorras, R. & Vespignani, A. The architecture of complex weighted networks. Proc Natl Acad Sci U S A 101, 3747–3752 (2004).

60. Fishell, G. & Kepecs, A. Interneuron Types as Attractors and Controllers. Annu. Rev. Neurosci. 43, 1–30 (2020).

61. Sanes, J. R. & Zipursky, S. L. Synaptic Specificity, Recognition Molecules, and Assembly of Neural Circuits. Cell 181, 536–556 (2020).

62. Kerstjens, S., Michel, G. & Douglas, R. J. Constructive connectomics: How neuronal axons get from here to there using gene-expression maps derived from their family trees. PLoS Comput. Biol. 18, e1010382 (2022).

63. Flaherty, E. & Maniatis, T. The role of clustered protocadherins in neurodevelopment and neuropsychiatric diseases. Curr. Opin. Genet. Dev. 65, 144–150 (2020).

64. Barabási, D. L. & Barabási, A.-L. A Genetic Model of the Connectome. Neuron 105, 435–445.e5 (2020).

65. Kovács, I. A., Barabási, D. L. & Barabási, A.-L. Uncovering the genetic blueprint of the nervous system. Proc. Natl. Acad. Sci. U. S. A. 117, 33570–33577 (2020).

66. Harris, T. E. The Theory of Branching Processes. (Springer, 1963).

67. Matsumoto, N., Barson, D., Liang, L. & Crair, M. C. Hebbian instruction of axonal connectivity by endogenous correlated spontaneous activity. Science 385, eadh7814 (2024).

68. Restrepo, J. G., Ott, E. & Hunt, B. R. Onset of synchronization in large networks of coupled oscillators. Phys Rev E 71, 036151 (2005).

69. Lee, D.-S. Synchronization transition in scale-free networks: clusters of synchrony. Phys Rev E Stat Nonlin Soft Matter Phys 72, 026208 (2005).

70. Towlson, E. K., Vértes, P. E., Ahnert, S. E., Schafer, W. R. & Bullmore, E. T. The rich club of the C. elegans neuronal connectome. J Neurosci 33, 6380–6387 (2013).

71. Betzel, R., Puxeddu, M. G. & Seguin, C. Hierarchical communities in the larval connectome: Links to cellular annotations and network topology. Proc Natl Acad Sci U S A 121, e2320177121 (2024).

72. Bassett, D. S. & Bullmore, E. Small-world brain networks. Neuroscientist 12, 512–523 (2006).

73. Li, Q., Del Ferraro, G., Pasquini, L., Peck, K. K., Makse, H. A. & Holodny, A. I. Core language brain network for fMRI language task used in clinical applications. Netw Neurosci 4, 134–154 (2020).

74. Cirunay, M. T., Batac, R. C. & Ódor, G. Learning and criticality in a self-organizing model of connectome growth. Sci Rep 15, 31890 (2025).

75. Lynn, C. W., Holmes, C. M. & Palmer, S. E. Heavy-tailed neuronal connectivity arises from Hebbian self-organization. Nat. Phys. 20, 484–491 (2024).

76. Cirunay, M., Ódor, G., Papp, I. & Deco, G. Scale-free behavior of weight distributions of connectomes. Phys. Rev. Res. 7, (2025).

77. Bianconi, G. Higher Order Networks: An Introduction to Simplicial Complexes. (Cambridge University Press, 2021).

78. Agi, E., Reifenstein, E. T., Wit, C., Schneider, T., Kauer, M., Kehribar, M., Kulkarni, A., von Kleist, M. & Hiesinger, P. R. Axonal self-sorting without target guidance in visual map formation. Science 383, 1084–1092 (2024).

79. Südhof, T. C. Towards an Understanding of Synapse Formation. Neuron 100, 276–293 (2018).

80. Valdes-Aleman, J., Fetter, R. D., Sales, E. C., Heckman, E. L., Venkatasubramanian, L., Doe, C. Q., Landgraf, M., Cardona, A. & Zlatic, M. Comparative Connectomics Reveals How Partner Identity, Location, and Activity Specify Synaptic Connectivity in Drosophila. Neuron 109, 105–122.e7 (2021).

81. Zhang, X.-Y., Moore, J. M., Ru, X. & Yan, G. Geometric Scaling Law in Real Neuronal Networks. Phys Rev Lett 133, 138401 (2024).

82. Meng, X., Piazza, B., Both, C., Barzel, B. & Barabási, A.-L. Surface Optimisation Governs the Local Design of Physical Networks. (2025). at <http://arxiv.org/abs/2509.23431>

83. Chklovskii, D. B., Schikorski, T. & Stevens, C. F. Wiring optimization in cortical circuits. Neuron 34, 341–347 (2002).

84. Chen, B. L., Hall, D. H. & Chklovskii, D. B. Wiring optimization can relate neuronal structure and function. Proc Natl Acad Sci U S A 103, 4723–4728 (2006).

85. Takagi, K. Energy constraints on brain network formation. Sci Rep 11, 11745 (2021).

86. Morone, F. & Makse, H. A. Symmetry group factorization reveals the structure-function relation in the neural connectome of Caenorhabditis elegans. Nature Communications 10, 1–13 (2019).

87. Avila, B., Augusto, P., Zimmer, M., Serafino, M. & Makse, H. A. Fibration symmetries and cluster synchronization in the Caenorhabditis elegans connectome. ArXiv arXiv:2305.19367v2 (2024). at <https://pmc.ncbi.nlm.nih.gov/articles/PMC10312817/>

