## SupplementalInformation for "Physical Network Constraints Define the Multiplicative Architecture of the Brain’s Connectome"

### Section 1: Data Collection and Standardization

In order to facilitate a comparative analysis of neuronal structure and connectivity, we began our study by compiling connectome data from multiple species. All datasets originate from Electron Microscopy (EM) imaging of neural tissue, in which the authors of cited studies utilized algorithmic and human annotation methods to reconstruct cell morphologies and synaptic connections. We limited our analyses to recent large-scale and highly complete datasets from *Caenorhabditis elegans* (nematode*), Drosophila melanogaster* (fruit fly), *Danio rerio* (zebrafish), *Mus musculus* (mouse), and human.

The datasets originated from different groups, which resulted in differences in the utilized microscope, imaging slice size, reconstruction algorithms, and proofreading benchmarks. Nevertheless, all groups extracted the full morphology of neurons by segmenting EM images to trace neuronal processes, involving automated algorithms and manual curation to accurately capture the complex branching structures of axons and dendrites. Next, the authors of cited studies extracted neuron skeletons from these detailed reconstructions to create simplified representations that retain essential features such as neurite diameter, branching topology, and spatial location. For all datasets, we downloaded all neuron skeleton morphology files (.swc format, where available), along with the synaptic connections of each cell. Given the differences in samples and collection methods, we began our analysis by addressing variations in spatial scaling and removing partially reconstructed neurons.

We first corrected for differences in the formatting of released datasets. During sample collection, researchers made independent decisions related to imaging sample size and resolution, which influences the x and y coordinates of the data, as well as the sample thickness, corresponding to the distance in z between samples. To address this, in cases where spatial measurements were not defined in microns (the Fly Larva and Human were released in nm, while the Hemibrain and MANC were shared in arbitrary units), we rescaled the units to microns based on the imaging resolution and z-thickness utilized in the source paper. This standardization enabled direct comparisons of neuron size and connectivity across datasets.

Next, we focused our analysis on high-quality neurons, whose definition varies between datasets, but generally corresponds to fully mapped and human-proofread neurons. The *C. elegans*, larval fly, and Flywire adult fly connectomes captured all cells in the brain of the animal. Thus, in our analysis we utilized all neurons reported in the reconstruction. The other five connectomes captured local regions of an animal's nervous system. In order to focus our analysis on complete cells, that do not exit the region mapped by electron microscope, we developed an "edge filter," that identified all neurons that were close (within 5% according to the volume's dimension) to the boundary of the reconstruction. We applied this edge filter to the mouse, as well as for neurons marked "proofread" in the adult fly hemibrain and adult fly ventral nerve cord (MANC) datasets. For the human connectome, we utilized the 104 published proofread cells but did not include further edge filtering, as the volume was comparable in size to a neuron's skeleton. We applied no filtering for the larval zebrafish dataset, as again, the reconstructed volume was comparable to neuronal size, thus, most, if not all, cells would have been lost. We report all our results in the manuscript based on the final list of high-quality cells in each dataset. However, for synapse number and degree, we include in our analysis connections from high-quality cells to low-quality cells. This means that we did not use solely the subgraph of high-quality cells, but rather included in our statistics both high-quality to high-quality connections ("core") as well as high-quality to low-quality neuron connections ("periphery"). We next discuss each dataset in more detail.

#### Section 1.1: *C. elegans*

The roundworm *C. elegans* represents the first collected whole-organism connectome[1]. Since this initial milestone, which was achieved using hand reconstructions of AEI 6B and AEI 802 electron microscope imaging, groups have created more modern maps of the worm nervous system through reanalysis of these original micrographs, as well as the collection of new connectomes. Here, we use the OpenWorm's data release of the hermaphrodite *C. elegans* adult[2], which contains the skeletons of neurons from a 2011 reanalysis[3] of the original 1986 connectome[1], as well as an associated list of synapses between neurons. The *C. elegans* hermaphrodite contains 302 cells, however two neurons, CANL and CANR, do not make synaptic connections in this reconstruction[3], thus we exclude them from our analysis, leaving 300 cells with 20,905 synapses between them. We note that in many *C. elegans* connectomes the synapse number is used interchangeably with synapse size, if the synapse shows up on multiple EM sections. The OpenWorm data release we downloaded aimed to correct for this fact, counting such "multi-synapses" only as a single synapse, although we find that both formulations are consistent with a lognormal distribution (Fig. S1).

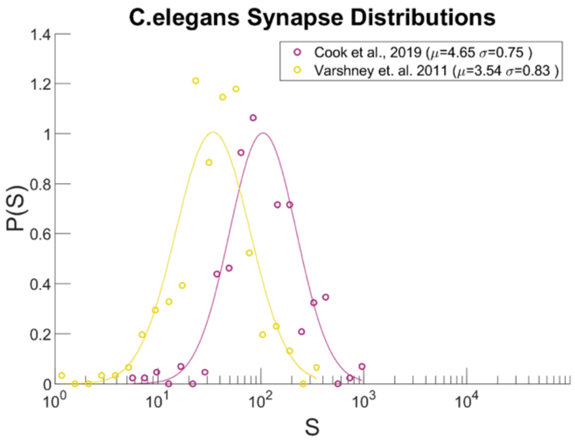

**Supplemental Figure 1: The impact of alternative definitions of synapse number in *C. elegans*.** Connectomics in *C. elegans* has previously utilized synapse size interchangeably with synaptic strength. More specifically, connections between neurons are counted as the total number of EM sections on which synapses occur between the two cells, meaning that each physical synapse may be counted multiple times. The Varshney et al[3] dataset we rely our analysis counts each synapse individually, whereas the Cook et al[4] dataset utilizes the multi-counting method. We find that both formulations of synapse number in *C. elegans* are consistent with a lognormal distribution, with shifted log average $\mu$.

#### Section 1.2: Larval Drosophila melanogaster

The 6-hour-old female *Drosophila melanogaster* 1st instar larva connectome represented the largest full-brain map at the time of publication[5], consisting of over 3,000 neurons at 3.8 × 3.8 × 50 nm resolution. These neurons span the full central nervous system and a significant portion of the ventral nervous system of the animal, where all released neurons were considered at least 75% complete in terms of their synaptic and morphological reconstructions, with many cells reaching 99% completion. We downloaded 3,066 neuron morphologies, of which 29 had an improperly formatted file structure, leaving 3,037 neurons for our analysis. These neurons make 599k synapses with each other, or with other cells in the volume. We note that these cells were released with measurements in nanometers, which we converted to micrometers.

#### Section 1.3: Adult Drosophila melanogaster

The entire body of the fly contains approximately 140,000 neurons in the central nervous systems and an additional 23,000 neurons in the Ventral Nerve Cord. Multiple teams have worked to reconstruct the fly's nervous system, and we describe each dataset we utilized in further detail below. However, since these teams shared resources, joint metrics exist for validating the accuracy of the reconstruction. One such metric is the "% completeness," for which all three datasets average around 50%, with the Hemi-Brain ranging between 20 and 85%, depending on the region. While these numbers seem concerningly low, the authors of the Hemi-Brain dataset highlight that this percentage mainly reflects the impracticality of tracing all fine branches of every cell in a fly brain. Further, the authors showed that the traced cells, which we utilized, provide a representative sample of neuronal circuits in these datasets, as the missing connections are independently distributed. Specifically, in a test where the study authors reconstructed a 30% complete region to 50% completeness, almost all connections that changed had more synapses, very few connections got fewer synapses, and no new strong (many synapse) connections appeared. Thus, the neuron length and synapse number may be lower than the "true" full reconstruction (which Hemi-Brain authors term "beyond reach and largely superfluous"), however given that these omissions are independent, the relative comparisons between these quantities, i.e. the distributions we analyzed in this manuscript, should remain consistent. We note that similar reconstruction completeness issues will be present for all datasets (which we discuss in upcoming section), except for the *C. elegans*, thus comparing distributions across species remains valid.

Hemi-Brain

The fly Hemi-Brain[6] represents the first major reconstruction of the fruit fly central nervous system, and, true to its name, covers roughly half of a female fly's central brain, including the mushroom body and central complex (circuits critical for associative learning and fly navigation). This 250 x 250 x 250 $\mu m^3$ dataset was acquired with a FIB-SEM at 8x8x8 $nm^3$ resolution. We downloaded 22,699 proofread neurons, which we found to be in arbitrary units corresponding to the voxel size, leading us to rescale by 125x along each dimension in order to arrive to a μm description of the data.

Again, as the name of dataset suggests, the Hemi-Brain connectome contained only a partial snapshot of the full central nervous system of this animal, and thus many cells had branches that extended out of the volume. In order to account for this, we restricted our analysis to neurons whose bodies were not cut off by the boundaries of the reconstructed volume, which we defined as not entering the outer 5% of the volume along any dimension. Given this filtering, we were left with 3,898 complete, proofread neurons (tagged "roughly traced" or "traced") as well as the 3.2 million synapses they make with each other, or with other cells in the volume. In Fig. S2, we compared the quantities we explored in the paper (distributions of neuron length, synapse number and degree) before (black) and after (blue) filtering out neurons close to the edge of the dataset. Note that the filtering resulted in the removal of the low-quality, incomplete (and therefore small) neurons at the left of the distribution. Note that while the filtering changes the $(\mu,\sigma)$ values of the lognormal, it does not alter the fact that the lognormal offers the best fit to the data.

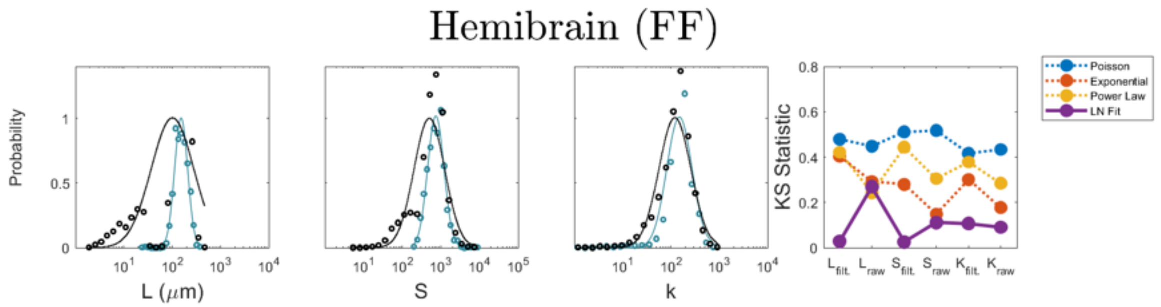

**Supplemental Figure 2: Effect of filtering on fly hemibrain measurements.** Distributions of neuron length, synapse number and degree before (black) and after (blue) filtering out neurons close to the edge of the dataset, which we considered most likely to be incomplete. KS tests show that lognormal fits increase significantly with filtering for S and L. Further, prior to filtering the L was best fit by a power law, and prior to filtering both S and L had comparable fits to lognormals with the exponential distribution.

Ventral Nerve Cord (MANC)

With about 23,000 neurons, 10 million pre-synaptic sites, and 74 million post-synaptic densities, the Male Adult Nerve Cord (MANC)[7] connectome is a densely reconstructed map of synaptic connections in the fruit fly nerve cord – a structure analogous to the human spinal cord that controls most of the fly's motor functions. At the time of release, it was the first complete nerve cord connectome and the first connectome of a bilaterally complete region of the central nervous system of an adult animal. The VNC sample is >500 µm long and, in places, as large as 250 µm diameter, and was imaged using a FIB-SEM at 8x8x8nm resolution, thus we again rescaled by 125x along each dimension in order to arrive to a µm description of the data.

Finally, as the full projection of cells out of the VNC, into the brain and out into the body, was not reconstructed, we again limited ourselves to cells that did not enter the outer 5% of the volume along any dimension. Given this filtering, we were left with 12,379 complete, proofread neurons (tagged "prelim roughly traced") as well as the 27.5 million synapses they make with each other, or with other cells in the volume. We again compared the distributions of neuron length, synapse number and degree before (black) and after (blue) filtering (Fig. S3), which resulted in the removal of low-quality, incomplete (and therefore small) neurons at the left of the distribution. Note that while the filtering changes the ($\mu$,$\sigma$) values of the lognormal fit, it does not alter the fact that the lognormal offers the best fit to the data.

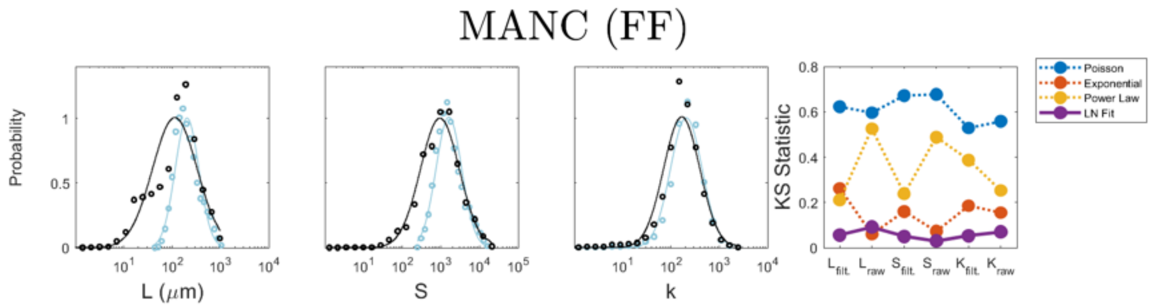

**Supplemental Figure 3: Effect of filtering on fly ventral nerve cord (MANC) measurements.** Distributions of neuron length, synapse number and degree before (black) and after (blue) filtering out neurons close to the edge of the dataset, which we considered most likely to be incomplete. KS tests show that lognormal fits improve with filtering for S and k. Further, prior to filtering the L was best fit by an exponential, and S had comparable fits to lognormals with the exponential distribution.

Full Brain (FlyWire)

Collecting the full wiring diagram of the fly nervous system represents one of the largest connectomics successes to date. This 750x369x286 $\mu m^3$ sample was imaged with a serial section TEM (ssTEM) at 4x4x40nm$^3$ resolution (286μm = 7,062 slices x 40nm)[8]. We downloaded the v630 release of this dataset with 129,278 neurons and 33 million synapses between them[9]. These cells were not only proofread, but annotations of cell classes and types, nerves, hemilineages and predictions of neurotransmitter identities were available for all neurons. Nearly all cells were fully encompassed in the volume (estimated 85%[9]), with the exception of a small fraction of cells that projected out of the brain into the VNC or body, however without a clear tag for these cells, we chose to utilize every neuron with a cell body (soma) in the brain. We admit that this will include a handful of partial cells, however with the large number of neurons reconstructed in this dataset, we believe this did not significantly influence our results (see SI 3.5 for relevant experiments).

The Neuronal Classes of Adult fly

A major achievement of the adult fly connectomics efforts has been the extensive cell type identification performed, with 8,453 unique subtypes annotated at the time of the FlyWire release[9]. The annotation hierarchy is produced at four levels: 3 flows, subdivided into 9 superclasses, which break down into 29 classes, and finally 8,435 cell types, which provide salient labels at different granularities[10] (Fig. S4, top). The first two levels, flow and superclass, were densely annotated: every neuron is either afferent, efferent or intrinsic to the brain (flow) and falls into one of the nine superclasses: sensory (periphery to brain), motor (brain to periphery), endocrine (brain to corpora allata/cardiaca), ascending (ventral nerve cord (VNC) to brain), descending (brain to VNC), visual projection (optic lobes to central brain), visual centrifugal (central brain to optic lobes), or intrinsic to the optic lobes or the central brain.

The 29 classes further subdivide the superclasses and are based on pre-existing neurobiological groupings from the literature. We utilize the 29 class-level grouping in the main text as it includes already specific cell types (Kenyon Cells), but still groups together broader regions (visual, gustatory, central, optic lobes). In this way, we arrive to a cell type level for which we can interpretability analyze *P(S)* (Fig. 5a), *P(L)* (Fig. 5e), *P(k)* (Fig. S4, middle), and *P(ρ)* (Fig. S4, bottom), testing the applicability of our theoretical framework to individual classes of neurons.

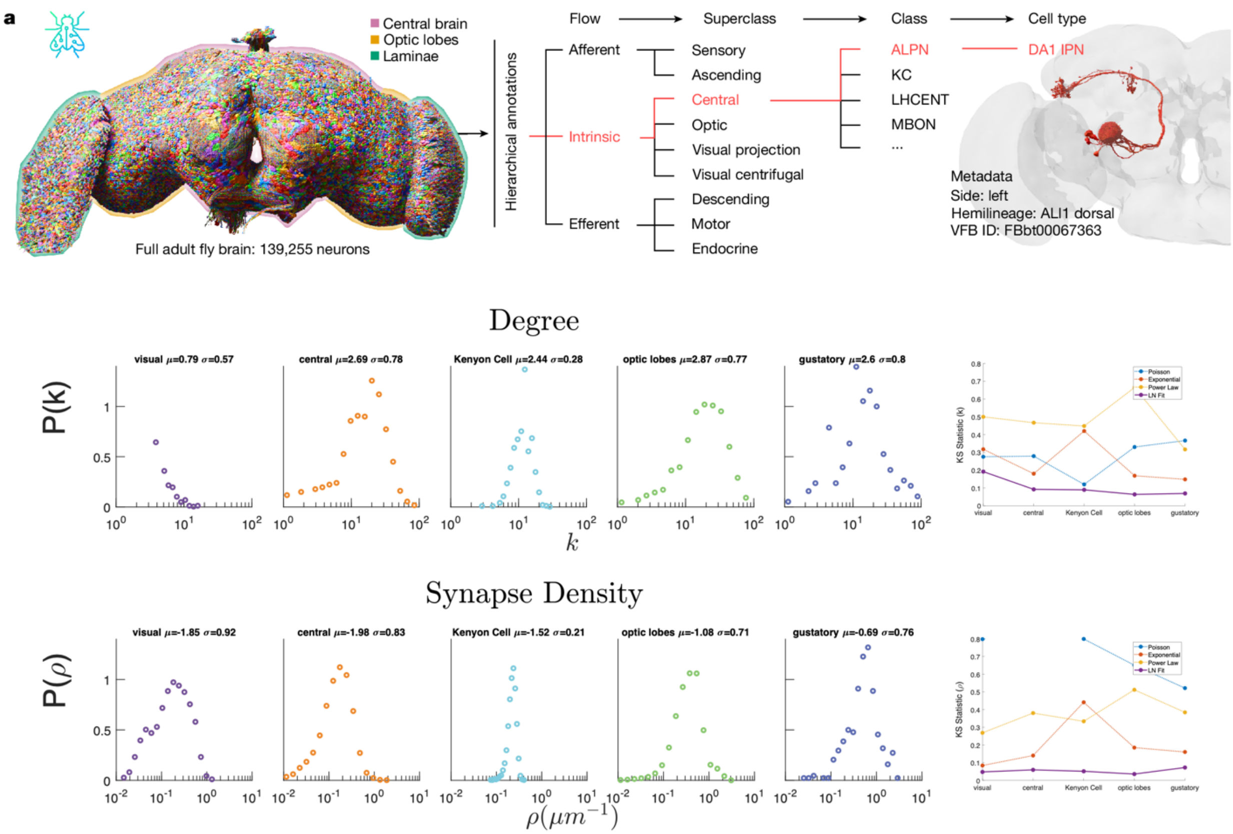

**Supplemental Figure 4: Degree and Synapse Density follow a lognormal distribution at the cell class level in FlyWire. Top:** The hierarchical classification scheme of adult fly cell types, from flow, to superclass, class and cell type. From Schlegel et al. (2024)[10]. **Middle:** The degree distribution for the five cell classes of Fig. 5 shown on a log-linear plot. On the right, we show the KS statistics for fitting the individual cell classes to a Poisson, Exponential, power law and lognormal distributions. We find that for each class, the lognormal has the lowest KS statistics, indicating that it offers the best fit to the data. **Bottom:** The synapse density distribution for the neurons, representing the total number of synapses each neuron has, for the five cell classes shown on a log-linear plot. On the right, we show the KS statistics for fitting the cell classes to a Poisson, Exponential, power law and log-normal distributions. We find that for each class, the lognormal has the lowest KS statistics, indicating that it offers the best fit to the data.

#### Section 1.4: Zebrafish

Multiple long-term efforts to map the full 7-day larval zebrafish nervous system, consisting of an approximate 100,000 neurons, are under way, yet the most complete released dataset maps only a portion of the fish's hindbrain[11,12]. This reconstructions represents a local snapshot of 220 μm by 112 μm by 57 μm, imaged by serial Electron Microscopy at a resolution of 5 × 5 × 45 nm and contained a total of 2,967 neuronal cell bodies with well-known landmarks such as the Mauthner neuron, the axon of the contralateral Mauthner neuron, neurons MiD2 and MiD3 of the reticulo-spinal network, and a number of commissural bundles. We note that this presents a small, but representative, volume for this animal, however most, if not all, cells are not fully contained by the imaged region. Specifically, filtering in a similar manner to the HemiBrain, MANC, or Mouse datasets, where we only include cells that do not enter the outer 5% of the volume, would only leave 189 "complete" cells. For this reason, we utilized all proofread cells and associated synapses released in the publication for our analysis, without any filtering, which corresponded to 2,587 neurons and 1.39 million synapses.

#### Section 1.5: Mouse

Mapping the full connectome of the approximately 400 cubic millimeters and 70 million neurons of the mouse brain is under way, however thus far the largest released reconstruction covers a 1.023 $mm^3$ of mouse visual cortical areas[13]. This dataset, collected as part of the IARPA MICrONS program, spans 1.4mm x .87mm x .84 mm, consisting of two merged transmission electron microscopy volumes spanning all cortical layers (pia to white matter) in an adult male mouse. The scale of the imaged region was sufficient to capture the entire dendritic arbor of typical cortical neurons, with proofreading of excitatory neurons aiming to reconstruct complete dendritic arbors, while proofreading of inhibitory neurons reconstructed both complete dendritic arbors and extensive but incomplete axonal arbors. The limitations of the region size means that the study authors reconstructed all incoming connections onto excitatory cells, although the long-range outgoing connections (axons) and morphology of excitatory neurons were largely absent, while for inhibitory cells it was possible to capture nearly the entire morphology, with all incoming connections as well as a significant portion of outgoing connections and morphology.

The study authors' automated detection methods estimated that 120,000 neurons, and more than 523 million synapses, exist within the imaged data. Of these cells, 52,059 have been reconstructed and proofread in v661, which we downloaded. We limited our analysis to the most complete proofread neurons in the reconstructed volume, which we defined as not entering the outer 5% of the volume along any dimension. Through this filtering, we focused our study to a set of 22,350 neurons, which retained diversity in layer depth and cell type labels, as well as the 135 million synapses they made with each other, or with other cells in the volume. In Fig. S5, we compared the distributions of neuron length, synapse number and degree before (black) and after (blue) filtering, although less of an effect was observed in this dataset than due to previous filtering efforts on the fly dataset, likely due to the large number of cells reconstructed. Note that while the filtering changes the ($\mu$,$\sigma$) values of the lognormal fit, it does not alter the fact that the lognormal offers the best fit to the data.

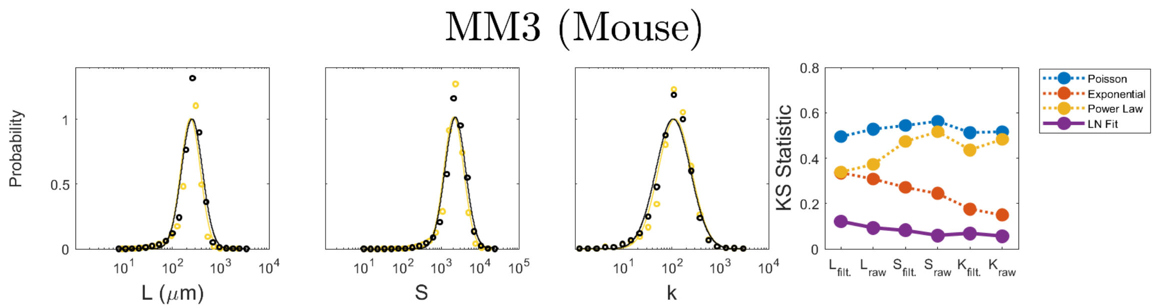

**Supplemental Figure 5: The input of filtering the mouse connectome.** Distributions of neuron length *P(L)*, synapse number *P(S)* and degree *P(k)* before (black) and after (yellow) filtering out incomplete neurons close to the edge of the dataset.

Long-Range Projections in the Mouse Brain

A major limitation of the mouse and human (Section 1.6) connectomics efforts lies in the limited volume reconstructed, when compared to the full size of the brain. Given that the long-range projections of many cells are truncated, a concern remains that the distributions we observe at the local level in the mouse and human connectome will not scale to larger reconstructions, where full cells are available. In order to address this, we analyzed the length distributions of neurons from injection-based techniques, which provide the opportunity to map the full projection structure of cells, albeit at a lower resolution and without synaptic connectivity information. These datasets had on the order of thousands of reconstructed cells, with either injections all over the brain (DEN-SEU[14], 10,860 cells (b); Peng et al[15], 1,741 cells (c); Liu et al[16], 1,876 cells (d)) or in specific regions, like the Prefrontal Cortex[17] (6,357 cells, e). This way, we gained access to a more diverse pool of neurons in mouse from multiple studies, including different cortical regions and subcortical structures. We find that the dendritic reconstructions (DEN-SEU) have a distribution similar to the local regions profiled in MM3, while datasets that contain larger neurons have a corresponding higher $\mu$ than the EM mouse connectome. Crucially, they all exhibit a lognormal distribution of neuron lengths (Fig. S6, top). While larger neurons, with more synapses and higher degrees, will be found in larger EM reconstructions, this concordance between EM and tracing datasets suggests that the lognormal shape will be confirmed by future datasets, allowing us to correct $\mu$ to higher values.

As axonic traces are limited and truncated in the EM connectome, we relied on light microscopy datasets to analyze axons and dendrites independently. To do so, we separated the dendrites and axons out of the full neurons in the Peng et al[15] and Liu et al[16] reconstructions, and added on the dendrites of DEN-SEU and axons of Prefrontal Cortex. We find that when considered separately, the length distributions of axons and dendrites individually follow a lognormal. Yet, the analysis also revealed systematic differences between the two classes: we find $\mu_{axon}$ = 10.53–11.46 (depending on the dataset), considerably larger than the $\mu$dendrite range of 7.66–8.96. Note that $\mu$ is the logarithm of the distance, hence these values indicate a roughly 5–45 fold difference in the length of dendrites and axons, differences that align with biological expectations, as axons are typically longer than dendrites. Note that the difference may be further amplified by the fact that many studies emphasize neurons with long-range axonal projections, while dendritic morphology is often less resolved than in EM datasets.

We also examined the branching patterns of these neurons, measuring the parameters predicted by the stochastic GW model to define neuronal characteristics (Section S4). On one hand, we confirm that both axons and dendrites exhibit a declining $p_s$ with layer number, in line with what we previously reported across all neurons and specific neuronal classes (Figs 4-5 in the manuscript). While at first these curves are largely indistinguishable across datasets, with do find a key difference between axons and dendirtes: the layer at which the $p_s$ crosses the critical value of 0.5 is 4-5 for dendrites and it higher, at 7-8 layers for axons. This difference in $p_s$ helps explain the observed length differences (and hence the different $\mu_{dendrite}$ and $\mu_{axon}$ values). Yet, the similar shapes of the curve suggests that the underlying generative processes governing the growth of axons and dendrites may be quite similar.

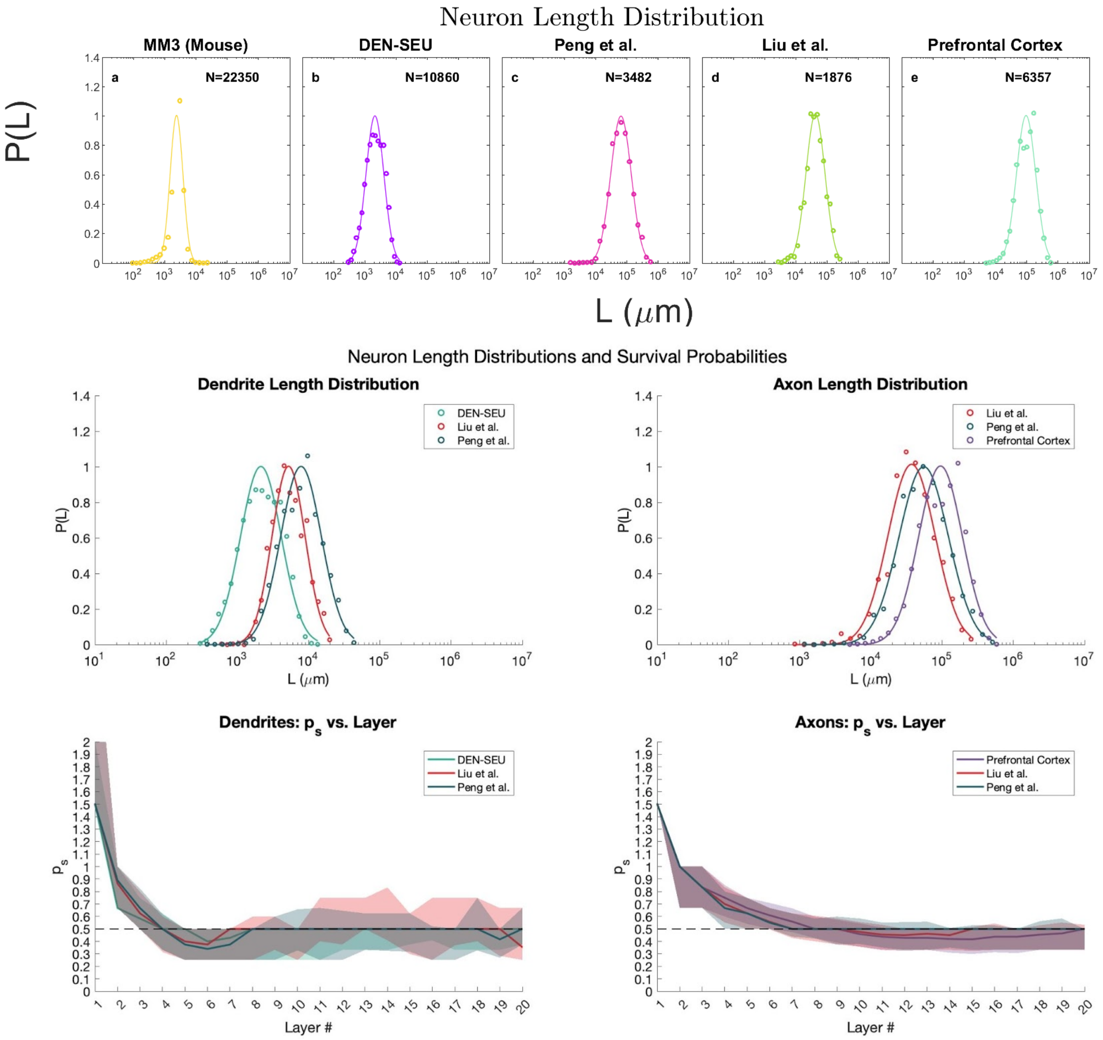

**Supplemental Figure 6: Incomplete and complete mouse neurons follow lognormal length distributions.**

**Top:** We compare the neuron length distributions of the mouse EM connectome (MM3) with those of dendritic reconstructions (DEN-SEU[14], 10,860 cells (b)) and entire cells reconstructed from tracer injections brain-wide (Peng et al[15], 1,741 cells (c); Liu et al[16], 1,876 cells (d)), as well in specific regions, like the Prefrontal Cortex[17] (6,357 cells, (e)). All neurons, either high quality, but local, as in the EM dataset, or lower quality but brain-wide (b-e) are well fit by a lognormal distribution. **Middle:** The length distributions of dendrites (left) and axons (right), fit by lognormals (KS test statistics 0.0147-0.0524), with similar average lengths across the diverse reconstructions. In this plot, points are the log bin values, and the curve is the bit fit lognormal line. **Bottom:** The branching probability $p_s$ vs layer number, as originally observed in FlyWire (Figures 4-5 in the manuscript), we find that the dendrites (bottom, left) and axon (bottom, right) each have a characteristic branching profile, that is similar across datasets. Solid lines show the median, while the error bands visualize the 25 and 75 percentile distributions of the data.

#### Section 1.6: Human

The full human brain contains approximately 86 billion neurons in 1.3 million cubic millimeters, which stands orders of magnitudes away from the capabilities of current electron microscopy techniques. In a recently published study, a representative sample of human temporal cortex, 1.05 $mm^3$ in volume, was obtained during surgery of a 45-year old female epileptic patient[18]. The study authors cut the sample into 5,019 sections at an average thickness of 33.9 nm) for a total sample thickness of 0.170 mm, and then imaged using a multibeam scanning EM at $4 \times 4$ $nm^2$ resolution. The resulting dataset contains over 45,000 neuronal cell bodies, and 130 million synaptic connections, spanning all cortical layers. However, many of the cells extend out of the volume, thus represent only fragments of the full cell, or are incompletely reconstructed. For this reason, we restricted our analysis to the 104 high quality proofread neurons released by the authors of the study, as well as all the synapses they make with any reconstructed (complete or incomplete) cells in the volume. We note that the reconstruction by the study authors were highly biased towards connections made onto the 104 proofread neurons, assumedly because the dendritic structure of these cells were well contained by the volume, but the axonal arborage, which would contain the connections from these cells onto other neurons, likely rapidly exited the volume, and thus was not available for analysis. This fact means that the degree and synapse number of the human dataset is highly driven by the in-degree and incoming synapses (see Section 1.7, Fig. S7). Further, the algorithmic reconstruction utilized by authors is expected to underestimate the number of spines of dendritic trees (which contain an expected 80% of synapses made by analyzed neurons), thus a higher number of synapses may be expected than are actually observed on all cells. However, as we discussed in the adult *Drosophila* section, this effect is considered independent, thus all analyzed cells are equally effected, and therefore any comparison of the distributions we studied in this manuscript should remain valid.

We found that one of the 104 cells had an improperly formatted file structure, thus we analyzed 103 neurons in total, as well as the 320,000 synapses they make with each other, or with other cells in the volume. Given that these cells were released with measurements in nanometers, we converted the dataset units to micrometers prior to any analysis.

| Datasets | *C. elegans* | **Larva (F.F.)** | **Hemibrain (F.F.)** | **MANC (F.F.)** | **FlyWire (F.F.)** | **Zebrafish** | **MM3 (Mouse)** | **Human** |
|---|---|---|---|---|---|---|---|---|
| **Total Neurons (estimated)** | 302 | 10,000 | 170,000 | 170,000 | 170,000 | 100,000 | 70M | 86B |
| **Neurons in the reconstruction** | 302 | 3,066 | 22,699 | 22,107 | 129,278 | 2,587 | 52,059 | 46,378 |
| **Complete Neurons (Filtered)** | 300 | 3,037 | 3,898 | 12,379 | 129,278 | 2,587 | 22,350 | 103 |
| **Total Synapse Count (filtered)** | 20,905 | 599k | 3.2M | 27.5M | 33M | 1.39M | 135M | 320k |
| **Brain Volume (cubic microns)** | $3x10^6$ | $2.6 x10^6$ | $8x10^7$ | $8x10^7$ | $8x10^7$ | $1x10^8$ | $1.5x10^{11}$ | $1.1x10^{15}$ |
| **Dataset Bounding Vol. (cubic microns)** | $3x10^6$ | $2.61 x10^6$ | $2.04 x10^7$ | $8.17 x10^7$ | $7.8 x10^7$ | $2.4 x10^6$ | $5.09 x10^8$ | $1.07 x10^9$ |

**Supplemental Table 1: Dataset statistics and completeness.** For all datasets we highlight the relative completeness of electron microscopy reconstructions, compared to the total number of cells in the animal, as well as the remaining cells after filtering for quality. Total Neurons and Brain Volume represents an estimate of the total number of neurons or volume, respectively, in the whole body for invertebrates, and in the brain for zebrafish, mouse and human. Neurons in the reconstructions highlights the estimate provided by study authors of the total number of neurons, in many cases defined by number of soma found, in the reconstructed volume, whereas Complete Neurons and Total Synapse Count quantifies the number of neurons and synapse remaining after filtering (SI Section 1). Dataset Bounding Volume quantifies the reported size of the Electron Microscopy experiment from which the cell morphologies and connections were extracted. B, M, and k correspond to billion, million, and thousand, respectively.

#### Section 1.7: Symmetrization of Connectome Data

The synaptic connections between neurons have an inherent directionality, defined by pre and post synaptic terminals, that constrain the flow of information from one cell to the other. While network science is equipped to handle such directed and weighted connectivity, several key metrics are best established for undirected graphs. In order to challenge these standard assumptions, we chose to work with symmetrized versions of neuronal degree, synapse number, and weight. This also helps reduce the complexity of the narrative — otherwise for many quantities we would have to plot separately the incoming and outgoing values. To achieve this systematically, for the degree we considered all unique partners of a neuron, independent of whether they were incoming or outgoing connections. Similarly, we symmetrized the synapse number by summing all connections received and output by a cell. Finally, for the connection weight, for each neuron pair we summed the total number of connections made between them, independent of the direction.

At the same time, we acknowledge that the brain is a directed network and can be treated such. To this end, in Fig. S7, we plotted the distributions of degree and synapse number separately for the *incoming* and *outgoing* connections, finding that the incoming and outgoing distributions are largely indistinguishable in most organisms. The only exception is the human data, where the considerable discrepancy in peak between in- and out- distributions arises from decisions made while reconstructing cells: the authors focused on completing all incoming connections onto the proofread cells utilized in this analysis, as the dendritic arborage was better contained in the volume than the axonal projections of these neurons.

Finally, we assessed the fit of various distributions using KS statistics (Fig. S8), finding that the directed versions of these networks are also best fit by a lognormal distribution, the quality of the fits being comparable to the symmetrized version discussed in the main text. We thus find that the directed graph is consistent with a lognormal degree and synapse number across all datasets.

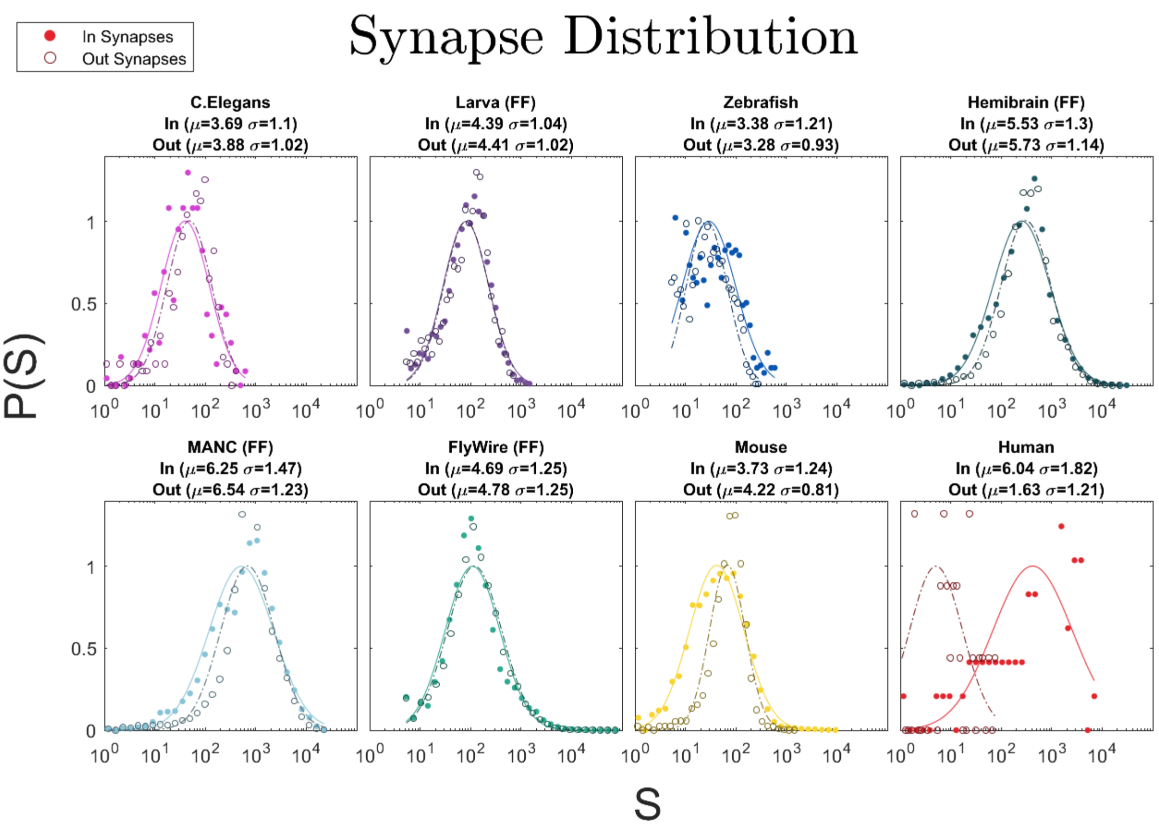
Synapse Distribution
In Synapses
Out Synapses
C.Elegans
In (μ=3.69 σ=1.1)
Out (μ=3.88 σ=1.02)
Larva (FF)
In (μ=4.39 σ=1.04)
Out (μ=4.41 σ=1.02)
Zebrafish
In (μ=3.38 σ=1.21)
Out (μ=3.28 σ=0.93)
Hemibrain (FF)
In (μ=5.53 σ=1.3)
Out (μ=5.73 σ=1.14)
MANC (FF)
In (μ=6.25 σ=1.47)
Out (μ=6.54 σ=1.23)
FlyWire (FF)
In (μ=4.69 σ=1.25)
Out (μ=4.78 σ=1.25)
Mouse
In (μ=3.73 σ=1.24)
Out (μ=4.22 σ=0.81)
Human
In (μ=6.04 σ=1.82)
Out (μ=1.63 σ=1.21)
P(S)
S

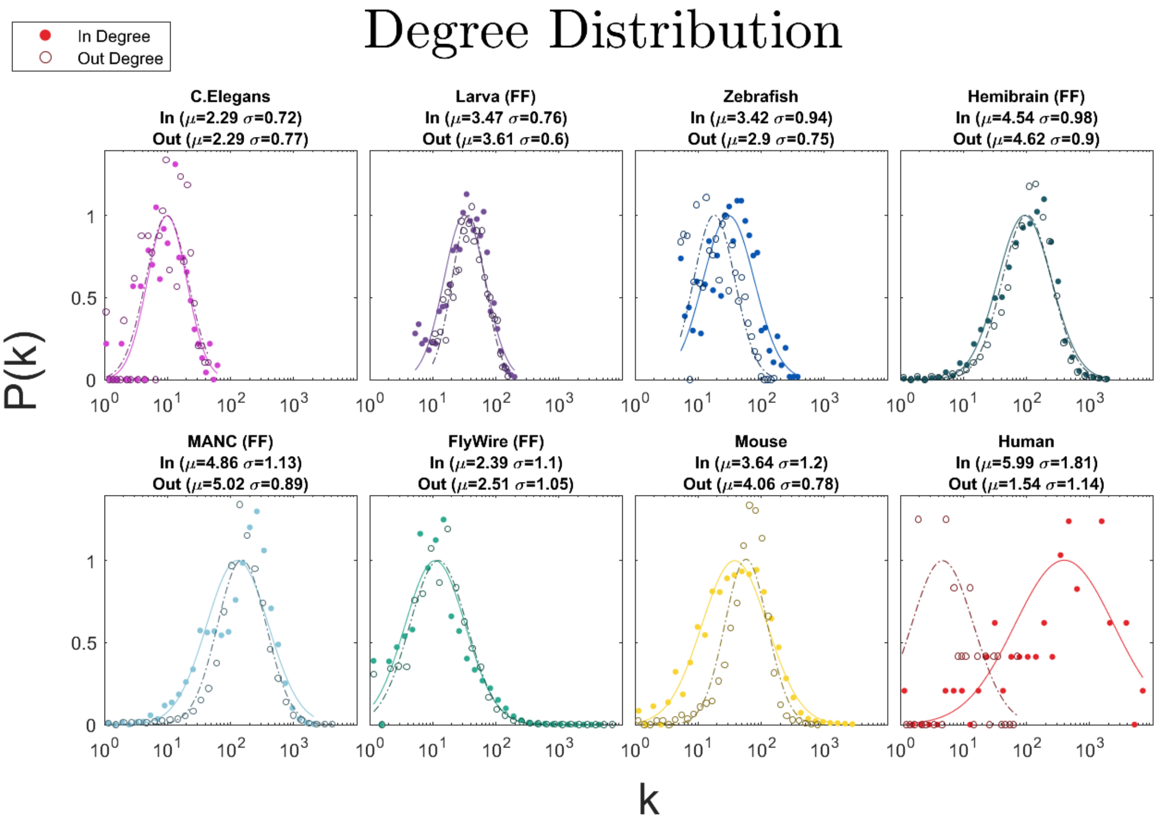
Degree Distribution
In Degree
Out Degree
C.Elegans
In (μ=2.29 σ=0.72)
Out (μ=2.29 σ=0.77)
Larva (FF)
In (μ=3.47 σ=0.76)
Out (μ=3.61 σ=0.6)
Zebrafish
In (μ=3.42 σ=0.94)
Out (μ=2.9 σ=0.75)
Hemibrain (FF)
In (μ=4.54 σ=0.98)
Out (μ=4.62 σ=0.9)
MANC (FF)
In (μ=4.86 σ=1.13)
Out (μ=5.02 σ=0.89)
FlyWire (FF)
In (μ=2.39 σ=1.1)
Out (μ=2.51 σ=1.05)
Mouse
In (μ=3.64 σ=1.2)
Out (μ=4.06 σ=0.78)
Human
In (μ=5.99 σ=1.81)
Out (μ=1.54 σ=1.14)
P(k)
k

**Supplemental Figure 7: Incoming and outgoing synapse and degree distributions.** In the top plots we show synapse distributions, with solid lines and points indicating incoming synapses and dashed lines and open points indicating outgoing synapses on neurons. In the bottom plots we show degree distributions, with similar notation. All distributions are well approximated by a lognormal shape regardless of directionality. In the larval fly and zebrafish connectomes, we remove neurons with less than 5 synapses from consideration, as this threshold is applied in the HemiBrain and FlyWire dataset releases by study authors. The considerable discrepancy in peak between human in- and out- distributions arises from decisions made while reconstructing cells: the authors of the study focused on completing all incoming connections onto the proofread cells utilized in this analysis, as the dendritic arborage was likely better contained in the volume than the axonal projections of these neurons. The same issue likely explains the smaller discrepancy between $S_{in}$ and $S_{out}$ (and $k_{in}$ and $k_{out}$) in the mouse datasets.

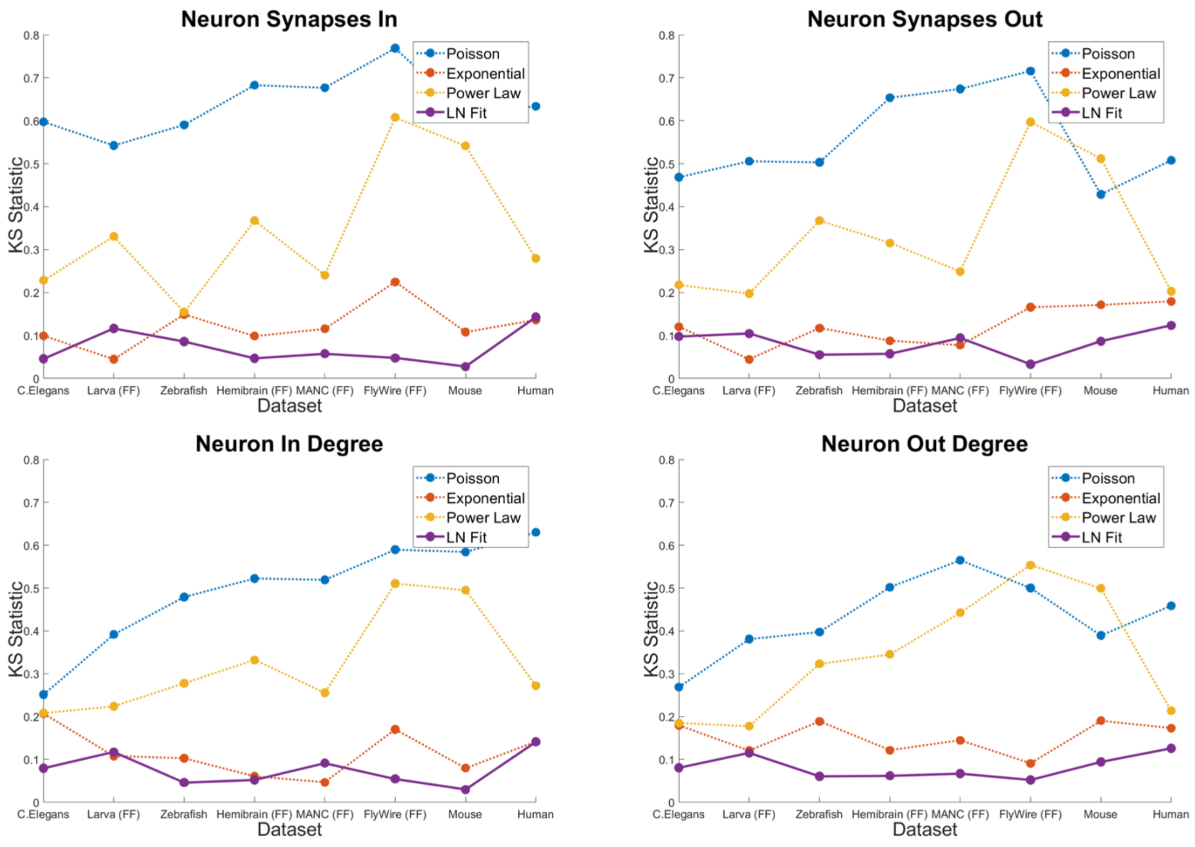

**Supplemental Figure 8: Incoming and outgoing synapse and degree KS statistics.** In the top row we show KS statistics for synapse distributions (incoming synapses on the left, outgoing synapses on the right), and in the bottom row we show KS statistics for degree distributions (incoming degrees on the left, outgoing synapses on the right). For most datasets the lognormal distribution has lower KS values over other distribution types, regardless of connection directionality.

### Section 2: The Lognormal Distribution

In this section we review the fundamental properties of the lognormal distribution, and discuss (and when needed, derive) the relationships we use to empirically falsify the relevance of the lognormal distribution to the brain connectome.

#### Section 2.1: Definition and Parameters

The lognormal distribution is a continuous probability distribution of a random variable whose logarithm is normally distributed. Specifically, if $X$ is a lognormally distributed random variable, then $Y = \log(X)$ follows a normal (Gaussian) distribution $N(\mu, \sigma)$, where $\mu$ is the logarithmic average and $\sigma$ in the logarithmic standard deviation, respectively. Note that throughout the text we use log(x) and ln(x) interchangeably, for natural logarithm. While the lognormal distribution is positively skewed in linear space, with a skewness that increases with $\sigma$, in the log-transformed space, $Y = \log(X)$ exhibits symmetry, with a skewness that approaches zero for large datasets (see Section 3.5).

The general form of the probability density function (PDF) for the lognormal distribution is:

$$f(x) = \frac{1}{(x-\theta)\sigma\sqrt{2\pi}} e^{\frac{-\log\left(\frac{x-\theta}{m}\right)^2}{2\sigma^2}} \qquad (1)$$

In this parameterization:

- $\theta$ is the location parameter, determining the horizontal position or "center" of the distribution.
- $m$ is the scale parameter, controlling the spread of the distribution.
- $\sigma$ is the shape parameter, influencing the skewness (asymmetry) and kurtosis (peakedness).

By setting $\theta = 0$ and $m = e^{\mu}$, we obtain the standard form of the lognormal distribution, also shown in Eq (2) in the manuscript:

$$f(x) = \frac{1}{x\sigma\sqrt{2\pi}} e^{\frac{-(\log(x)-\mu)^2}{2\sigma^2}} \tag{2}$$

It is important to note that the roles of the parameters differ between linear and log-transformed spaces. Linear-space shape and scale parameters become the scale and the location parameters in the log-space. Therefore, under a log-transformation, $\mu$ for the lognormal distribution acts as location and $\sigma$ acts as scale. This interplay between the parameters is the origin of the lognormal distribution's name, since the linear space $\mu$ and $\sigma$ for the normal distribution also map to location and scale respectively.

#### Section 2.2: Transformations of the Lognormal Distribution

When a lognormal random variable $X$ is multiplied by another lognormal random variable $Y$, the resulting product is also lognormally distributed. Specifically, if $P(X) = LN(\mu_X, \sigma_X^2)$ and $P(Y) = LN(\mu_Y, \sigma_Y^2)$, then:

$$P(XY) = LN(\mu_X + \mu_Y, \sigma_X^2 + \sigma_Y^2 + 2\rho_{XY}\sigma_X\sigma_Y) \tag{3}$$

Here, the standard deviation of the resulting distribution depends on the correlation $\rho_{XY}$ between X and Y. For our data, this correlation is nonzero (e.g., between length and local synapse density), hence it needs to be explicitly measured and accounted for.

When a lognormal random variable $X$ is multiplied by a power law function of the form $Y = \alpha X^\beta$, the resulting random variable also follows a lognormal distribution, with the following parameters:

$$P(X) = LN(\mu, \sigma^2), \;\; P\left(\alpha X^\beta\right) = LN(\log\alpha + \beta\mu, \beta^2\sigma^2) \tag{4}$$

For $\beta \approx 1$ and a scaling constant $\alpha \approx 1$, the power-law-transformed lognormal distribution closely resembles the original lognormal distribution.

#### Section 2.3: Maximum Degree

If the degree distribution follows a lognormal distribution $P(k) = LN(\mu_k, \sigma_k^2)$ then the maximum expected degree, $k_{max}$, will follow:

$$k_{max} = e^{\sqrt{2}\sigma\, erfc^{-1}\left(\frac{2}{N}\right)+\mu} \tag{5}$$

Where $erfc^{-1}$ is the inverse error function:

$$erfc(x) = 1 - \text{erf}(x) = \frac{2}{\sqrt{\pi}}\int_x^{\infty} e^{-t^2} dt\,. \tag{6}$$

We can write $k_{max}$'s dependence on N as:

$$k_{max} \approx e^{\mu+\sigma\sqrt{2\log(N)-\log\left[\frac{\log\left[\frac{N^2}{2\pi}\right]}{2\pi}\right]}} \approx e^{\mu+\sigma\sqrt{2\log N}} \tag{7}$$

#### Section 2.4: Moments of a Lognormal Distribution

The lognormal distribution has a moment generating function of the following form:

$$< X^n >= e^{n\mu+\frac{n^2\sigma^2}{2}} \tag{8}$$

Hence the first moment and second moments are

$$< X >= e^{\mu+\frac{\sigma^2}{2}}, < X^2 >= e^{2\mu+\frac{4\sigma^2}{2}}\,, \tag{9}$$

with a moment ratio of

$$\kappa \equiv \frac{< X^2 >}{< X >} = e^{\mu+\frac{3\sigma^2}{2}} =< x > e^{\frac{3\sigma^2}{2}} \tag{10}$$

that only depends on the parameters of the lognormal, being independent of the system size, *N*.

#### Section 2.5: Alternative Distributions and Fitting

To evaluate the suitability of the lognormal distribution, we compared it against several alternative distributions with known generative models in network science, including the Poisson, exponential, and power law distributions. We also compared it with the gamma and Weibull distributions, that lack a network generative mechanism, but are often used to fit fat-tailed distributions. The power law, Poisson, and exponential distributions are widely observed (and analytically predicted by suitable network models) in network science for the degree distribution. In contrast, the gamma and Weibull distributions are occasionally used to fit skewed data in log-space. While the Weibull distribution does emerge from network models, in some limiting cases[19], the condition for their emergence are hard to satisfy in real systems, as they require constraints that are rarely met. We have no theoretical network model to predict a Gamma distribution for *P(k)* or *P(S).* These comparisons allow us to assess the performance of the lognormal model relative to other commonly used and theoretically grounded distributions. The Gamma distribution can arise from summing a finite number of exponentially distributed independent variables — a condition unlikely to hold in networks of interdependent neurons. Similarly, the Negative Binomial results from combining a Poisson variable with a Gamma-distributed rate, but this too lacks a network-based interpretation. A key insight of network science is that not all statistical distributions are compatible with network structure. Once a distribution appears in empirical networks, its origin must be explained by a predictive generative mechanism.

Distributions such as the lognormal, gamma, and Weibull are parameterized by scale and shape parameters, whereas the normal distribution and many other linear-space distributions employ scale and location parameters. These differences in parameterization underscore the need to account for the underlying space (linear or log-transformed) when selecting an appropriate model. All parameter values as estimated by maximum likelihood estimation (MLE) for all distributions, unless stated otherwise.

#### Section 2.6: The Kolmogorov-Smirnov Test

The Kolmogorov-Smirnov (KS) test is a nonparametric statistical test used to compare the distribution of a sample to a reference probability distribution (one-sample KS test) or to compare the distributions of two independent samples (two-sample KS test). The one-sample test takes as input data and a distribution name, fit parameters for the distribution are calculated during the test. The two-sample KS test takes data from two different distributions and may require that samples from the two distributions be of equal length. The KS test can be used to assess how well data fits a particular distribution and to compare the fit quality of different distributions.

A one-sample KS test compares the empirical cumulative distribution function (ECDF) of the sample against a reference distribution (such as normal, exponential, etc.). The KS statistic is calculated as the maximum absolute difference between the ECDF and the CDF of the reference distribution: $D = \max\left(\left|F_{empirical}(x) - F_{reference}(x)\right|\right)$ , after calculating the reference distribution parameters, usually via maximum likelihood estimation (used in this paper). The actual KS "test" lies in comparing the obtained KS statistic to a reference value (often called the p-value or critical value) to determine if the null hypothesis is true, and of the empirical sample is drawn from the reference distribution.

Likewise, a two-sample KS test compares two CDFs, but instead of using a reference distribution, the two-sample test compares two sets of empirical data. By generating two empirical CDFs (ECDFs), and calculating their KS statistic, $D = \max\left(\left|F_{empirical1}(x) - F_{empirical2}(x)\right|\right)$, we can test whether the null hypothesis that the two distributions are identical is true by comparing them to a reference value (again called the p-value or critical value).

In this work, we used the Kolmogorov–Smirnov (KS) statistic to facilitate relative comparisons across the observed and the predicted distributions. A lower KS statistic indicated a closer fit of the dataset to a cumulative distribution function (CDF), making it an effective metric for evaluating fit quality across different measurement types, datasets, and candidate distributions. In fitting the CDFs of the various distributions compared in this work, we use maximum likelihood estimation (MLE) for all distribution types excluding the power law, which is estimated through a linear fit of log-transformed data. We have found that MLE parameter estimation significantly exaggerates the poorness of power law fits, and thus, we use the linear method to provide a fairer comparison between distributions.

While validating the null hypothesis of an exact match to a given distribution type would be ideal, this is often impractical in our context. The datasets analyzed are derived from coarse-grained skeletons of neurons and represent distributions composed of diverse cell types, making a perfect match to any single distribution highly unlikely. Instead, our goal was to compare the relative fit quality across candidate distributions. Metrics such as the KS statistic, alongside other non-parametric measures, are well-suited to this purpose.

#### Section 2.7: Other Goodness of Fit Tests

Cramér-von Mises Test: The Cramér-von Mises test evaluates the goodness-of-fit of a sample distribution $F_n(x)$ to a theoretical cumulative distribution function (CDF) $F(x)$, or it compares two empirical distributions, $F_n(x)$ and $G_m(x)$. The test is based on the integrated squared difference between the CDFs and is defined for both one-sample and two-sample scenarios. In this work, we use the one sample version of the test.

The Cramér-von Mises (CvM) test and the Kolmogorov-Smirnov (KS) test are both used to evaluate similarity of distributions, but they differ in focus and sensitivity. The KS test measures the maximum vertical difference between two cumulative distribution functions (CDFs), making it particularly sensitive to deviations near the center of the distributions. In contrast, the CvM test considers the integrated squared difference between the CDFs, giving equal weight to deviations across the entire range of the data. Consequently, the CvM test is generally more sensitive to differences in the tails of the distributions and provides a more holistic measure of fit, whereas the KS test emphasizes localized discrepancies.

In the one-sample test with a sample of n independent observations $\{x_1, x_2, \dots, x_n\}$, the Cramér-von Mises test statistic is given by:

$$T_n = n \int_{-\infty}^{\infty} [F_n(x) - F(x)]^2 \, dF(x) \quad (11)$$

where $F_n(x)$ is the empirical CDF of the sample, $F(x)$ is the reference distribution, and $T_n$ is the test statistic.

In practice, $T_n$ can be computed as:

$$T_n = \frac{1}{12n} + \sum_{i=1}^{n} \left[\frac{2i-1}{2n} - F(x_i)\right]^2 \quad (12)$$

where $x_i$ are the order statistics of the sample (sorted observations), with the summation term accounting for the deviation of the empirical CDF from the theoretical CDF.

Like with the KS test, this statistic can be compared to a threshold value to evaluate whether the null hypothesis is true.

Log-Likelihood: Log-likelihood is a statistical measure used to assess the goodness-of-fit of a probabilistic model to a given set of data. It is the natural logarithm of the likelihood function, which quantifies the probability of observing the data under a specific model and parameter set.

Given a set of independent observations $\{x_1, x_2, \dots, x_n\}$, and a probability density (or mass) function $f(x;\theta)$ parameterized by $\theta$, the likelihood function is:

$$L(\theta) = \prod_{i=1}^{n} f(x_i;\theta) \quad (13)$$

and the log-likelihood is:

$$l(\theta) = \log\big(L(\theta)\big) = \sum_{i=1}^{n} f(x_i;\theta) \quad (14)$$

While log-likelihood is typically used to optimize a parametric model's fit to data, it can also serve as the basis for non-parametric comparisons by leveraging maximum likelihood estimation (MLE) fit parameters. Although log-likelihood alone is not particularly informative as a standalone metric, it plays a crucial role in the computation of model comparison metrics such as the Akaike Information Criterion (AIC) and Bayesian Information Criterion (BIC). These metrics, widely used in model selection, provide a framework for evaluating likelihood ratios and comparing the relative performance of candidate models.

RMSE: Although root mean square error (RMSE) is not commonly employed for model comparison, it is a widely used metric for assessing the fit of a model to observed data. RMSE quantifies the average magnitude of prediction errors, offering a single value that succinctly summarizes a model's performance. This metric is particularly valuable in regression and predictive modeling, where the goal is to minimize deviations between predicted and observed values.

Given n observed values $\{x_1, x_2, \ldots, x_n\}$ and their corresponding model predictions $\{\hat{x}_1, \hat{x}_2, \ldots, \hat{x}_n\}$, RMSE is defined as:

$$RMSE = \sqrt{\sum_{i=1}^{n} (y_i - \hat{y}_i)^2} \tag{15}$$

Due to the squaring of errors, RMSE is very sensitive to outliers but is still a useful metric for quantifying prediction error for continuous outcomes in regression or predictive models. Like with log-likelihood, model parameters can be estimated via maximum-likelihood estimation, allowing us to use it as a non-parametric test.

Normalized Log Likelihood: Our data convincingly indicates that a lognormal offers the best fit to the length and degree distributions. However, as we illustrate in Fig. S12, for large variance, the lognormal can predict a monotonically decreasing function with a tail that can be fit to power law[20]. Prompted by the potential confusion over the possible fits, that often pervades the literature as well[21–24], we use additional statistical tools to assess whether the neuron length distribution (that, as we show in the paper, drives the rest of the measured distributions) is better approximated by a power-law distribution, indicative of a scale-free network, or a lognormal, indicative of a multiplicative network. To do so, we adopt a standard procedure of comparing the normalized log-likelihood ratio between the power-law and lognormal distributions. The normalized log likelihood, as defined by Vuong (1989)[25], is computed based on the mean and variance of the log-likelihood differences across multiple bootstrap resamples of the data:

$$R = \frac{\sum_i \left(\log L_1 (x_i) - \log L_2 (x_i)\right)}{\sqrt{Vn}} \tag{16}$$

where $L_1$ and $L_2$ are the likelihoods under the competing models, $V$ is the sample variance of the log-likelihood differences, and $n$ is the number of observations.

Following the methodology of Ref. [26], we use this normalized log-likelihood ratio to evaluate whether our data are better described by a power-law or lognormal distribution. The procedure involves generating numerous bootstrap resamples of the data, computing the log-likelihood for each distribution on each resample, and then analyzing the distribution of log-likelihood differences to assess model preference.

Using the publicly released code by Ref 20, we calculated the normalized log likelihood for all datasets under both distributions (with 1000 bootstrap samples each) to determine which distribution is a more appropriate fit. For all datasets and for all distribution, we find that the normalized log likelihood is negative, indicating that the lognormal outperforms the power law as a fit for each datasets across all four key measures (Fig. S9).

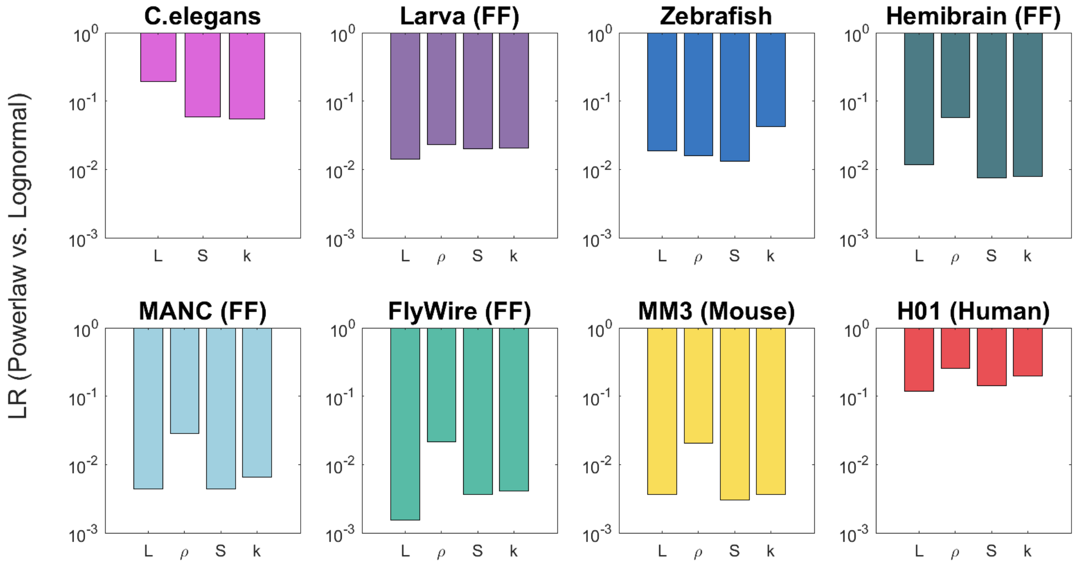

**Supplemental Figure 9: Normalized Likelihood Ratios Between the Powerlaw and Lognormal Distributions** For all datasets and quantities measured, we find that the normalized log likelihood is negative, suggesting that the lognormal outperforms the power law as a fit for our data.

### Section 2.8: Moment Estimation

When fitting a statistical model to data, it is crucial to compare multiple parameter estimation methods to ensure consistency and robustness of the inferred distribution. Different estimators can exhibit varying sensitivities to sample size, data dispersion, and underlying assumptions, which may impact the reliability of conclusions drawn from the analysis. In the case of lognormal distributions, the **maximum likelihood estimator** (MLE, used throughout our analysis) and the **method of moments** (MoM) are particularly well-suited due to their complementary strengths. The MLE is asymptotically efficient, meaning it achieves the lowest possible variance among unbiased estimators in large samples, and it naturally arises from the probabilistic structure of the data, making it a theoretically sound choice. In some distributional families (e.g., gamma), the MoM estimator provides a computationally straightforward, closed-form alternative that avoids iterative optimization and can be more robust to convergence issues in small or complex datasets. For the lognormal, however, both the MLE (eqs. 17–18) and MoM (eqs. 19–20) admit closed-form expressions, so MoM does not confer additional computational simplicity. Nevertheless, comparing the two remains valuable for assessing robustness to estimator-specific biases.

The MLE estimators for $\mu$ and $\sigma$ for a lognormal are the sample mean and sample variance of the log transformed data:

$$\mu_{MLE} = \frac{1}{n}\sum_{i=1}^{n} X_i \tag{17}$$

$$\sigma_{MLE}^2 = \frac{1}{n}\sum_{i=1}^{n}(X_i - \mu_{MLE})^2 \tag{18}$$

By contrast, the method of moments (MoM) relies on equating sample moments to theoretical moments of the lognormal distribution. Given the sample mean $\bar{X}$ and sample variance $S_X^2$ of the lognormal, the moments estimators for $\mu$ and $\sigma$ are:

$$\sigma_{MoM}^2 = \log\left(1 + \frac{S_X^2}{\bar{X}^2}\right) \tag{19}$$

$$\mu_{MoM} = \log(\bar{X}) - \frac{1}{2}\sigma_{MoM}^2 \tag{20}$$

We apply both estimators to all datasets and quantities measured. As Fig. S10 indicates, there is an excellent correlation between the Lognormal parameters determined for each of the distribution explored in the paper, providing confidence in the robustness of our parameter estimates. Specifically, we find that the MLE and MoM estimators yield similar values for $\mu$ across a wide range of sample sizes, suggesting that our results are not unduly influenced by estimator-specific biases. We observe small variations between the $\sigma$ of the two methods, however, these are less important as $\mu$ is the directly measurable parameter that determines the average of each distribution.

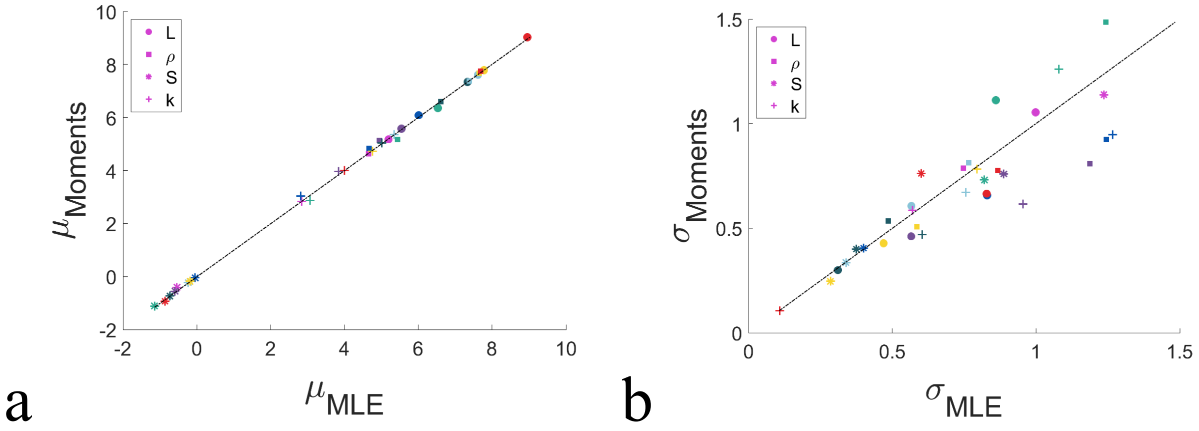

**Supplemental Figure 10: MLE vs Method of Moments Parameter Estimation (a)** We compare parameter estimation for $\mu$ using MLE and MoM and find excellent agreement between the values extracted by the methods. **(b)** We compare parameter estimation for $\sigma$ using MLE and method of MoM, and while agreement between the two methods is weaker, we do observe a strong correlations between the $\sigma$ measured by the two estimators.

#### Section 2.9: Log Binning

Logarithmic binning is the preferred method for analyzing and visualizing data that span several orders of magnitude. This approach divides data into bins whose widths increase logarithmically, such that each bin covers a constant ratio (e.g., $10^0$ to $10^1$, $10^1$ to $10^2$, and so on).

Linear binning, in contrast, uses bins of equal width. For data that spans several orders of magnitude, if we apply linear binning, a few bins dominate the histogram, containing an excessive number of data points, and other bins remain mostly empty, with insufficient datapoints to address their value and statistical significance. In Fig. S11 we highlight how linear binning and linear axis scales can be misleading, using examples of the FlyWire neuron length and synapse distributions, and how log-binning is able to extract from the data the full lognormal nature of the distribution.

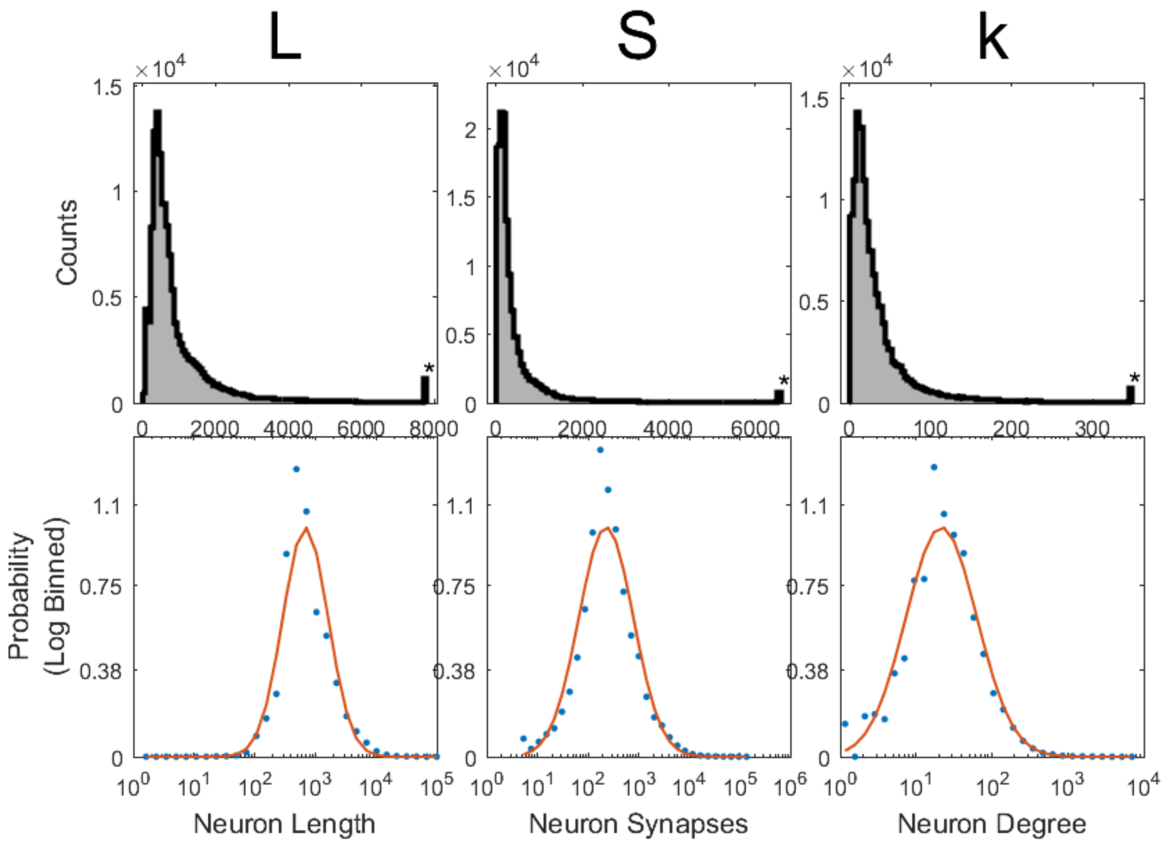

**Supplemental Figure 11: Linear versus log binning (FlyWire).**
In the top row, we show length, synapse count and degree under linear binning with a linear x-scale. This allows us to see only the right tail of the distribution, leaving the small values of *L*, *S*, and *k*, and their distribution, unresolved. By contrast Log-binning (bottom row) unveils the full extent over which *L*, *S*, and *k* vary, allowing us to see the lognormal nature of the FlyWire length, synapse count and degree distributions.

<u>Linear Binning Steps</u>
Linear binning divides the data range into bins of equal width.

1. Determine the range of the data:
   a. First find the minimum ($x_{min}$) and maximum ($x_{max}$) values in the dataset.
2. Specify the number of bins:
   a. Choose the number of bins ($N_{bins}$) based on the desired resolution.
3. Calculate bin width:
   a. Bin width is given by: $w_{bin} = \frac{x_{max} - x_{min}}{N_{bins}}$
4. Create bin edges:
   a. Bin edges are defined as: $Edges = \{x_{min}, x_{min} + w_{bin}, x_{min} + 2w_{bin}, \ldots, x_{max}\}$

5. Count data points:
    a. Count the number of data points in each bin.

<u>Log Binning Steps</u>

Logarithmic binning divides the data range into bins whose widths increase logarithmically.

1. Determine the range of the data:
    a. Exclude zero or negative values, as logarithmic binning requires positive data. Identify the minimum ($x_{min} > 0$) and maximum ($x_{max}$) values.
2. Specify the number of bins:
    a. Choose the number of bins ($N_{bins}$) based on the desired resolution.
3. Calculate bin width and bin edges:
    a. Bin edges are spaced logarithmically: $Edges = \{10^{\log_{10} x_{min} + i\Delta_{log}}\}$
    Where $i = 0, 1, 2, \ldots$, and $\Delta_{log} = \frac{\log_{10} x_{max} - \log_{10} x_{min}}{N_{bins}}$
4. Count data points:
    a. Count the number of data points in each logarithmic bin.

#### Section 2.10: Network Models and Their Degree Distributions

In the paper we test the data against four, theoretically well-founded distributions: the Poisson distribution expected for *P(k)* for a random network[27], the exponential distribution, predicted by a randomly growing network[28,29], a power law, expected for scale-free networks[28], and finally a lognormal distribution. In this section we review briefly the theoretical foundations for each of these distributions.

Random Networks

In the random network model, known also as the *Erdős-Rényi network*[27], we start with *N* isolated nodes (neurons), and connect each node pair independently with probability $p$. The average degree of a random network is $\langle k \rangle = p(N-1)$. In a random network the probability that node *i* has exactly *k* links follows the binomial distribution:

$$P(k) = \binom{N-1}{k} p^k (1-p)^{N-1-k} \tag{21}$$

The shape of this distribution depends on the system size *N* and the probability *p*.

Most real networks are sparse, $\langle k \rangle \ll N$, a limit for which the degree distribution of a random network is well approximated by the Poisson distribution

$$P(k) = e^{-\langle k \rangle} \frac{\langle k \rangle^k}{k!} \tag{22}$$

This form does not explicitly depend on the number of nodes *N*, therefore the degree distribution of random networks of different sizes but the same average degree $\langle k \rangle$ are indistinguishable from each other.

Scale-Free Networks

The random network model assumes that we have a *fixed* number of nodes, *N,* connected randomly. Yet, in real networks *the number of nodes continually grows thanks to the addition of new nodes, and new nodes prefer to link to the more connected nodes*, a process called *preferential attachment*. The recognition that growth and preferential attachment coexist in real networks has inspired the *Barabási-Albert* (BA) model[30], which starts with $m_0$ nodes, the links between which are chosen arbitrarily, as long as each node has at least one link. The network develops following two steps:

**a) Growth:** At each timestep we add a new node with $m$ ($\leq m_0$, the initial number of nodes) links that connect the new node to $m$ nodes already in the network.

**b) Preferential attachment:** The probability $\Pi(k)$ that a link of the new node connects to node $i$ depends on the degree $k_i$ as $\Pi(k_i) = k_i \sum_j k_j$

The degree distribution of the Barabási-Albert network can be calculated exactly[31,32],

$$P(k) = \frac{2m(m+1)}{k(k+1)(k+2)}. \tag{23}$$

Therefore, for large $k$ the degree distribution follows a power law $P(k) \sim k^{-\gamma}$ with degree exponent $\gamma$=3. Subsequent modeling efforts have built models that can vary $\gamma$, the degree exponent[24,29,33,34].

Growth Without Preferential Attachment (Model A)

While virtually all real networks emerge through a growth process, not all are governed by preferential attachment. To test the role of growth on the network topology, we keep the growing character of the network and eliminate preferential attachment, starting with $m_0$ nodes and evolve following these steps (known as Model A[28,29]):

**a)** At each time step we add a new node with $m(\leq m_0)$ links that connect to $m$ nodes added earlier.

**b)** The probability that a new node links to a node with degree $k_i$ is $\Pi(k_i) = \frac{1}{m_0+t-1}$.

That is, $\Pi(k_i)$ is independent of $k_i$, indicating that new nodes choose randomly the nodes they link to. In this case the degree distribution follows an exponential function,

$$P(k) = \frac{e}{m} \exp\left(-\frac{k}{m}\right), \tag{24}$$

which decays much faster than a power law, hence it does not support hubs.

Lognormals in Network Science

The potential of lognormal degree distributions has been long discussed in network science[20,35,36], given their ability to fit fat-tailed data (Fig. S12). Indeed, a recent reanalysis of datasets utilized in network science found that 50% of *P(k)*s are consistent with a lognormal[23]. However, in network science to establish the nature of the degree distribution, fitting is not sufficient — we also need a mechanistic model to offer a plausible explanation of the observed *P(k)*. Consequently, lognormals are rarely used in network science, because we lack theoretical justification for their emergence in real networks. Yet, a few rather specific generative models for lognormal *P(k)* have been proposed in the context of social networks, which we review next.

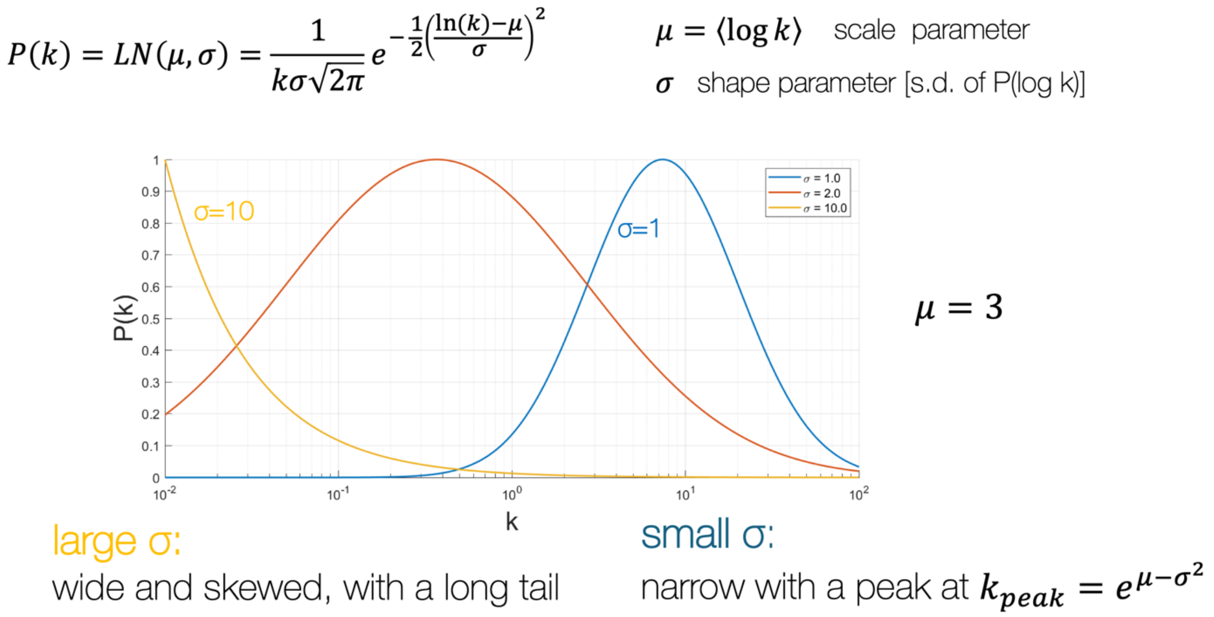

**Supplemental Figure 12: Lognormals for fitting heavy tails.** The shape of a lognormal as a function of $\sigma$. For small $\sigma$ the lognormal has a clear peak with tails extending towards the high and low regime, as illustrated for the case of $\sigma = 1$ and $\sigma = 2$ in the figure. For large $\sigma$, however, we only observe the decaying right tail, which is often used to fit a power law.

Modeling Citations: A generative model was proposed to address the potential lognormal nature of the degree distribution of paper citations from 110 years of articles published in the journal *Physical Review*[36]. The authors proposed a preferential attachment-based model, a modification of the BA model, to explain the potential emergence of a lognormal degree distribution. In passing, Redner suggested a modified linear preferential attachment rule[36], proposing the form:

$$\Pi_k = \frac{k+1}{1 + a\log(k+1)}, \tag{25}$$

which, as it was shown[35], can lead to a lognormal distribution. Nevertheless, these models were not investigated in more depth, and remain specific to the citation network.

Homophilic Connectivity in Social Networks: A model that can generate lognormals by introducing a large number of node attributes emerged from efforts to capture homophilic connections in social networks. Multiplicative attribute graphs (MAG)[37] were proposed as a class of stochastic network models that capture the interactions between node attributes. Specifically, each node has a vector of categorical attributes associated with it. Attribute values of two nodes are then combined in order to predict the emergence of a link between them. The choice of interactions between node attributes can lead to homophily (preference for the same features) as well as heterophily (preference for different features). The MAG paper found that for appropriate choice of the attributes, the tail of the degree distribution presented a lognormal.

Taken together, while a few models (and remarkably few compared to the extensive network science literature) can generate lognormal *P(k)*, the existing modeling efforts remain limited to social systems, and their wider relevance remains to be established. Most importantly, their assumptions and features do not apply to neuronal wiring. In the context of the connectome, to accept the relevance of the lognormals, the fits are not sufficient — we also need to identify the multiplicative process that is responsible for it, and we need to be able to analytically derive *P(k)* based on processes that are known to be present in the brain.

### Section 2.11: Connectome Configuration Model (CCM)

Generative models, designed to capture some key features of a particular network, while ignoring other less consequential properties, play a key role in network science. While a comprehensive generative model of the connectome is still elusive, our results naturally lend themselves to a null model that helps us explore the validity of the scaling laws derived in the manuscript, and offers the foundations of a more accurate modeling framework. The **Connectome Configuration Model (CCM)** is based on our central hypothesis that the brain's architecture is driven by the physical nature (specifically, the length) of individual neurons and is defined as follows:

(1) *Neurons*: We generate incomplete trees using the Galton-Watson (GW) process to mimic the tree structures of individual neurons. We set the survival probability of each branch at $p_s = p_c + \epsilon + \eta$, where $p_c$=1/2 is the critical threshold predicted by the analytic calculation (see SI 4.2). The parameter $\epsilon$ controls the expected neuron length and $\eta$ represents the noise amplitude, chosen to have zero mean, capturing the variability $\sigma_L$ (see SI 4.2). To best mimic the real tree sizes observed in the data, we also fix a starting number of branches $a$ and a maximum number of layers $n$ for the simulation. The tree sizes generated by the GW process ($\mu_L = \langle \log L \rangle$) are then converted to neuron lengths by multiplying by a constant segment length, that for simplicity we choose to be $l = 1$.

(2) *Synapses*: All branches are populated by synaptic terminals by multiplying neuron length with a constant synapse density $\rho$, following Eq (1) in the main text. We randomly designate half of these terminals as presynaptic and the other half as postsynaptic.

(3) *Network Construction (Configuration Model)*: Each presynaptic terminal is connected to a randomly chosen postsynaptic neuron.

The CCM intentionally simplifies many key aspects of neurons and their connectivity:

- **Rule 1** ignores the existence of multiple neuron classes and the distinct survival probabilities (and layer dependent survival functions) characterizing each class, that we document in Fig. 5 in the main text. It also overlooks the fact that branches near the soma often lack synaptic terminals. Finally, by choosing *l = 1*, it ignores the variability in the length of the individual branches. Needless to say, these features can be easily implemented, if desired.

- **Rule 2** ignores the difference between axons and dendrites. Indeed, the canonical view of a neuron consists of a soma, from which a single axon emerges that carries all presynaptic terminals, along with separate dendrites that receive signals from presynaptic neurons. However, in the fruit fly, most branches contain both presynaptic and postsynaptic terminals, though some branches exhibit enrichment of one type over the other. Ignoring this enrichment, we randomly assign half of the boutons to be presynaptic and half to be postsynaptic, hence ignoring the distinction between axons and dendrites. This too can be corrected, if desired.
- **Rule 3** disregards the fact that synapses can only form between neurons in close physical proximity. As such, this configuration approach allows us to assess the role of the neuron's length, neglecting spatial constraints imposed by physicality. We need more accurate modeling of the physical layout of the neurons to address this.

All of these limitations can be relaxed using more complex assumptions. Yet, despite these simplifications the CCM model captures many key features of the connectome, reproducing several common features of the eight datasets studied in the paper:

(a) **Neuron Length Distribution:** In line with our theoretical prediction (see SI 4.3), the CCM model predicts a lognormal neuron skeleton length distribution *P(L)* (Fig. S13a). Further, with the suitable choice of the $(\epsilon, \eta)$ parameters, the model can offer a reasonable approximation for the empirically observed *P(L)* for the FlyWire data. In Fig. S13a we choose $\epsilon = 0$ and $\eta = 0.1$, to mimic noise levels in the data. We also fix $a = 24$ and $n = 15$. With these parameters we obtain $\mu_L^{CCM} = 6.53$, identical to the value observed in the FlyWire data, and $\sigma_L^{CCM} = 0.85$, which is slightly smaller than the empirically observed $\sigma_L = 0.65$, a difference that is largely immaterial for our subsequent analysis.

**(b) Synapse Density Distribution:** By choosing a fixed density by design we have a delta function for the synapse density distribution (Fig. S13b) rather than the lognormal observed in the data. However, that can be easily corrected, if desired, by choosing a lognormal $P(\rho)$ as input. This difference is immaterial for now, so we are staying with the fixed synapse density.

**(c) Strength (Synapse) Distribution:** The model correctly predicts a lognormal *P(S)* (Fig. S13c). According to Eqs. (3) and (4) in the main text, the parameters of *P(S)* are expected to be governed by the parameters of *P(L)* and *P(ρ)*. To check if this is the case, in Table S2 we show the empirically measured $(\mu, \sigma)$ parameters for *P(L)* and *P(ρ)*, and *P(S),* along with the values of $(\mu, \sigma)$ predicted by Main Text Eqs. (3) and (4). We find an excellent agreement between the predicted and the observed values. While we observe a smaller $\sigma_S$ compared to the empirical data, this difference is rooted in the constant synapse density. In other words, the model is internally self-consistent, hence while some of its parameters are different from those observed in the data, it closely follows the theory outlined in the paper.

**Supplemental Table 2: CCM Synapse and Degree Moments Prediction**

| $\epsilon = 0,$ $\eta = 0.1$ | $L\ measured$ | $\rho\ measured$ | $corr(L, \rho)$ | $S\ measured$ | $S\ predicted$ |
|---|---|---|---|---|---|
| $\mu$ | $\mu_L = 6.53$ | $\mu_\rho = -1.1$ | 0 | $\mu_S = 5.43$ | $\mu_{S^*} = \mu_L + \mu_\rho = 5.43$ |
| $\sigma$ | $\sigma_L = 0.65$ | $\sigma_\rho = 0$ | 0 | $\sigma_S = 0.65$ | $\sigma_{S^*} = \sqrt{\sigma_L^2 + \sigma_\rho^2 + 2\sigma_L\sigma_\rho corr(L, \rho)} = 0.65$ |

**(d) Degree Distribution:** The model correctly predicts a lognormal degree distribution *P(k)*, albeit with a smaller $\mu_k$ than observed in the data (Fig. S13d). The difference is rooted in the fact that we ignore the agglomeration of the *in* or *out*-synapses on the specific branches of the neuron. Yet, once again the model parameters are self-consistent—indeed, as we show in Table S3, the analytically predicted parameters of *P(k)* (see Eqs. 6,7 in the manuscript) are in perfect agreement with the empirically measured parameters.

**Supplemental Table 3: CCM Synapse and Degree Moments Prediction**

| $\epsilon = 0,$ $\eta = 0.1$ | $S\ measured$ | $\beta$ | $\alpha$ | $k\ measured$ | $k\ predicted$ |
|---|---|---|---|---|---|
| $\mu$ | 0.65 | 1 | 1 | $\mu_k = 5.43$ | $\mu_{\beta S^\alpha} = 5.43$ |
| $\sigma$ | 0.65 | 1 | 1 | $\sigma_k = 0.65$ | $\sigma_{\beta S^\alpha} = 0.65$ |

**(e) Synapse dependence on the neuronal length: Scaling Law (1).** In the CCM model the number of synapses depends linearly on the length of the neuron in line with the asymptotic behavior observed in empirical data (Fig. S13e).

**(f) Degree and Synapse Scaling: Scaling Law (5).** The CCM displays a linear dependence between degrees and the number of synapses of individual neurons, i.e. it corresponds to $\beta = 1$ in Main Text Eq 5, in contrast with the slightly sublinear dependence ($\beta < 1$) observed in the real data. This indicates that we need to account in a more precise manner for the physicality of the neurons, that determine (and limit) proximity between neuron pairs. Modeling this process remains an open question in connectomics.

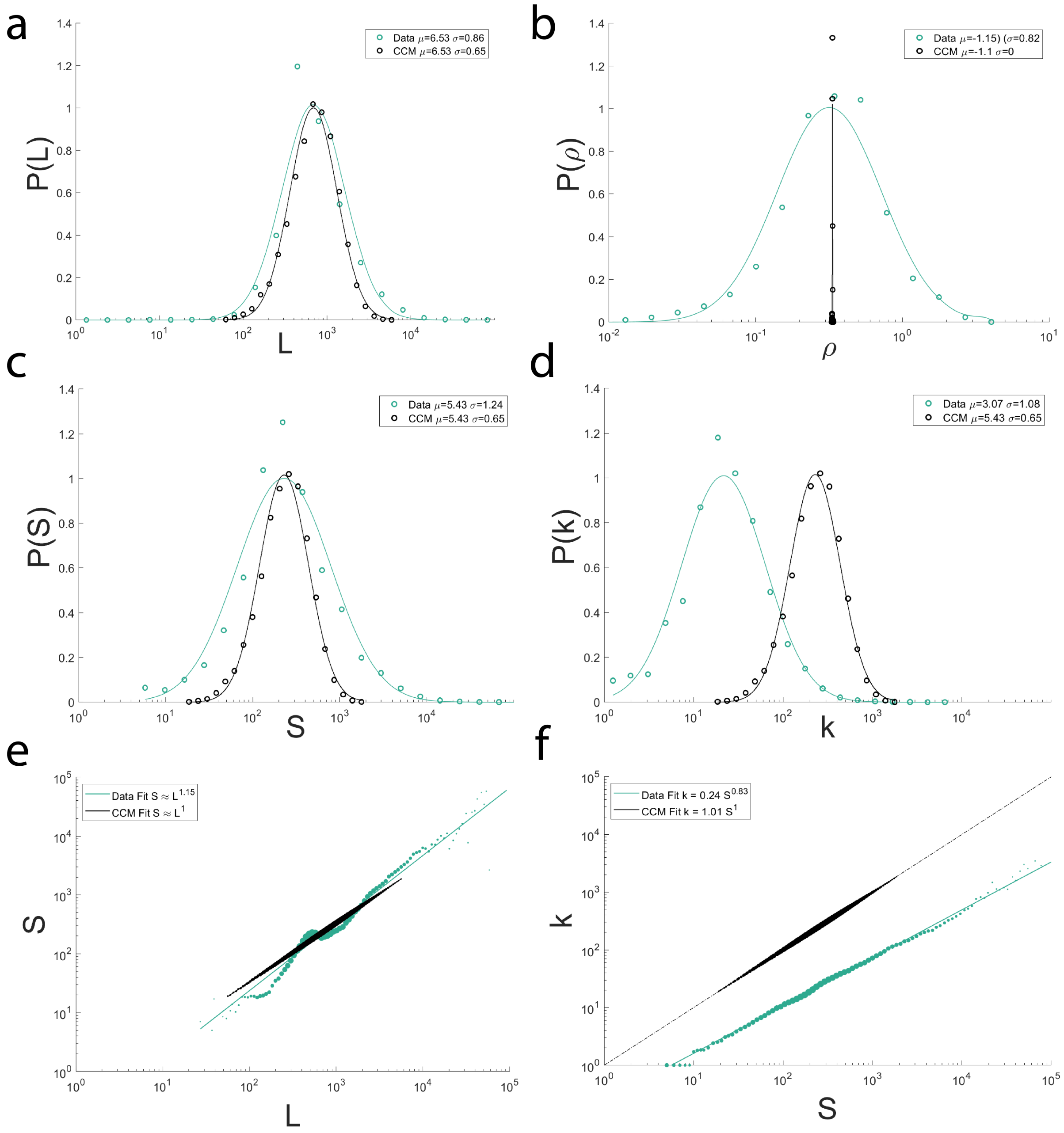

**Supplemental Figure 13: The Connectome Configuration Model (CCM). (a)** The CCM model for $\epsilon = 0,\ \eta = 0.1$ generates a lognormal distribution with the correct $\mu$ and a slightly smaller $\sigma$ than the data. **(b)** The CCM model by design has a delta function for *P(ρ),* with the same peak. **(c)** The model reproduces the lognormal nature of *P(S)*, but given the single synaptic density value its $\sigma$ is smaller. **(d)** *P(k)* of the CCM model is lognormal, with higher $\mu$ than that of the data, as randomizing connections makes high connection weights less likely. **(e)** The CCM model capture the scaling law (1). **(f)** The CCM model provides a linear k vs S scaling, while the data is slightly sub linear.

Varying the Parameters of the CCM Model

The results shown above were for model parameters $\epsilon = 0$ and $\eta = 0.1$. However, the self-consistency of the model goes beyond these parameters. To test this, we have varied $\epsilon$ and $\eta$, and for each parameter pair we measured the respective $(\mu, \sigma)$ parameters for the distributions *P(L)* and *P(ρ), P(S)* and *P(k)*. In Fig. S14 we show predicted vs the measured distribution parameters for the different ($\epsilon$ , η) models, finding that independent of our choice of the model parameters, Eqs. (3) and (4), and (6) and (7) that link the various neuron characteristics remain valid.

In summary, the CCM model and the Synapse Configuration Model (SCM), discussed next, reproduces some of the key network characteristic of the connectome, generating a lognormal network. As such, it serves as a starting point to explore higher order properties of the connectome. It can help us understand which of the observed features are rooted in the theoretical framework developed in this paper, and which features require a deeper understanding of physicality, as well as a biologically more accurate modeling of the neuron's ability to synapse with each other.

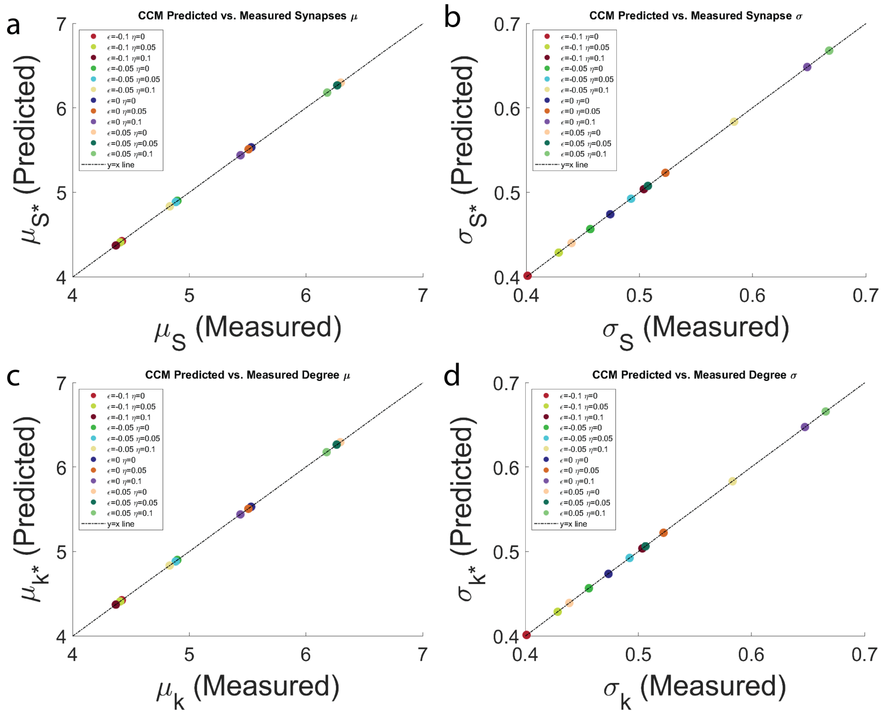

**Supplemental Figure 14: CCM Predicted and Measured Lognormal Parameters**
**(a)** The CCM model predicted $\mu_S$, confirming that eq. 3 holds for all parameter ($\epsilon$,η) pairs.
**(b)** The CCM model predicted $\sigma_S$, confirming that eq.4 holds for all parameter ($\epsilon$,η) pairs.
**(c)** The CCM model predicted $\mu_k$, confirming that eq.6 holds for all parameter ($\epsilon$,η) pairs.
**(d)** The CCM model predicted $\sigma_k$, confirming that eq.7 holds for all parameter ($\epsilon$,η) pairs.

#### Section 2.12: The Synapse Configuration Model (SCM)

To evaluate the role of randomness and physicality, we also find useful to explore the **Synapse Configuration Model (SCM)**. The model is based on the FlyWire empirical data, which includes 129,278 neurons. The SCM preserves the total number of neurons, along with the number of synapses on each neuron and their identities (pre- or postsynaptic). It randomizes, however, the connections between them using **Rule 3** of the CCM model (configuration rule)—connecting each presynaptic terminal to a randomly chosen postsynaptic terminal. Hence the SCM model maintains the full neuronal organization while randomizing connectivity. As such, it automatically reproduces many features of the original data (Fig. S15), only generating shifts in the degree distribution (panel d). It also captures the scaling laws (1) and (6), predicting a linear dependence in each case, indicating that once again the sublinear (6) is rooted in the physical packing of the neurons.

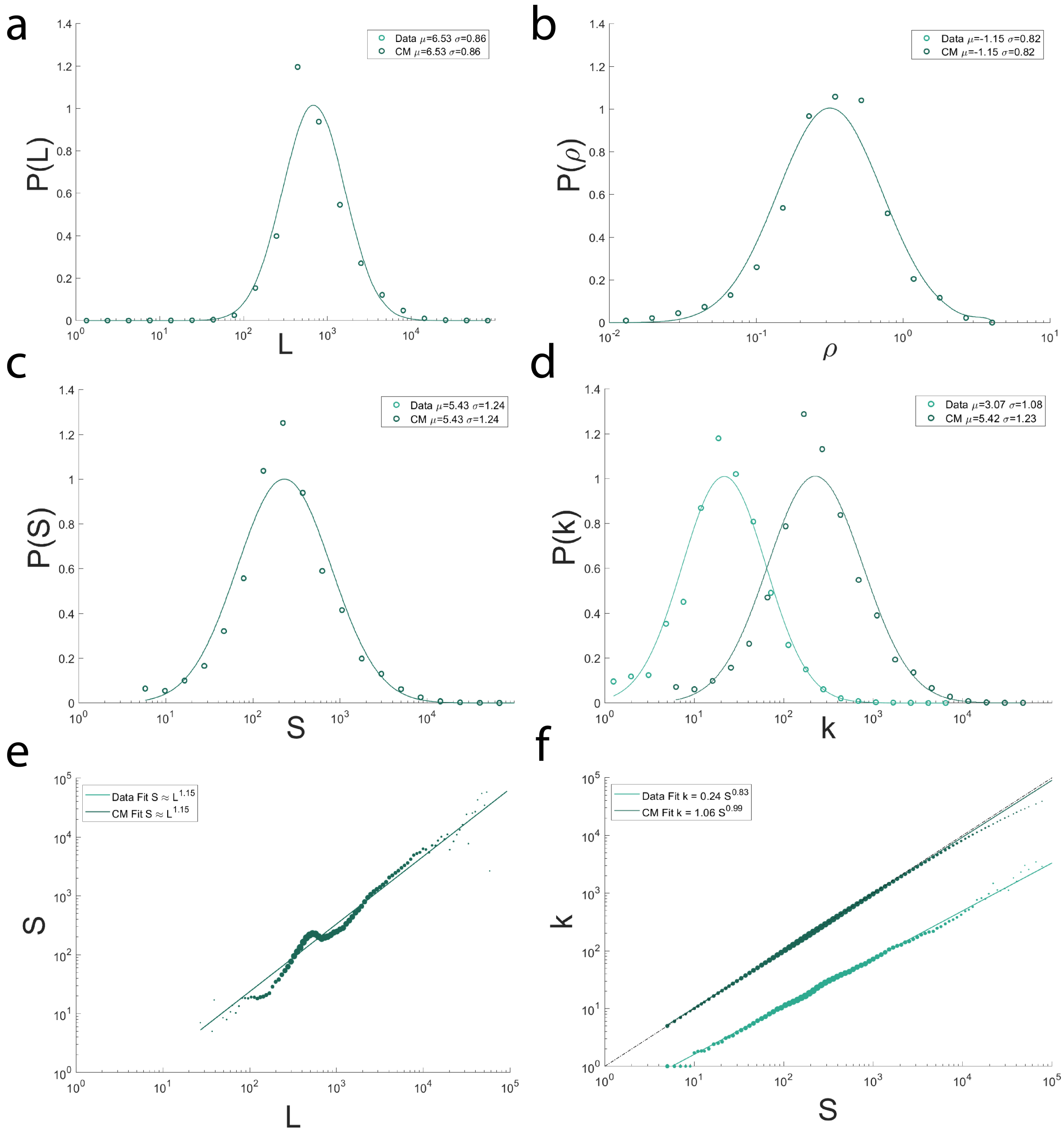

**Supplemental Figure 15: The Synaptic Configuration Model (SCM).**
**(a)** The length distribution of SCM trivially generates a lognormal *P(L)* that matches the FlyWire dataset, as this value is conserved in the model. **(b)** The synapse density distribution of SCM trivially generates a lognormal *P(ρ)* that matches the FlyWire dataset, as this value is conserved in the model. **(c)** The synapse number distribution of SCM trivially generates a lognormal *P(S)* that matches the FlyWire dataset, as this value is conserved in the model. **(d)** *P(k)* of the SCM model is lognormal, with higher $\mu$ than that of the data, as randomizing connections makes high connection weights less likely. **(e)** The SCM trivially capture the scaling law (1), as *S* and *L* values are conserved in this model. **(f)** The SCM model provides a linear k vs S scaling, while the data is slightly sub linear.

### Section 3: Unveiling the Properties of the Connectome

#### Section 3.1: Neuron Length Distribution *P(L)*

The size of a neuron, *L*, is one of its most fundamental properties, as the identity and number of a cell's partners depends on the physical surface area available for it to make connections. Indeed, a larger neuron has the potential to make more synapses, thus can have a higher degree. In our study, we parameterized neuron size by the total length of its neurites, which we calculated by summing over all segments of downloaded neuron skeletons. Specifically, the commonly used .swc format breaks down neurons into a series of points. Each point, except for the "source" point, has a "parent" point, to which it connects. Thus, to arrive at the length of a neuron, we summed over the Euclidean distance between each provided point and its parent point. By applying this procedure to all neurons in a dataset, we arrive to the distribution of neuron lengths for each profiled system, *P(L)*. Next, we apply log binning to each dataset (see SI 2.8), and fit several candidate distributions to them. Fig. S16 shows the obtained distributions, along with the best lognormal fit to each distribution. Further, we observe that neuron size increases with brain size, over nearly two orders of magnitude, which is to be expected (Fig. S17).

We further note that the observed distributions have unique features that result in local deviations from the idealized form of a lognormal distribution. Some of these deviations likely reflect their composite nature, as seen in FlyWire with the different cell types characterized by different length scales, and are also affected by data incompleteness, that induces skewness (see Section 3.5). Nonetheless, statistical analyses, including the Kolmogorov–Smirnov (KS) test, identify the lognormal distribution as the overall best fitting model among the fundamental models typically studied in network science (see SI 2.9, Fig. S16).

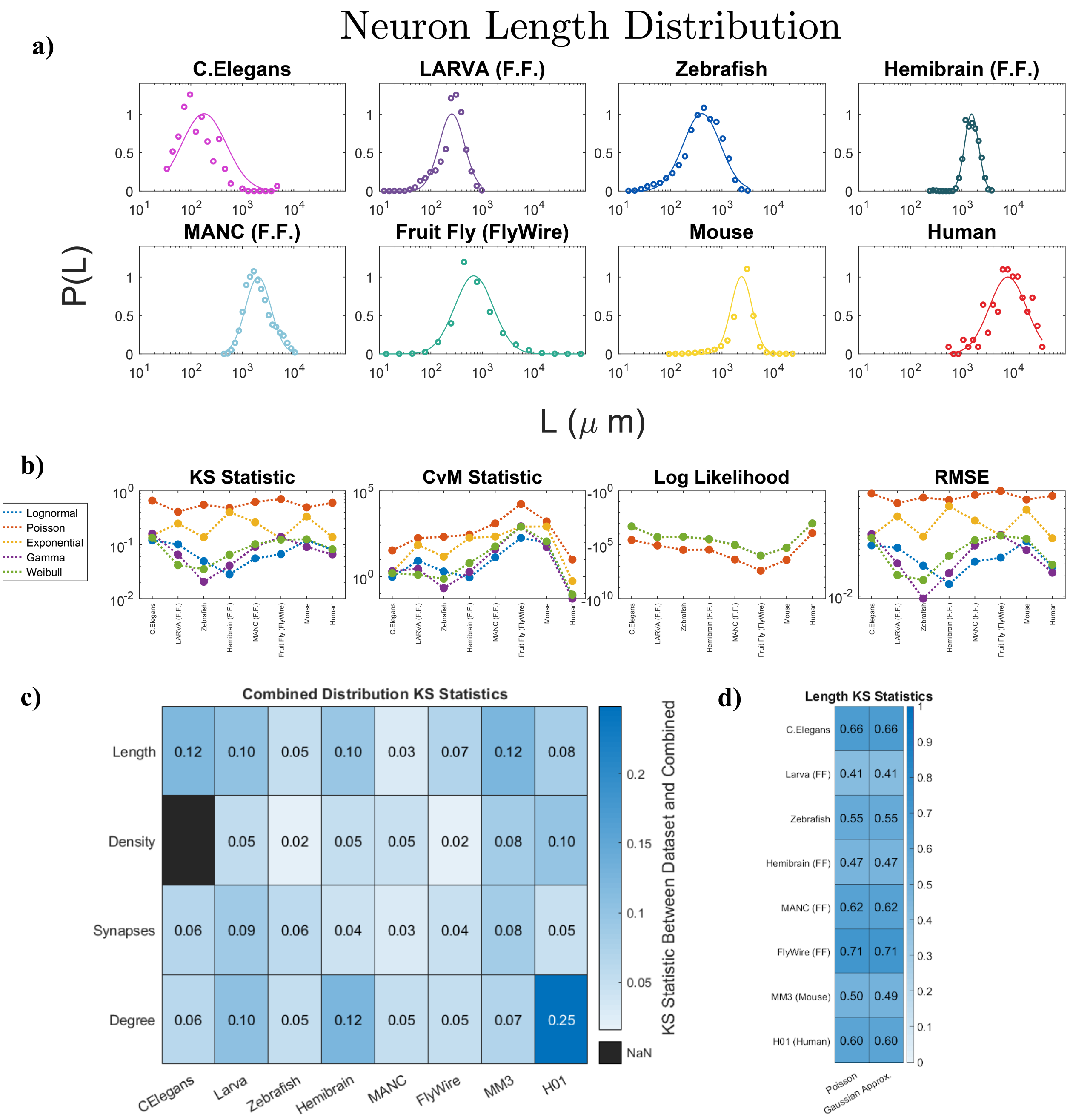

**Supplemental Figure 16: Neuron length distributions across organisms.**
**a)** Length distribution of all analyzed datasets, plotted on a log-x scale, with probability plotted on the (linear) y-axis. Scatter points show the empirical data and colored lines depict the best lognormal fit.
**b)** For each dataset, we compare the goodness-of-fit using the Kolmogorov-Smirnov (KS) test, the Cramér-von Mises (CvM) test, the probabilistic log-likelihood measure, and the root mean square error (RMSE). In all cases, the y-axis displays the fit statistics, with a lower value indicating the better fit, and the x-axis indicates the dataset for which the fit was performed. Under log-likelihood, we find that Lognormal, Exponential, Weibull and Gamma distributions overlap. Note that in some

panels we fit a discrete Poisson to a continuous variable. This is justified since for all datasets $\lambda \geq 20$, where Poisson closely approximates a normal distribution, making the KS test appropriate.
**c)** Kolmogorov–Smirnov (KS) statistics comparing individual datasets (columns) to the combined distribution for four key neuronal features: length, density, synapse number, and degree. To ensure statistical fairness, equal-sized samples (matching the smallest dataset) were drawn without replacement from each distribution before comparison. The consistently low KS values for length, synapse number, and density confirm the robustness of the rescaling-based data collapse observed in Figure 2. The primary exception is the degree distribution in the H01 (Human) dataset in the case of degree P(k) shows a poorer match due to its severely limited sample size, a constraint affecting most measurements in this dataset.
**d)** For sufficiently large parameter values (e.g., $\lambda \geq 20$, as is the case even for C. elegans), the Poisson closely approximates a Gaussian (continuous) distribution. We explicitly test this assumption, by comparing the Poisson KS statistics against the corresponding Gaussian approximation, with $\mu = \lambda$. We find that the Gaussian approximation is either only marginally better than or offers comparable fit quality to the Poisson.

| | Lognormal | Poisson | Exponential | Gamma | Weibull |
|---|---|---|---|---|---|
| KS | **0.078000854** | 0.565544527 | 0.225284566 | 0.084639645 | 0.089041286 |
| CVM | **37.46396944** | 2606.00569 | 265.9322915 | 118.6852665 | 124.0058136 |
| LL | **25269752.55** | 21553723.41 | 25264759.72 | 25267362.43 | 25266776.05 |
| RMSE | **0.039331264** | 0.299880728 | 0.125061719 | 0.044154723 | 0.04970582 |

**Supplemental Table 4: Mean Length Test Statistics.** For the KS, CvM, Log likelihood and RMSE tests visualized in the Figure above, we calculated the mean test statistic across all organisms, finding that the Lognormal emerges as the top-performing model across all four measures.

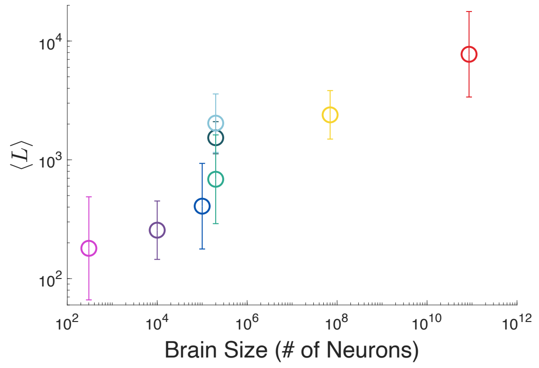

**Supplemental Figure 17: Correlation between brain size and neuron length.** Comparison of average neuron length with total neuron number (see Table S1) for all eight datasets (see Fig. S16 for color code).

#### Section 3.2: Local Synapse Density Distribution *P(ρ)*

The synapse density of a neuron is defined in this paper as the average number of synapses per micron of neuronal length. However, calculating synapse density as the total number of synapses on a neuron divided by its total length can be misleading. Indeed, neurons often exhibit significant regional variation, with some areas densely populated with synapses and others devoid of them. In mouse and human neurons, axons, which typically make outgoing connections, will have regions where they make no connections at all, as they serve to project long distances, while the dendrites, which mainly receive connections, typically make synapses only on protrusions from the brain branches, called spines. Such synaptic density variability is also evident in the fruit fly, where synapse density is notably low near the soma (Fig. S18b). Therefore, a more nuanced measure of synapse density must focus on regions where synapses are actually present.

To address this, we employed two methods to estimate a local synapse density:
Method 1: We measured the distances between synapses and their neighbors on the same neuron (Fig. S18c). By averaging these distances across many random samples, we derived an estimate of the typical spacing between synapses, which inherently de-emphasizes regions devoid of synapses.

Method 2: We performed random walks originating from individual synapses, traversing a fixed length in one direction (10 microns in this study). By counting the number of synapses encountered during each walk (Fig. S18d), we calculated a local synapse-per-micron density, a procedure that excludes large empty regions, as these walks cannot penetrate far into synapse-free zones. Averaging the results from a large number of the random walks (1000 per neuron) provided a robust estimate of local synapse density.

While the two approaches offered comparable results (Fig. S18e), in the main text we present the results of Method 2, the random walk approach, as we considered averaging over multiple local synapses, versus simply counting the nearest one, a more robust metric.

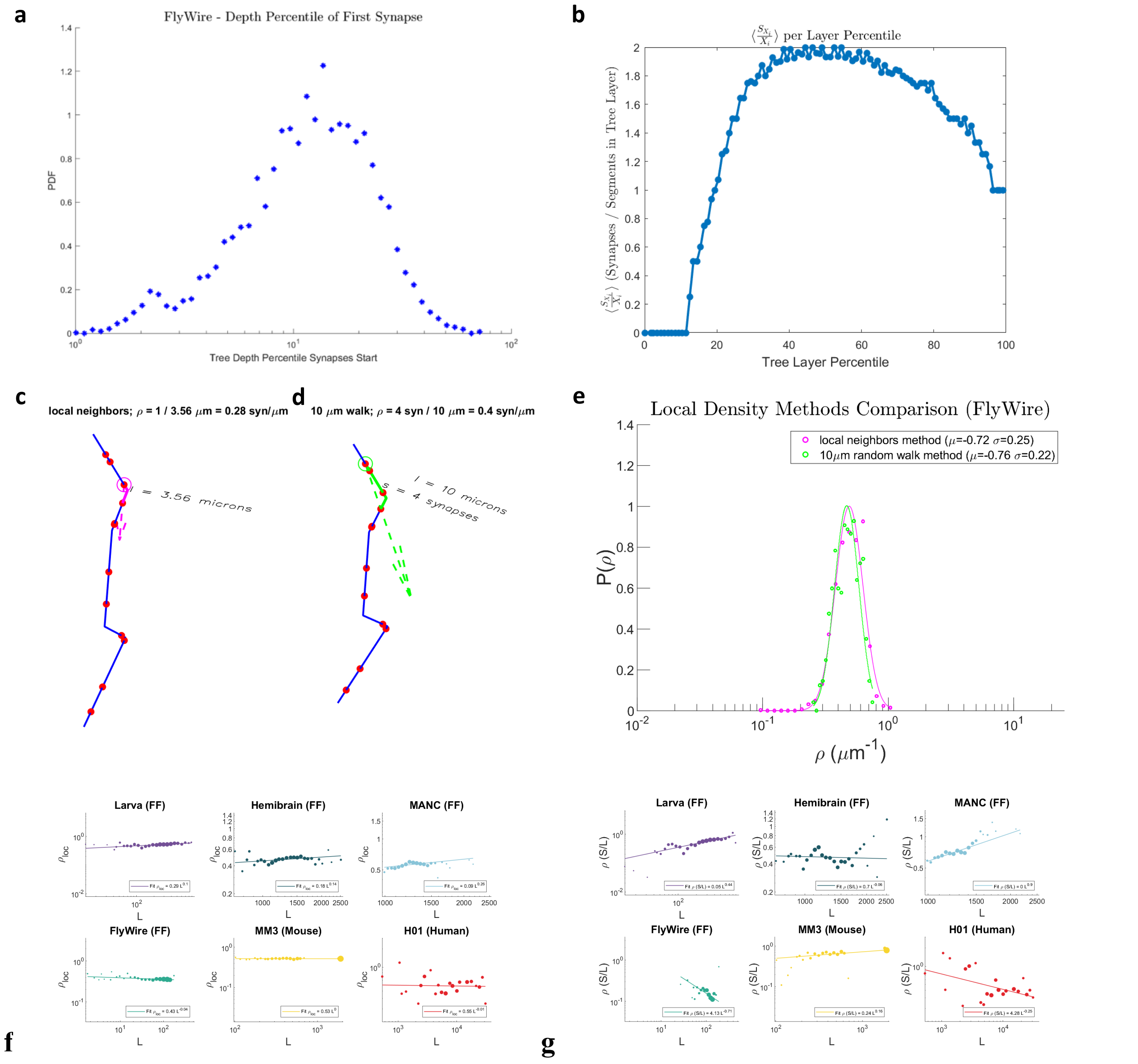

**Supplemental Figure 18: Local synapse densities for FlyWire.** To evaluate how uniformly synapses are distributed with respect to depth for FlyWire, we measured how far (in terms of tree layers) the first synapse in each neuron is from its soma. **(a)** We find that for most neurons, the first 10% of the neuron closest to the soma has no synapses. **(b)** We only start seeing synapses 14 tree layers from the soma on average. Y-axis quantifies the ratio of number of synapses to the number of total segments at a given tree layer depth. **(c)** The *neighbor method* estimates local synaptic density by picking a random synapse (arrow) and finding the distance to the next synapse on the same cell, in one direction. Blue lines indicate neuronal segments, red dots are synapses, and the magenta segment is the distance between two nearby synapses, in this case 14.38 μm. Note that the distance is not necessarily the nearest synapse, but simply the nearest synapse in one direction. **(d)** The *local synapses method* estimates synapse density by counting the total number of synapses within a certain distance, in one direction, of a randomly chosen synapse. Blue lines indicate neuronal segments, red dots are synapses, and the green region is the local neighborhood searched, in this case of size 10 μm, which contained 2 synapses. **(e)** Local synapse distributions in FlyWire according to the two methods in (c-d), both of which are well fit by a lognormal and offer comparable μs and $\sigma$s. **(f-g) Local synapse density versus naïve calculation.** We plotted both local density and total synapses over neuron length against neuron length, to determine if there are correlations. **(f)** Local synapse density remains approximately constant across all datasets, regardless of neuron length. **(g)** In contrast, the naïve synapse density calculation (total synapse count divided by neuron length) shows stronger correlations with length, demonstrating that this measure is unsuitable to argue proportionality between synapses and length.

Using local synapse density instead of total synapses per neuron length is crucial for accurately predicting synapse counts. When plotting synapse counts against neuron lengths (Fig. 3A), some datasets may appear superlinear or sublinear. This suggests that the naïve synapse density (synapses per unit length) is at least weakly correlated with length. However, local synapse density differs from this measure as it only considers regions where synapses are present. Consequently, we expect correlations between local synapse density and length to be significantly weaker. Indeed, when comparing both density measures against length, we find that local synapse density exhibits substantially lower correlations, indicating that synapse count is linearly proportional to length only when using a local measure of density.

As with the neuron length distributions, KS and other statistical tests indicate that the lognormal distribution is generally the best-fitting model among the candidate distributions for local synapse density (Fig. S19). For *C. elegans* we were not able to obtain reliable synapse locations, despite extended correspondence with the individuals who collected the data, thus we lack local synapse density information for this organism.

Crucially, neuron density does not seem to be significantly affected by brain size, fluctuating less than an order of magnitude between animals[38] (Fig. S20). This would indicate that differences in synapse number and degree between these animals will primarily be due to increased neuron size, which at a fixed density can then take more synapses. In fact, the lower synapse density in the human data is likely due to an undersampling of dendritic spines (and thus synapses) on the neurons (see Section 1.6).

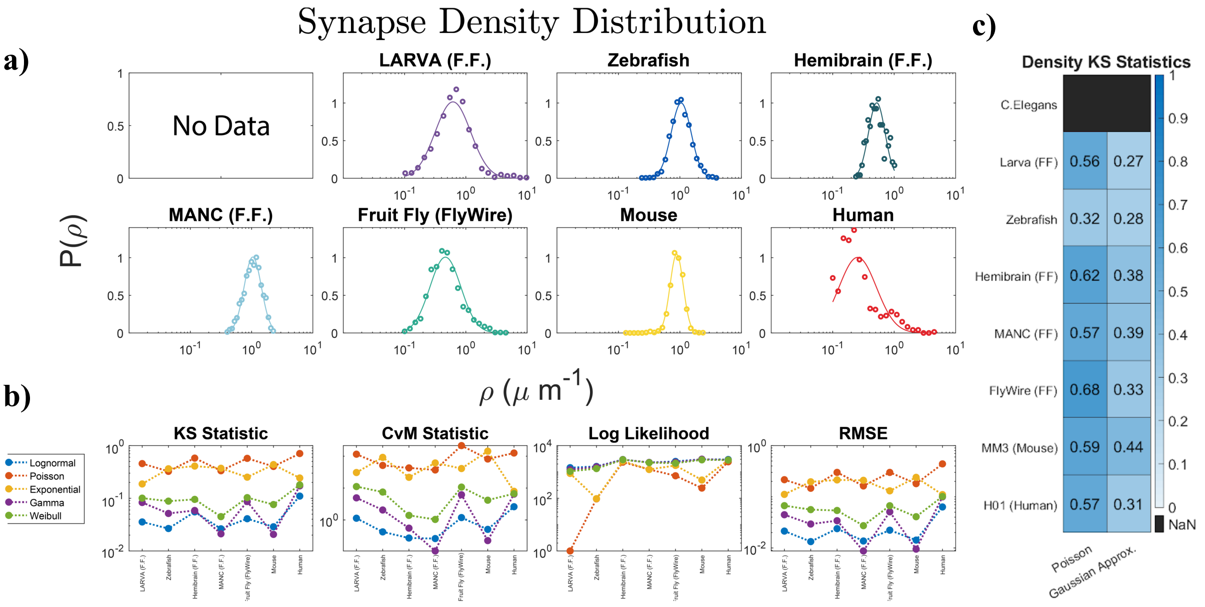

**Supplemental Figure 19: Neuron synapse density distributions across organisms.**
**Top two rows:** Synapse density distribution of all analyzed datasets, plotted on a log-x scale, with probability plotted on the (linear) y-axis. Scatter points show the empirical data and colored lines depict the best lognormal fit. For *C. elegans* (data not shown) we were not able to obtain reliable synapse locations thus we could not calculate the local synapse density for this organism.
**Bottom row:** For each dataset, we compare the goodness-of-fit using the Kolmogorov-Smirnov (KS) test, the Cramér-von Mises (CvM) test, the probabilistic log-likelihood measure, and the root mean square error (RMSE). In all cases, the y-axis displays the fit statistics, with a lower value indicating the better fit, and the x-axis indicates the dataset for which the fit was performed. Under log-likelihood, we find that Lognormal, Weibull and Gamma distributions overlap. In some cases we fit a discrete Poisson to a continuous variable. This is justified since for all datasets $\lambda \geq 20$, where Poisson closely approximates a normal distribution, making the KS test appropriate.
**c)** For sufficiently large parameter values (e.g., $\lambda \geq 20$), the Poisson closely approximates a Gaussian (continuous) distribution. We explicitly test this assumption, by comparing the Poisson KS statistics against the corresponding Gaussian approximation, with $\mu = \lambda$. We find that the Gaussian approximation is either only marginally better than or offers comparable fit quality to the Poisson.

| | Lognormal | Poisson | Exponential | Gamma | Weibull |
|---|---|---|---|---|---|
| KS | **0.046190099** | 0.4817983 | 0.32276125 | 0.070516819 | 0.098141306 |
| CVM | **0.912078295** | 95.52409694 | 56.87638863 | 2.930164775 | 6.092831512 |
| LL | **2398.003539** | 1011.044997 | 1405.289896 | 2323.41072 | 2201.93975 |
| RMSE | **0.024561048** | 0.249166008 | 0.173054226 | 0.039513347 | 0.059055469 |

**Supplemental Table 5: Mean Local Density Test Statistics.** For the KS, CvM, Log likelihood and RMSE tests visualized in the Figure above, we calculated the mean test statistic across all organisms, finding that the Lognormal emerges as the top-performing model across all four measures.

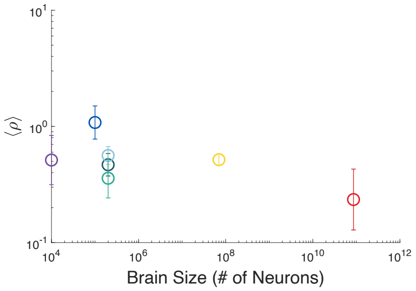

**Supplemental Figure 20: Correlation between brain size and neuron synapse density.** Comparison of average neuron synapse density with total neuron number (see Table S1) for all eight datasets (see Fig. S19 for color code).

#### Section 3.3: Node Strength (Synapse Count) Distribution *P(S)*

Synaptic connections provide the foundation for specific, directed information transfer between neurons. The number of synapses a neuron has thus defines its communication potential, both in the number of partners (degree) it can have (less than or equal to the number of synapses), as well as the connection strength it maintains with other cells (number of synapses made between a pair of neurons). For each studied system, we obtained the location of all synapses in the reconstructed volume, as well as their pre- and post-synaptic neuron identity. The exception lies in *C. elegans*, for which we only had access to the adjacency matrix, without synaptic locations. Both types of data structures were amenable to calculating the total number of synapses, both pre- and post-synaptic, a neuron has (Fig. S21, representing an undirected metric). We also plot the log-log scale of synapse number with brain size (Fig. S22). Note that human synapse numbers are expected to be higher, canonically averaging around 104, so the lower values observed here are likely due to an undersampled reconstruction (Section 1.6).

Among all the datasets we analyzed, the lognormal distribution provided the best fit to the synapse counts across all tests performed (Fig. S21). This is further supported by the visual inspection of the data, which reveals relatively clean distributions, likely due to the high abundance of synapses in the datasets. Log skewness is also minimal for most datasets, suggesting approximate symmetry in log space.

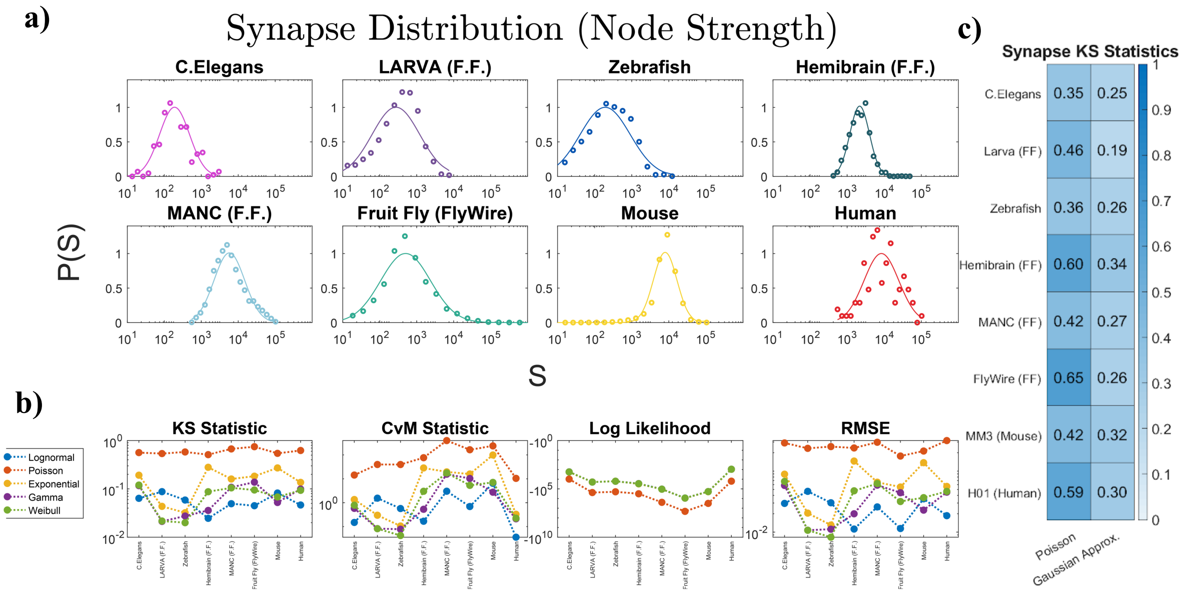

**Supplemental Figure 21: Neuron synapse number distributions across organisms.**
**a)** Synapse number distribution of all analyzed datasets, plotted on a log-x scale, with probability plotted on the (linear) y-axis. Scatter points show the empirical data and colored lines depict the best lognormal fit.
**b)** For each dataset, we compare the goodness-of-fit using the Kolmogorov-Smirnov (KS) test, the Cramér-von Mises (CvM) test, the probabilistic log-likelihood measure, and the root mean square error (RMSE). In all cases, the y-axis displays the fit statistics, with a lower value indicating the better fit, and the x-axis indicates the dataset for which the fit was performed. Under log-likelihood, we find that Lognormal, Exponential, Weibull and Gamma distributions overlap.
**c)** For sufficiently large parameter values (e.g., $\lambda \geq 20$, as is the case even for C. elegans), the Poisson closely approximates a Gaussian (continuous) distribution. We explicitly test this assumption, by comparing the Poisson KS statistics against the corresponding Gaussian approximation, with $\mu = \lambda$. We find that the Gaussian approximation is either only marginally better than or offers comparable fit quality to the Poisson.

| | Lognormal | Poisson | Exponential | Gamma | Weibull |
|---|---|---|---|---|---|
| KS | **0.05719429** | 0.597548701 | 0.162326865 | 0.074799572 | 0.076254639 |
| CVM | **1.58304116** | 122.7833739 | 20.63229489 | 3.772578301 | 4.265131829 |
| LL | **21049380.71** | 17751731.41 | 21045121.16 | 21046939.86 | 21047596.04 |
| RMSE | **0.026106656** | 0.252852621 | 0.072928277 | 0.033928473 | 0.038974035 |

**Supplemental Table 6: Mean Synapse Number Test Statistics.** For the KS, CvM, Log likelihood and RMSE tests visualized in the Figure above, we calculated the mean test statistic across all organisms, finding that the Lognormal emerges as the top-performing model across all four measures.

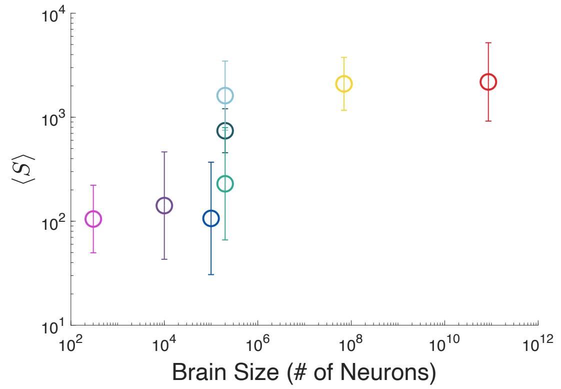

**Supplemental Figure 22: Correlation between brain size and neuron synapse number.** Comparison of average neuron synapse number with total neuron number (see Table S1) for all eight datasets (see Fig. S21 for color code). It has not escaped our attention that the data is consistent with a potential power law scaling between <S> and N, suggesting <S>~$N^{\delta}$, whose origins and consequences need to be still unveiled.

**Supplemental Table 7: Mean Log Synapses Prediction**

| | Measured | | | Predicted |
|---|---|---|---|---|
| | $\mu_L$ | $\mu_D$ | $\mu_S$ | $\mu_{S^*} = \mu_L + \mu_D$ |
| Human | 8.96 ± 0.83 | -0.85 ± 0.60 | 7.69 ± 0.87 | 8.11 ± 0.83 |
| Mouse | 7.94 ± 0.39 | -0.16 ± 0.25 | 7.78 ± 0.51 | 7.78 ± 0.64 |
| FlyWire (F.F.) | 6.54 ± 0.85 | -1.15 ± 0.83 | 5.43 ± 1.24 | 5.39 ± 1.27 |
| MANC (F.F.) | 7.05 ± 1.06 | -0.18 ± 0.48 | 6.86 ± 1.15 | 6.86 ± 1.15 |
| Hemibrain (F.F.) | 6.93 ± 1.01 | -0.69 ± 0.46 | 6.22 ± 0.92 | 6.24 ± 0.92 |
| F.F. Larva | 5.55 ± 0.58 | -0.60 ± 0.90 | 4.95 ± 1.20 | 4.95 ± 1.20 |
| Zebrafish | 6.01 ± 0.83 | -0.46 ± 0.39 | 5.96 ± 0.92 | 5.96 ± 0.69 |
| *C. elegans* | 5.18 ± 0.99 | -0.53 ± 1.24 | 4.65 ± 0.74 | 4.65 ± 0.74 |

**Supplemental Table 8: Standard Deviation Log Synapses Prediction**

| | Measured | | | | Predicted |
|---|---|---|---|---|---|
| | $\sigma_L$ | $\sigma_D$ | $\sigma_S$ | $corr(L, D)$ | $\sigma_{S^*} = \sigma_L^2 + \sigma_D^2$ $+2\sigma_L\sigma_D corr(L, D)$ |
| Human | 0.83 | 0.60 | 0.87 | -0.85 | 0.90 |
| Mouse | 0.39 | 0.25 | 0.51 | 0.60 | 0.64 |
| FlyWire (F.F.) | 0.85 | 0.83 | 1.24 | 0.35 | 1.27 |
| MANC (F.F.) | 1.06 | 0.48 | 1.15 | -0.07 | 0.92 |
| Hemibrain (F.F.) | 1.01 | 0.46 | 0.92 | -1.01 | 0.92 |
| F.F. Larva | 0.58 | 0.90 | 1.20 | 0.69 | 1.20 |
| Zebrafish | 0.83 | 0.39 | 0.92 | 0.02 | 0.69 |
| *C. elegans* | 0.99 | 1.24 | 0.74 | -1.84 | 0.74 |

#### Section 3.4: Neuron Degree Distribution *P(k)*

The number of unique neurons a selected neuron *i* connects to defines its degree $k_i$, the most fundamental property of network analysis. In neuroscience, the degree of a cell provides a quantification of its capacity to integrate signals from and disperse to diverse sources; the higher the degree, the more information input/output a cell has. The degree of a neuron can be calculated by taking the list of all synapses it makes, and counting all unique partners in that list, a process we performed for all neurons in all 8 datasets (Fig. S23). The low mouse degrees are likely due to the truncated volume, and a low number of reconstructed axons, which often exit the volume, offering an incomplete count of the full cells' degrees (see SI 1.5). The human degrees are similarly lower due to an undersampled reconstruction, as well as missing axons (SI 1.6).

In studying the neuron degrees, we found that this metric is generally well approximated by a lognormal distribution (Fig. S23), however KS and other tests indicate that the Weibull and gamma distributions can also provide comparable fits in terms of KS parameter (Fig. S23). Additionally, neuron degrees tends to exhibit a negative log skewness across all datasets (see SI 3.5).

A neuron's degree is determined by its synapse number, as the highest degree that a cell may have is the total number of synapses (connections) it makes. Yet, the actual degree of a cell is in most cases considerably less than its synapse number, as neurons may make multiple synapses with any given cell, thereby providing a stronger information propagation to that partner. Thus, synapse number and neuron degree provide complimentary, but distinct, perspectives on cell identity.

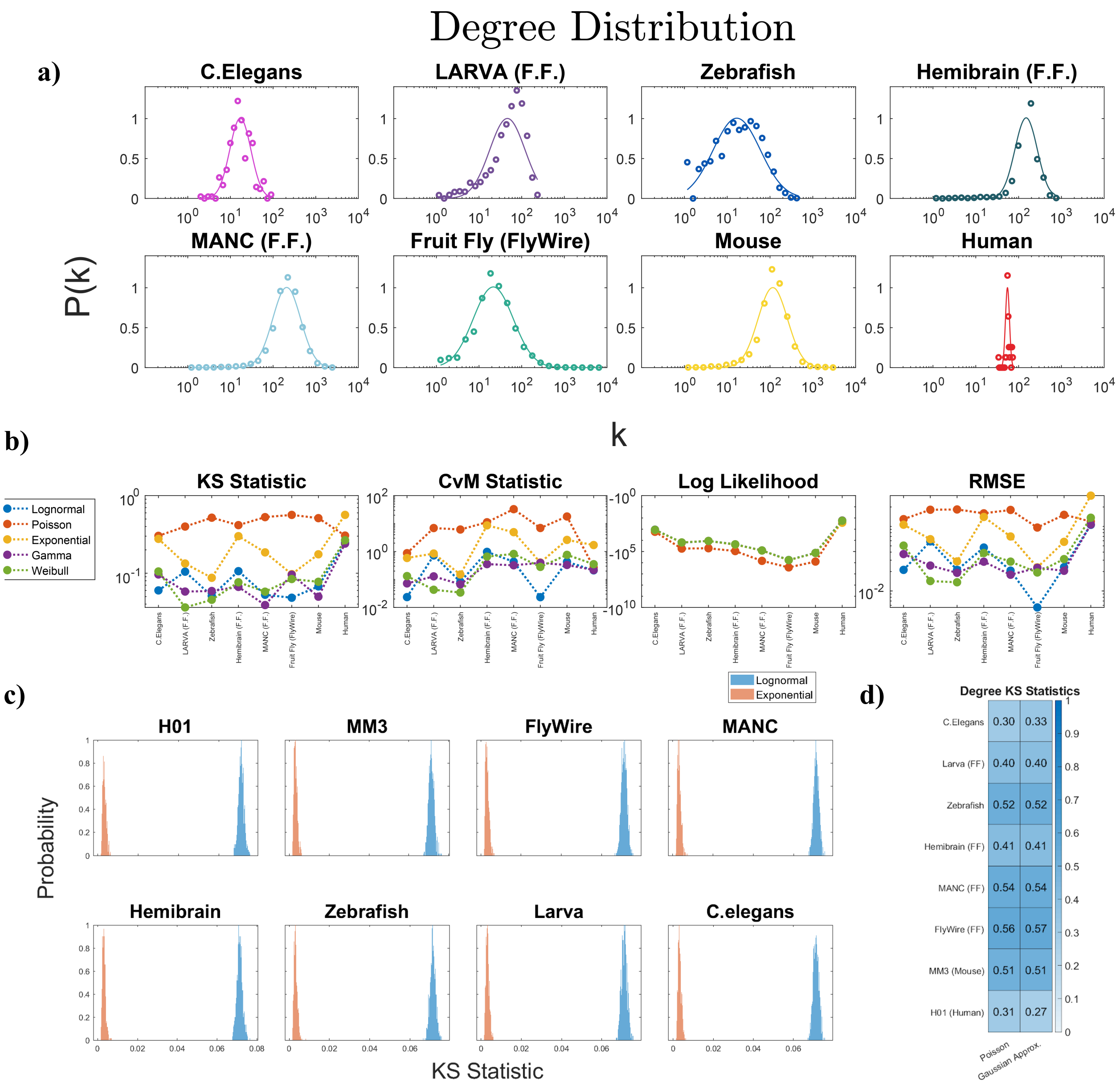

**Supplemental Figure 23: Neuron degree distributions across organisms.**

**a)** Degree distribution of all analyzed datasets, plotted on a log-x scale, with probability plotted on the (linear) y-axis. Scatter points show the empirical data and colored lines depict the best lognormal fit.

**b)** For each dataset, we compare the goodness-of-fit using the Kolmogorov-Smirnov (KS) test, the Cramér-von Mises (CvM) test, the probabilistic log-likelihood measure, and the root mean square error (RMSE). In all cases, the y-axis displays the fit statistics, with a lower value indicating the better fit, and the x-axis indicates the dataset for which the fit was performed. Under log-likelihood, we find that the Lognormal, Exponential, Weibull and the Gamma distributions overlap.

**c)** To illustrate the comparability of one- versus two-parameter distributions, we generated 1000 sets of 50,000 exponential random variables and computed KS statistics for exponential (one-parameter) and lognormal (two-parameter) fits, with the exponential parameter λ set to the mean of each dataset. Across all eight connectomes, exponential fits consistently yielded lower KS statistics than lognormal fits, despite the latter having an additional free parameter. These results

demonstrate that the superior fits observed for two-parameter distributions in our manuscript are not simply due to parameter count, but rather reflect properties of the underlying data.
**d)** For sufficiently large parameter values (e.g., $\lambda \geq 20$, as is the case even for C. elegans), the Poisson closely approximates a Gaussian (continuous) distribution. We explicitly test this assumption, by comparing the Poisson KS statistics against the corresponding Gaussian approximation, with $\mu = \lambda$. We find that the Gaussian approximation is either only marginally better than or offers comparable fit quality to the Poisson.

| | Lognormal | Poisson | Exponential | Gamma | Weibull |
|---|---|---|---|---|---|
| KS | 0.092417388 | 0.443794695 | 0.227717906 | **0.088147799** | 0.093718868 |
| CVM | 0.362732127 | 10.37713889 | 2.481804507 | **0.236114822** | 0.385806336 |
| LL | **2522873.264** | 2089763.331 | 2520826.28 | 2521555.408 | 2521582.776 |
| RMSE | 0.0379467 | 0.149970532 | 0.098618505 | **0.034762474** | 0.041557381 |

**Supplemental Table 9: Mean Neuronal Degree Test Statistics.** For the KS, CvM, Log likelihood and RMSE tests visualized in the Figure above, we find the mean test statistic across all organisms. Here, the lognormal closely rivals the gamma; the relative difference between them is a few percent across most measures. Yet, as we discuss in SI Sect 2.5, while the gamma and lognormal distributions are often hard to differentiate based on statistics alone, there is no theoretical mechanism that would substantiate a gamma distribution in a network environment. This leaves only the lognormal as a valid model, as that is the only one which offers a statistically acceptable fit, and we have theoretical arguments for its relevance to the connectome.

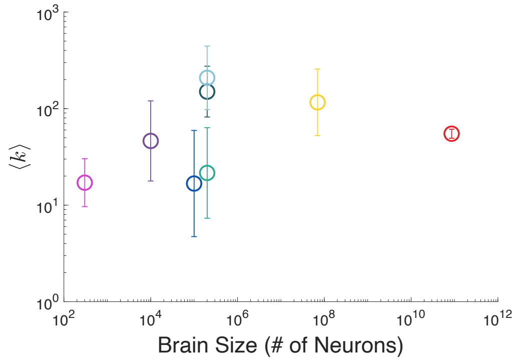

**Supplemental Figure 24: Correlation between brain size and neuron degree.** Comparison of average neuron degree with total neuron number (see Table S1) for all eight datasets (see Fig. S23 for color code).

**Supplemental Table 10: Mean Log Degree Prediction**

| | Measured | | | Predicted |
|---|---|---|---|---|
| | $\mu_S$ | $\mu_{\frac{k}{S}}$ | $\mu_k$ | $\mu_{k^*} = \mu_S + \mu_{\frac{k}{S}}$ |
| Human | 7.69 ± 0.87 | -4.03 ± 0.87 | 4.01 ± 0.12 | 3.66 ± 0.12 |
| Mouse | 7.69 ± 0.64 | -3.57 ± 0.62 | 4.01 ± 0.81 | 4.12 ± 0.83 |
| FlyWire (F.F.) | 5.43 ± 1.24 | -3.22 ± 0.35 | 2.44 ± 1.08 | 2.21 ± 1.08 |
| MANC (F.F.) | 6.86 ± 1.15 | -2.03 ± 0.99 | 5.11 ± 0.85 | 4.84 ± 0.87 |
| Hemibrain (F.F.) | 6.22 ± 0.92 | -1.84 ± 0.62 | 4.79 ± 0.76 | 4.70 ± 0.76 |
| F.F. Larva | 4.95 ± 1.20 | -1.84 ± 0.44 | 3.27 ± 0.97 | 3.11 ± 0.99 |
| Zebrafish | 5.96 ± 0.92 | -1.15 ± 0.94 | 2.33 ± 1.13 | 2.51 ± 1.08 |
| *C. elegans* | 4.65 ± 0.74 | -1.93 ± 0.44 | 2.83 ± 0.58 | 2.72 ± 0.55 |

**Supplemental Table 11: Standard Deviation Log Degree Prediction**

| | Measured | | | | Predicted |
|---|---|---|---|---|---|
| | $\sigma_S$ | $\sigma_{\frac{k}{S}}$ | $\sigma_k$ | $corr\left(S, \frac{K}{S}\right)$ | $\sigma_{k^*} = \sigma_S^2 + \left(\sigma_{k/S}\right)^2 + 2\sigma_{\frac{k}{S}}\sigma_S corr\left(S, \frac{k}{S}\right)$ |
| Human | 0.87 | 0.87 | 0.12 | -2.28 | 0.12 |
| Mouse | 0.64 | 0.62 | 0.81 | -0.39 | 0.83 |
| FlyWire (F.F.) | 1.24 | 0.35 | 1.08 | -1.29 | 1.08 |
| MANC (F.F.) | 1.15 | 0.99 | 0.85 | -1.57 | 0.87 |
| Hemibrain (F.F.) | 0.92 | 0.62 | 0.76 | -1.29 | 0.76 |
| F.F. Larva | 1.20 | 0.44 | 0.97 | -1.45 | 0.99 |
| Zebrafish | 0.92 | 0.94 | 1.13 | -0.76 | 1.08 |
| *C. elegans* | 0.74 | 0.44 | 0.58 | -1.50 | 0.55 |

#### Section 3.5: Skewness

Skewness measures the asymmetry of a probability distribution around its mean. For a dataset $\{x_1, x_2, \dots, x_n\}$ with sample mean $\bar{x}$ and sample standard deviation $s$, the sample skewness $\gamma$ is given by:

$$\gamma = \frac{\frac{1}{n}\sum_{i=1}^{n}(x_i - \bar{x})^2}{s^3} \tag{26}$$

A positive skewness value indicates a distribution with a longer right tail (e.g., the lognormal distribution), whereas a negative skewness value indicates a longer left tail. If skewness is close to zero, the distribution is approximately symmetric (e.g., the normal distribution).

Skewness can also be evaluated for log-transformed data, referred to here as *log skewness*. The lognormal distribution is symmetric in log-space but asymmetric in linear space, leading to distinct values for skewness and log skewness.

For log skewness, only a few distributions are expected to approximate symmetry, notably the lognormal, gamma, and Weibull distributions. We compare the log skewness for these distributions in the table below:

| Distribution | Log Skewness | Explanation |
|---|---|---|
| Lognormal | Approximately symmetric (zero) | Taking the logarithm transforms a lognormal distribution into a normal distribution, removing skewness in the log space. |
| Gamma | Positive (for k<3), ~Symmetric (for k≥3) | For smaller shape parameters (k), the distribution remains right-skewed after log transformation but approaches symmetry as k increases. |
| Weibull | Depends on shape parameter k:<br>- ~Symmetric (k≈1)<br>- Negative (k>1) | - A Weibull distribution with k≈1 behaves similarly to an exponential distribution, and its log transformation is symmetric.<br>- For k>1, the log-transformed Weibull becomes left-skewed due to compression of its long right tail. |

All observed distributions *P(k), P(S), P(L)* and *P(ρ)* display a small negative skewness, indicating that these quantities have a longer left tail than a right tail. As we discuss below, part of this skewness can be attributed to data incompleteness. Specifically, when neurons are only partially reconstructed, the observed neurons will be smaller than expected based on their true *P(L)* (incomplete *L*), thus having fewer synapses *S* and lower degrees *k*. As a result, a neuron that would be at the center or right tail of the distribution moves to the left. This incompleteness is reflected in the log-skewness. To test our hypothesis that a source of the observed skewness is data incompleteness, we compare the skewness of the eight datasets, finding that the more complete datasets, like the *C. elegans* and FlyWire, have some of the lowest skewness, close to zero across multiple distributions (*P(k), P(S), P(L)* and *P(ρ))*. In contrast, the highly incomplete Fly Hemibrain, where half the brain and its connections are missing, display a strong skewness (Fig. S25a-d).

To validate the hypothesis that the incomplete reconstruction leads to skewness, we focused on the FlyWire dataset, which has a high completeness and a log degree skewness close to zero. We used simulations to test the impact of incomplete mapping by defining a volume box of linear size L', and only considering cells that were fully contained within the restricted volume (Fig. S25e). Consistent with our hypothesis, we find that log-skewness for all variables of interest, namely *L*, *k* and *S*, becomes more negative as the reconstructed volume decreases (Fig. S25f-h).

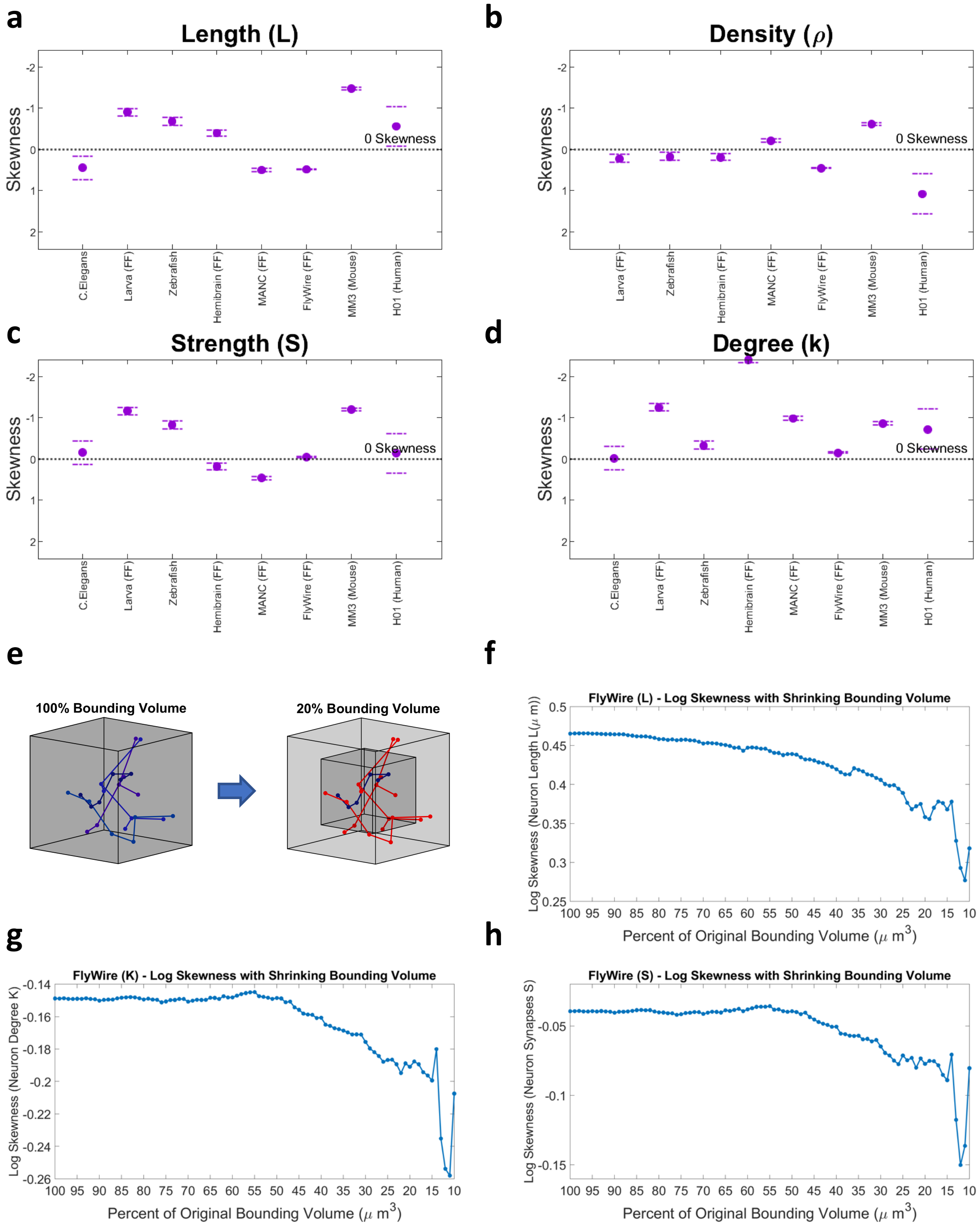

**Supplemental Figure 25: Log-skewness of the lognormal fit across datasets and measures. (a-d)** In all panels, we show the log-skewness measured (y-axis) when fitting a lognormal to the empirical distributions. The dashed line indicates 0 skewness, and higher values (top region) are more negatively skewed. Error bars indicate confidence intervals calculated through standard error (1.95 * standard error of skewness). **(e)** Utilizing the large, and highly complete, dataset of the FlyWire, we relied on simulations to define a volume box of size L', and we reduced the L' and only considered cells that were fully contained within the restricted volume. **(f-h)** We compare the log-skewness of L, K and S (y-axis, lower is more negatively skewed), with the relative amount of volume remaining (x-axis, with values further to the right are further restricted in volume).

### Section 4: Theoretical Results

In this section, we investigate the mechanistic origin of the empirically observed lognormal distribution that characterizes the key variables describing the connectome. As mentioned in the main manuscript, while lognormal distributions have occasionally been employed to fit fat-tailed degree distributions, these efforts have remained purely statistical exercises. This is due to the lack of a mechanistic model and an underlying theoretical framework that would explain the emergence of a lognormal *P(k)* within the network context (SI 2.9).

Here, we explore the implications of the connectome being a physical network, where the fundamental unit—serving simultaneously as both node and link—is the neuron. Mathematically, a neuron can be modeled as an incomplete binary tree. Consequently, we utilize the well-established Galton-Watson (GW) process to describe the growth of individual neurons and analytically demonstrate that, under a minimal set of assumptions consistent with biological data, the size distribution of these binary trees is expected to follow a lognormal distribution.

We provide both analytical and numerical evidence to support the lognormal nature of the resulting tree size distribution *P(L)* within the biologically relevant regimes of the connectome. This approach not only offers a mechanistic understanding of the observed distributions but also bridges the gap between statistical observations and the underlying biological processes governing network formation.

#### Section 4.1: Modeling Neuron Growth

As stated in the introduction of Section 4, the goal of utilizing a Galton-Watson process lies in developing a minimal, tractable model for understanding the observed distributions, rather than replicating the exact 3D morphology of neurons. The crucial assumption of this model lies in the fact that axons and dendrites form predominantly binary trees, which is commonly accepted in the field[39]. The main exception to this rule would lie in soma, or cell body, from which multiple neurites can emerge, however from there generally branching remains binary.

There is a rich history of studying branching statistics aimed at capturing their characteristic properties[40]. These studies can be grouped into two classes: many studies focus on the branching structure and not their geometry (structural statistics)[41–43], while other studies aim to capture aspects of their geometry (geometrical statistics)[44–46]. Most of the early studies did not have access to the fine resolution of EM connectomics, and therefore relied on lower-quality morphologies. Nevertheless, the branching structures were already found to be compatible with a terminal growth hypothesis[47,48] and associated dendrograms were shown to be well-described by simple branching processes[48].

Most of these studies focused on classifying dendritic morphologies, which were more readily reconstructable, and utilized descriptive morphometrics, which are sufficient for categorizing mature cell types and inferring functional roles[49]. However, analyzing only the developed cells poses challenges in uncovering general principles of structure-function relationships due to the inherent variability caused by distinct growth environments. To address this, many researchers turned to generating synthetic morphologies *in silico* based on specific principles or mechanisms, as we have done with the Galton-Watson process. By comparing these synthetic cells with real counterparts, researchers can infer the validity of the underlying principles[50,51].

Generative models fall into three categories:

1. **Reconstruction Models** are the simplest and focus on replicating the static shape of mature dendritic trees. These models quantify various morphometrics from real cells and fit statistical distributions to the data. By making assumptions about correlations between morphological variables, algorithms generate synthetic morphologies that mimic real dendrites[46,52–54]. While limited to static datasets, these models offer valuable insights into dendritic morphology at specific developmental stages.

2. **Growth Models** aim to capture the spatiotemporal dynamics of dendritogenesis by modeling how morphometrics evolve over time[51,55,56]. These models often incorporate effects such as branch self-avoidance[57] and interactions between dendrites[58]. Functional assumptions, like wiring minimization[50] or connectivity maximization[59], enhance their ability to generate realistic morphologies while explaining structure-function relationships.

By requiring fewer parameters, growth models achieve a balance between simplicity and explanatory power.

3. **Biophysical Models** provide the most detailed and mechanistic approach to dendrite growth, simulating the role of biophysical parameters like production rates, local interactions, and molecular transport[60,61]. These models incorporate intracellular and extracellular factors to derive arborization patterns. Despite their detail, biophysical models face challenges due to the limited availability of high-resolution data and the computational demands of simulating complex interactions[62]. Nonetheless, they hold significant promise for unraveling the intricate mechanisms of dendritic growth.

Our work has a novel perspective in its utilization of the high-quality comparative connectomics datasets recently made availability. We note for the sake of analytical simplicity, in the Galton-Watson process simulated below we utilized a branching process with segments of fixed length, which diverges from standard assumptions of variable segment length often utilized in the above models. However, this assumption can be easily relaxed. In turning to the data, we find empirically that segment lengths across all organisms are also lognormal (Fig. S26). Yet, when we rescale all segments within a neuron to the organism's mean segment length, thereby simulating our model's fixed segment length assumption, we find that the rescaled neurons are still lognormally distributed (Fig. S27, black points and lines), and generally do not deviate significantly from the empirical data (Fig. S27, colored points and lines).

A major limitation of our utilized connectomics datasets is the uneven biology of axon and dendrite identity across the eight datasets— in C. elegans and Drosophila connectomes, axon/dendrite annotations exist, but many neurons are multipolar, with pre- and post-synaptic sites intermixed on the same neurite—so the classical dichotomy is blurred. In zebrafish, mouse, and human datasets polarity is clearer, yet the currently existing reconstructions cover only local regions, so full axonal trajectories are limited and a compartment-level growth model would be under-constrained. Given these limitations, we chose not to include additional structural considerations in our current growth model.

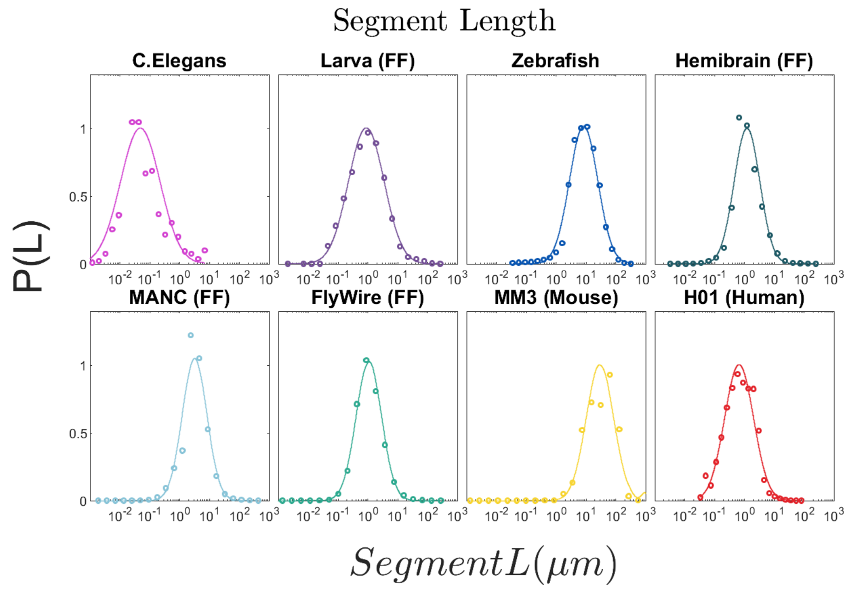

**Supplemental Figure 26: Segment Length Distributions.** Segment length distribution of all analyzed datasets, plotted on a log-x scale, with probability plotted on the (linear) y-axis. Segments are defined as stretches between any two points of branching on a neuron. Scatter points show the empirical data and colored lines depict the best lognormal fit.

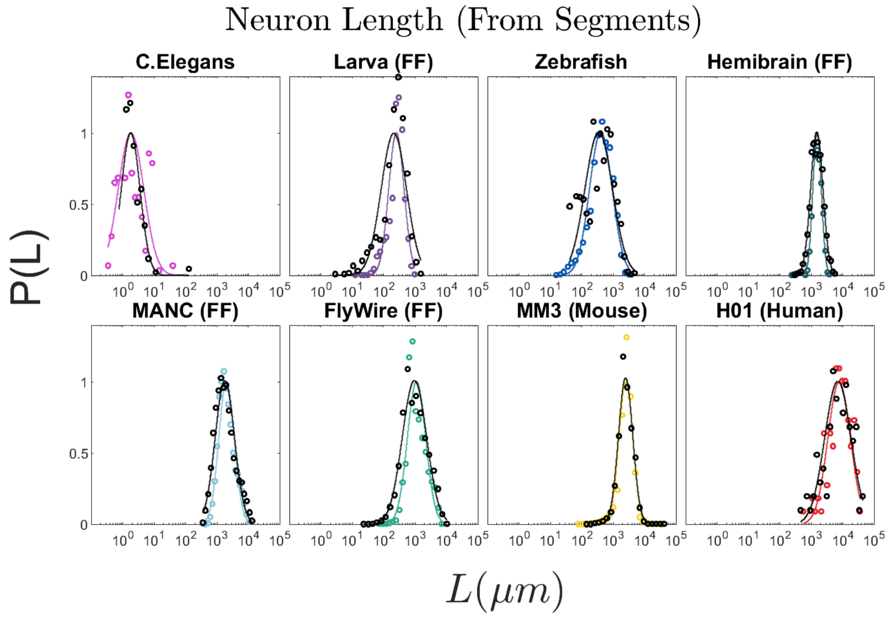

**Supplemental Figure 27: Neuron lengths do not significantly change when rescaled to have constant segment length.** Neuron length distribution of all analyzed datasets, plotted on a log-x scale, with probability plotted on the (linear) y-axis. Colored scatter points show the empirical data and colored lines depict the best lognormal fit. Black points show the datasets when all segments within a neuron to the organism's mean segment length, thereby simulating our model's fixed segment length assumption, and the black line depicts the best lognormal fit to this rescaled data.

#### Section 4.2: The Galton-Watson Process

The branching structure of each neuron can be mathematically represented as a random binary tree—where the root corresponds to the soma or an initial cluster of segments, and each node, representing a neuronal segment, can give rise to at most two child branches (Fig. 4a in the manuscript). To model this process, we use a Galton-Watson (GW) framework, a mathematical model originally developed to study population dynamics across generations[63]. While this approach captures the stochastic nature of neuronal branching, it does not account for the distribution of individual segment lengths.

The Galton-Watson model[64], first introduced in the 19th century to analyze the extinction of family names, has since found widespread applications in diverse fields, including biology, computer science, and the study of branching systems. In its most basic form, a GW process begins with an initial population $X_0$ of individuals or nodes, typically set to $X_0 = 1$ for simplicity. As generations unfold, each node gives rise to a random number of offspring, leading to an evolving tree-like structure that captures the stochastic nature of neuronal growth (Fig. 4a-b in the manuscript).

At generation *i,* the number of elements $X_i$ follows

$$X_i = \sum_{m=1}^{X_{i-1}} Z_j^{i-1} \tag{27}$$

where $Z_j^{i-1}$ is a random variable capturing the number of descendants from a single element of the population. For neurons, $X_i$ indicates the number of segments at distance *i* from the soma.

A standard GW process assumes $\{Z_j^{i-1}\}\ \forall\,(j,m)$ to be identical and independently distributed. This means that each $\{Z_j^{i-1}\}$ follows the same *offspring distribution* $p(m)$, which represents the likelihood that a given neuronal segment branches into *m* subsegments.

Here, we simplify the offspring distribution by allowing a given neuronal segment to branch into two segments with *probability of survival* $p_s$, or to die out with probability $(1 - p_s)$. The offspring distribution is then a rescaled Bernoulli variable,

$$p(m) = p_s\delta(m-2) + (1 - p_s)\delta(m), \tag{28}$$

where δ is the Kronecker delta function, ensuring that the segment either produces exactly two offspring or none. The expected number of offspring per segment, or the mean of the distribution, is given by:

$$\mu = 2p_s\,. \tag{29}$$

Similarly, the variance of the distribution, which quantifies the variability in the number of offspring per segment, is expressed as:

$$\sigma^2 = 4p_s(1 - p_s) = 2\mu - \mu^2. \tag{30}$$

For simplicity, we assume that each neuronal segment can generate at most two subsegments, as this is the most common branching pattern observed in real data. Although instances of segments branching into three or more subsegments do occur, they are not considered in this model. However, it is straightforward to generalize the model to higher-order branches[46].

At generation $i$, the probability $P(X_i)$ of observing $X_i$ segments at distance $i$ from the soma will follow a Binomial probability distribution on even numbers

$$P(X_i = y) = \binom{X_{i-1}}{\frac{y}{2}} p_s^{\frac{y}{2}} (1 - p_s)^{X_{i-1}-\frac{y}{2}}, \tag{31}$$

with average

$$\langle X_i \rangle = X_{i-1} 2p_s = X_{i-1}\mu, \tag{32}$$

and variance

$$\sigma^2[X_i] = X_{i-1} 4p_s(1 - p_s) = X_{i-1}\sigma^2. \tag{33}$$

To characterize $P(X_i)$, which depends only on Eqs. (29) and (30), we next introduce the probability-generating function (PGF) — a mathematical tool that encodes the probabilities of different outcomes in a compact form, making it easier to analyze random processes. The PGF is expressed as a power series, $\mathrm{G(z) = p_0 + p_1 z + p_2 z^2 + p_3 z^3 +}$ ..., where each coefficient $p_m$ represents the probability of an event occurring *m* times, and the variable *z* is a placeholder that helps manipulate and extract probability-related information. By differentiating and evaluating the function at specific values of *z*, one can determine key statistical properties such as expected values and variances.

For the offspring probability distribution, the PGF is defined as

$$G(z) = (1 - p_s) + p_s z^2 . \tag{34}$$

For $P(X_i)$ , the PGF follows

$$F_i(z) = \sum_{y=0}^{\infty} P(X_i = y) z^y . \tag{35}$$

The properties of the compound probability distributions allow us to reformulate Eq. (35) as

$$F_i(z) = G\left(G(\ldots G(z))\right) = G\left(F_{i-1}(z)\right) \tag{36}$$

where $G(z)$ is compounded *i* times to obtain $F_i(z)$.
Leveraging Eqs. (30)-(32), we obtain the average and variance of $P(X_i)$

$$\langle X_i \rangle = \langle X_{i-1} \rangle \mu = \mu^i \tag{37}$$

$$\sigma^2[X_i] = \sigma^2 \mu^{i-1} \frac{(\mu^i - 1)}{(\mu - 1)} = \mu^i(\mu^i - 1)\frac{2-\mu}{\mu - 1} \qquad p_s \neq 0.5$$
$$\sigma^2[X_i] = i\sigma^2 = i \qquad p_s = 0.5 \tag{38}$$

When the initial population $X_0 > 1$, corresponding to $X_0$ sub-branches emerging from the same root—such as a soma giving rise to multiple dendrites—Eqs. (37) and (38) are multiplied by the factor $X_0$, reflecting the assumption that these branching processes are independent and identically distributed. Increasing $X_0$ improves the accuracy of the continuous approximation for this inherently discrete stochastic process, much like how a Binomial distribution converges to a Gaussian as the number of trials *n* increases.

Note that, depending on the parameters of the GW process, a tree (neuron) has a finite probability of growing indefinitely or stopping its growth. Hence, a key variable is the probability that the random tree will eventually stop growing, known as *the probability of extinction* $p_E$. This probability is determined by the offspring distribution $p(m)$, and is given by

$$p_E = G(p_E) = (1 - p_s) + p_s(p_E)^2 . \tag{39}$$

The smallest root of this second-order equation is within 0 and 1 for varying $p_s$, and follows

$$p_E = \frac{1 - \sqrt{1 - 4p_s(1 - p_s)}}{2p_s} = \frac{1 - \sqrt{1 - \sigma^2}}{\mu}. \tag{40}$$

Eqs. (37)-(38)-(40) predict the existence of two distinct regimes, along with the critical $p_s$ variable that separates them. In the *subcritical regime*, where $p_s < 0.5$, the neuron will ultimately stop growing (branching), with probability of extinction $p_E = 1$. When $p_s > 0.5$, the growth process becomes *supercritical*, with $p_E = \frac{1-p_s}{p_s} < 1$ and approaching 0 for $p_s \rightarrow 1$, implying that the neuron has a finite probability of growing indefinitely.

#### Section 4.3: Introducing Variability in the Galton-Watson Process

For simplicity, we have thus far assumed that $p_s$ remains constant across all neuronal layers. However, an examination of Main Text Figs. 4d-e and 5d,h suggests otherwise:

(1) **Stochastically:** For individual neurons the value of $p_s$ fluctuates from generation to generation, particularly during the early stages of neuronal growth.

(2) **Layer dependence:** The probability of branching $p_s$ is not uniform across neurons; instead, it varies due to multiple factors, including cell type, geometry, and the surrounding environment.

These sources of variability can collectively be regarded as biological noise, introducing an additional layer of complexity to the growth process.

Layer Dependence of $p_s$

To incorporate (1) and (2) into our model, we first generalize $p_s$ to a family of random variables $\{p_s^i\}$ that depend on the layer number $i$, measured as the distance from the soma. To achieve this, we first extend Eq. (36) to account for a set of probabilities $\{p_s^i\}$, where each generation $i$ has a distinct branching probability. This generalization leads to the recursive formulation:

$$F_i(s) = G_1\left(G_2\left(\dots G_{i-1}(z)\right)\right) = F_{i-1}\left(G_i(z)\right), \tag{41}$$

which systematically integrates layer-dependent variability into the branching process.

Eqs. (37)-(38) generalize to

$$\langle X_i|\{p_s^i\}\rangle = \langle X_{i-1}|\{p_s^{i-1}\}\rangle\mu_i = \prod_{k=1}^{i}\mu_k \ , \tag{42}$$

$$\begin{aligned}\sigma^2\left[X_i|\{p_s^i\}\right] &= \prod_{k=1}^{i}\mu_k - \left(\prod_{k=1}^{i}\mu_k\right)^2 + \sum_{k=1}^{i}(\sigma_k^2 - \mu_k + (\mu_k)^2)\prod_{j=1}^{k-1}\mu_j\prod_{j=k+1}^{i}{\mu_j}^2 \\ &= \prod_{k=1}^{i}\mu_k - \left(\prod_{k=1}^{i}\mu_k\right)^2 + \sum_{k=1}^{i}\mu_k\prod_{j=1}^{k-1}\mu_j\prod_{j=k+1}^{i}{\mu_j}^2 \ ,\end{aligned} \tag{43}$$

which describe the expected number of segments and variance at layer $i$, given a set of measured $\{p_s^i\}$.

These equations characterize a scenario in which $\{p_s^i\}$ changes deterministically, and each generation's mean and variance are influenced by all previous generations. Since $X_i$ is derived from the repeated summation of random offspring numbers, and given that the offspring distribution is not heavy-tailed (as described in Eq. 28), this setup satisfies the conditions of the Central Limit Theorem (CLT). The CLT states that the sum of a large number of independent or weakly dependent random variables with finite variance converges to a normal distribution. Consequently, in the case of deterministically varying $\{p_s^i\}$, while the expected value $\langle X_i|\{p_s^i\}\rangle$ and variance $\sigma^2[X_i|\{p_s^i\}]$ grow exponentially, as $i$ increases, the probability distribution $P(X_i)$ approaches a Gaussian form $P(X_i) \approx \mathcal{N}(\langle X_i|\{p_s^i\}\rangle, \sigma^2[X_i|\{p_s^i\}])$. This implies that the overall distribution of the number $X_i$ of segments at layer $i$ becomes increasingly Gaussian for large $i$, with generation-specific mean and variance.

Stochastic $p_s$

What if the branching probabilities $\{p_s^i\}$ do not change just deterministically over the generations? This implies introducing noise into the model, meaning that even if each neuron follows the same underlying GW process, the observed sets $\{p_s^i\}$ will vary from neuron to neuron, beyond mere measurement error. As a result, the generalized GW process governing these observations does not have a fixed branching probability at each generation $i$; instead, $p_s^i$ is drawn from a stochastic distribution with an average $\langle p_s^i\rangle$ and variance $\sigma^2[p_s^i]$. A natural and practical choice for modeling $p_s^i$ is the Beta distribution, commonly used in Bayesian inference to represent probabilities under uncertainty.

The simplest scenario we address is a GW process in which the branching probabilities $\{p_s^i\}$ are independent and identically distributed random variables, all drawn from the same distribution with average $\langle p_s^i\rangle = \langle p_s\rangle$, and variance $\sigma^2[p_s^i] = \sigma^2[p_s]$. This scenario effectively reduces the complexity of the model by assuming a single representative average branching probability $\langle p_s\rangle$ which can be inferred from Main Text Figs. 4d-e and 5d,h, using Lagrange's mean value theorem. The variance $\sigma^2[p_s]$ then accounts for the biological noise present in the system.

The generalized GW process described by Eq. (41) is compounded with $i$ independent and identical probability distributions, extending Eqs. (42)-(43) to

$$E[X_i] = E_{\{p_s^i\}}\left[E\left[X_i\middle|\{p_s^i\}\right]\right] = (2\langle p_s\rangle)^i = \langle X_i\rangle_{no\ noise}, \tag{44}$$

$$\sigma^2[X_i] = E_{\{p_s^i\}}\left[\sigma^2\left[X_i\middle|\{p_s^i\}\right]\right] + \sigma^2{}_{\{p_s^i\}}\left[E\left[X_i\middle|\{p_s^i\}\right]\right] = 4(\langle p_s\rangle(1-\langle p_s\rangle) - \sigma^2[p_s])\frac{\left(4(\langle p_s\rangle^2+\sigma^2[p_s])\right)^i-(2\langle p_s\rangle)^i}{4(\langle p_s\rangle^2+\sigma^2[p_s])-2\langle p_s\rangle} + \left(4(\langle p_s\rangle^2+\sigma^2[p_s])\right)^i - (2\langle p_s\rangle)^{2i} \tag{45}$$

which describe a stochastic process with an average identical to a standard Galton-Watson with fixed probability $\langle p_s\rangle$ (Eq. (37)), but with inflated variance compared to Eq. (38). The second term of the variance, the "environmental" noise $\sigma^2{}_{\{p_s^i\}}\left[E\left[X_i\middle|\{p_s^i\}\right]\right]$, is multiplicative (variance of a product) and dominates as depth increases, whereas the branching noise $E_{\{p_s^i\}}\left[\sigma^2\left[X_i\middle|\{p_s^i\}\right]\right]$ contributes only a lower-order correction.

A similar set of equations applies when $\{p_s^i\}$ are independent but not identically distributed random variables, with each $p_s^i$ drawn from a distribution characterized by a specific mean $\langle p_s^i\rangle$ and variance $\sigma^2\left[p_s^i\right]$ that depend on the distance $i$ from the soma. This scenario effectively captures the gradual decline in branching probability observed in Fig. 4e, where $\langle p_s^i\rangle$ exhibits a decreasing trend as $i$ increases, converging to $\langle p_s\rangle = 0.5$. Despite this decay in expected values, the developmental noise $\sigma^2\left[p_s^i\right]$ remains approximately constant, suggesting a consistent level of variability in the branching process across different layers.

As we will demonstrate next, although Eqs. (44)–(45) partially resemble the expectation and variance of a GW process without noise, the introduction of multiplicative stochasticity fundamentally alters the system's behavior. This added variability transforms the distribution of $X_i$, enabling the emergence of a lognormal distribution, which would not occur in a purely deterministic branching process (see Section 4.5 Galton-Watson Simulations, "The role of noise in the Galton-Watson process").

#### Section 4.4: How Lognormality arises in the Galton-Watson Process

In the supercritical regime ($p_s > 0.5$), where a neuron has a finite probability of growing indefinitely, the expected number of segments at a distance $n$ from the soma follows a multiplicative growth pattern. Each new generation of segments is approximately equal to the previous generation multiplied by a constant greater than one, which depends on the branching probability. This self-reinforcing growth results in an exponential increase in the number of segments, so that the latest generation is the most populated. This contrasts sharply with the subcritical regime, where the number of segments decreases exponentially with the distance from the soma.

Motivated by the supercritical and asymptotic criticality patterns observed in Figs. 4e and 5d,h, we approximate the total length of a neuron as:

$$L_n = \langle l \rangle T_n = \langle l \rangle \sum_{j=1}^{n} X_j \approx \langle l \rangle X_n \, , \tag{46}$$

where $\langle l \rangle$ is the average length of a segment and $T_n$ represents the total size of the neuronal tree. Given the exponential growth characteristic of the supercritical regime, the latest generation, $X_n$, dominates the total tree size $T$.

To explore the properties of neuronal length, we approximate $X_n$ using its expected value from Eq. (42), allowing us to express:

$$X_n \approx X_0 \left( 2^n \prod_{k=1}^{n} p_s^k \right). \tag{47}$$

This equation reveals the multiplicative nature of neuronal growth, where the number of segments at distance $n$ is determined by a product of branching probabilities across generations.

Here, instead of analyzing the complete compounded distribution of the final generation size, $X_n$, we focus on its expected value, which is itself a random variable with multiplicative properties, as derived in Eq. (42). This approximation holds well, as we demonstrate later in Section 4.5, Figure S29, because the stochasticity typical of a standard GW process is negligible compared to the developmental noise introduced in Section 4.3, making the multiplicative central limit dominate over the expected standard central limit for GW processes.

In the presence of biological noise, as described in Section 4.3, the branching probabilities $\{p_s^k\}$ vary randomly. As a first-order approximation, this means that $X_n$ behaves as a random variable governed by a multiplicative process, where a sequence of independently drawn $\{p_s^k\}$ values is multiplied together. Consequently, the distribution $P(X_n)$ across multiple neurons follows the Multiplicative Central Limit Theorem, leading to a log-normal approximation:

$$P(X_n) \approx \frac{1}{X_n\sqrt{s^2 2\pi}} e^{-\frac{(\log X_n - \mu_m)^2}{2\sigma_m^2}} \tag{48}$$

where $\mu_m$ and $\sigma_m^2$ depend on the specific variability affecting $\{p_s^k\}$. If the probabilities $\{p_s^k\}$ are independent and identically distributed random variables, then:

$$\mu_m = \log(X_0) + n\left(\mu_q + \log(2)\right), \tag{49}$$

$$\sigma_m^2 = n\sigma_q^2\ , \tag{50}$$

where $\mu_q = \langle \log(p_s) \rangle$ and $\sigma_q^2 = \sigma^2(\log(p_s))$. If the probabilities $\{p_s^k\}$ are not drawn from the same probability distribution, Eqs. (49)-(50) generalize to

$$\mu_m = \log(X_0) + \sum_{k=1}^{n} \left(\mu_q\right)_k + n\log 2\,, \tag{51}$$

$$\sigma_m^2 = \sum_{k=1}^{n} \left(\sigma_q^2\right)_k, \tag{52}$$

where $\left(\mu_q\right)_k = \langle \log(p_s^k) \rangle$ and $\left(\sigma_q^2\right)_k = \sigma^2(\log(p_s^k))$.

Note that Eq. (48), invokes the multiplicative central limit theorem. The natural question is how many multiplicative steps are required for the lognormal approximation to be valid. While the precise number depends on the properties of the random variable and the degree of precision required, a common heuristic is that the limit behavior becomes reliable after approximately 10 multiplications (or even faster– according to Benford's Law by Alex Ely Kossovsky[65], the lognormal convergence takes place after five iterations, see page 222 in the book). Since the typical neuronal tree depth in our data is around 30 layers, we are way beyond the theoretical limits, hence the use of this approximation is appropriate. To confirm this, in Section 4.5 we carried out a detailed numerical study to test how the lognormal distribution emerges as the best fit depending on the number of generations in Figures S28-29.

To further improve the approximation of the total tree size given in Eq. (46), we can incorporate multiple generations leading up to the final one. This refinement is justified by the fact that the sum of lognormal distributions is itself well approximated by another lognormal distribution[66,67], ensuring a more accurate representation of the overall neuronal growth dynamics.

Given the observed empirical parameters, the final generation will be only ~1.1 times larger than the previous one under the observed empirical parameters. However, this approximation only affects the intercept in the final expression, not the slope, which is what we aim to derive here.

We could indeed approximate total tree size not as the final layer, but instead as a constant multiple of the final generation size (e.g., the sum of the last few layers). This approach introduces a constant pre-factor in the expression, shifting the intercept in Eq. 9 in the paper, but leaves the scaling behavior (slope) unchanged. In fact, while Eq. 9 predicts a slope of nearly 20 for the fit shown in Figure 5h, we empirically observe an intercept slightly larger than 2—consistent with a small pre-factor difference. Eq. 10 in the paper, which depends on the scaling rather than the absolute magnitude, would remain unaffected by this approximation. Regardless of whether we consider this pre-factor or not, we use the original approximation to easily interpolate between tree length and tree size, which simplifies many calculations.

Overall, our findings reveal two key insights into neuronal growth. Firstly, we demonstrate that the underlying multiplicative process driving neuronal expansion gives rise to a lognormal distribution, as analytically predicted in Eq. (48). This distribution captures the inherent variability in neuron size resulting from stochastic branching events. Secondly, Eqs. (49)-(51) establish a formal link between the distribution of neuronal lengths, $P(L_n)$ and the empirically measurable parameters that define individual neurons, as illustrated in Fig. 5h. This connection not only validates the theoretical framework but also enhances our understanding of how microscopic branching behaviors translate into macroscopic structural diversity.

In summary, we find that

- Neurons can be modeled as random binary trees governed by a Galton-Watson process.
- The empirical data indicates (Figs. 4d and 5d) that initially, neuronal growth follows a strongly supercritical regime ( >> 0.5), leading to exponential expansion for several generations before asymptotically stabilizing toward a critical phase (~0.5).
- Developmental noise introduces variability in the branching probabilities across generations, transforming the deterministic exponential growth with Gaussian fluctuations into a multiplicative stochastic process. This variability gives rise to a lognormal distribution, which characterizes the number of segments at a given distance from the soma and ultimately shapes the overall neuronal tree structure.

### Section 4.5: Galton-Watson Simulations

In this section, we present results from Galton-Watson simulations to validate two key aspects of our theoretical framework:

1. The impact of developmental noise, as discussed in Section 4.3, on neuronal growth dynamics.
2. The accuracy of Eqs. (48)–(52) in describing the evolution of the Galton-Watson process, ensuring that our analytical predictions align with simulated outcomes.

<u>The role of noise in the Galton-Watson process</u>

We begin by simulating Galton-Watson (GW) processes with a constant survival probability of $p_s = 0.65$, examining how the stochastic process evolves both with and without variability in the survival probability across generations. To introduce the empirically observed developmental noise, we model $p_s$ as the mean of a Beta distribution with a standard deviation $\sigma[p_s]$ =0.1, a value informed by the FlyWire data. The initial population size is set to $X_0$ = 20, and the maximum distance from the soma is 50. For each scenario, we generate 5,000 simulated neuronal trees.

The results of these simulations, summarized in Fig. S28, compare the deterministic case (left) with the noisy case (right), highlighting several key findings:

1. **Estimating $p_s$ from Data:**
   Across 5,000 realizations, different maximum likelihood estimation methods can be employed to infer $p_s$ directly from the data. Fig. S28a (no noise) shows minimal deviations, which progressively diminish as the number of generations increases. In contrast, Fig. S28b (with noise) demonstrates persistent variability in $p_s$ across generations, in line with the empirical data (Fig. 4e in the manuscript).
2. **Convergence of Standard Deviation $\sigma(p_s)$:**
   Without noise, the standard deviation of $p_s$ across realizations steadily approaches zero as the number of segments per generation increases (Fig. S28c). When biological noise is introduced, the standard deviation instead stabilizes at 0.1 (Fig. S28d). At no point do these fluctuations become comparable, confirming that the noise-induced variability is intrinsic rather than a sampling artifact.
3. **Emergence of Lognormality with Noise:**
   Using a Kolmogorov-Smirnov (KS) test to assess the distribution of segment numbers across generations, we find that lognormality only emerges in the presence of noise. Specifically, in Fig. S28f, a lognormal distribution becomes the best approximation from generation 20 onward, whereas Fig. S28e (without noise) exhibits a standard additive central limit behavior.
4. **Differences in Distribution Shape Despite Identical Mean:**
   Although both cases converge to the same expected value (Eq. 44), their distribution shapes differ significantly. In Fig. S28g, when plotted in logarithmic space, both the final

generation and the total tree size follow the expected additive central limit behavior in the absence of noise, with considerable skewness in the logarithmic space. Conversely, in Fig. S28h, with noise, both distributions clearly follow a lognormal form, as predicted in Section 4.4 (Eqs. 46–48), the logarithmic skewness disappears.

These results numerically confirm our analytical predictions that developmental noise fundamentally alters the statistical properties of neuronal branching, shifting the growth process from an additive to a multiplicative framework, thereby shaping the overall structure of neuronal trees.

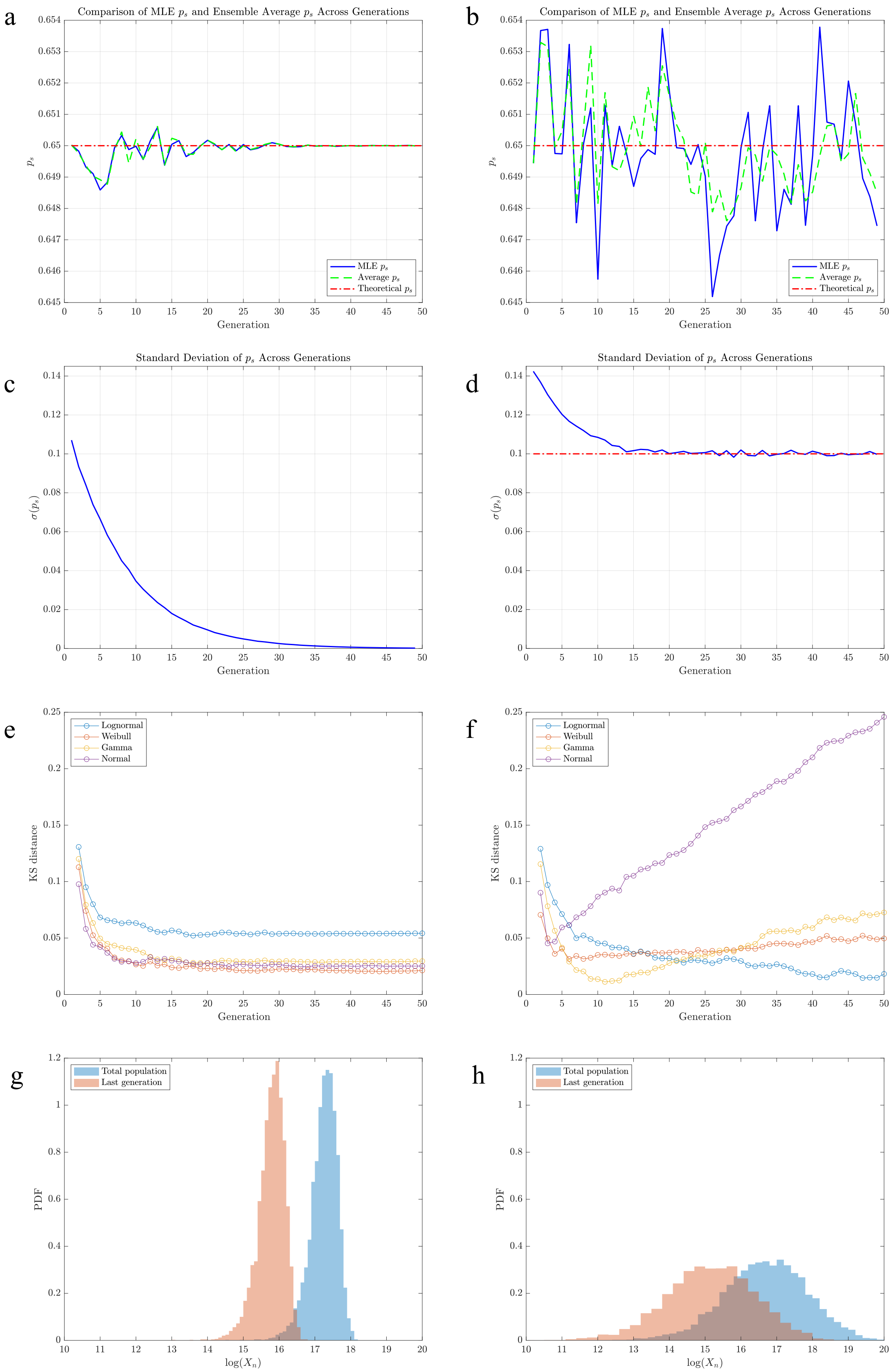

a
Comparison of MLE $p_s$ and Ensemble Average $p_s$ Across Generations
$p_s$
Generation
MLE $p_s$
Average $p_s$
Theoretical $p_s$
b
Comparison of MLE $p_s$ and Ensemble Average $p_s$ Across Generations
$p_s$
Generation
MLE $p_s$
Average $p_s$
Theoretical $p_s$
c
Standard Deviation of $p_s$ Across Generations
$\sigma(p_s)$
Generation
d
Standard Deviation of $p_s$ Across Generations
$\sigma(p_s)$
Generation
e
KS distance
Generation
Lognormal
Weibull
Gamma
Normal
f
KS distance
Generation
Lognormal
Weibull
Gamma
Normal
g
PDF
$\log(X_n)$
Total population
Last generation
h
PDF
$\log(X_n)$
Total population
Last generation

**Supplemental Figure 28: The Role of Noise in the Galton-Watson Process**

Comparison of deterministic (left) vs. stochastic (right) Galton-Watson processes, highlighting the impact of developmental noise on the evolution of neuronal trees. The deterministic case (left) follows a standard Galton-Watson process with a fixed $p_s = 0.65$, while the stochastic case (right) introduces variability $\sigma[p_s]$ =0.1. Each scenario is simulated over 5,000 realizations with an initial population of $X_0$ = 20 and a maximum of 50 generations. *Estimation of $p_s$ Over Generations* **a)** No Noise: Estimated $p_s$ across generations remains stable with minimal deviations, which disappear as the number of segments per generation increases. **b)** With Noise: The estimated $p_s$ fluctuates persistently across generations, reflecting the intrinsic biological variability observed in real neurons. *Standard Deviation $\sigma[p_s]$ Across Generations.* **c)** No Noise: The standard deviation $\sigma[p_s]$ decreases with generation number, eventually approaching zero, confirming the deterministic nature of the process. **d)** With Noise: The standard deviation stabilizes at $\sigma[p_s] = 0.1$, demonstrating persistent variability in $p_s$ that is intrinsic rather than a sampling artifact. *Kolmogorov-Smirnov (KS) Distance for Lognormality*. **e)** No Noise: The KS test suggests that segment number distributions follow a Gaussian-like trend due to additive accumulation of variance. **f)** With Noise: Lognormality emerges from generation 20 onward, confirming that multiplicative variability in $p_s$ transforms the process from Gaussian to lognormal. *Distribution of Segment Number in Log Space and Total Tree Size.* **g)** No Noise: The final generation and total tree size exhibit significant skewness in log-space, consistent with an additive process. (H) With Noise: The distribution follows a clear lognormal shape, as predicted by Eqs. (44–46), confirming that developmental noise transforms the process into a multiplicative framework.

Exponential Decay of $p_s$ Across Layers

Fig. 4d and 4e indicate that the branching probability $p_s$ is not constant, but starts at high values near the soma (low layer number) and gradually decreases, converging toward 0.5 for deeper layers. Motivated by these observations, we approximate $p_s$ with an exponential decay function, capturing the biological subtleties in the expected value of $p_s$, starting from 0.95 and decreasing to 0.5 over 60 generations, with a standard deviation $\sigma[p_s]$ =0.1. To assess the impact of this decay, we compare simulation results across 100,000 sampled neurons, initialized with different population sizes: $X_0$=3, $X_0$ =16, and $X_0$ = 32, corresponding to the left, middle, and right panels of Fig. S29, respectively.

Key Findings from Fig. S29:

1. **Lognormality Emerges as the Best Fit**
   Across all scenarios, lognormality consistently emerges as the best fit for the distribution of neuronal segments from approximately generation 20, in agreement with the convergence of the multiplicative central limit in Eq. (48). As the initial population size increases, the KS distance slightly decreases, indicating a more robust convergence to lognormality. This occurs because larger initial populations mitigate discrete stochastic effects (Fig. S29a, b, and c).
2. **The Logarithmic Mean Peaks Near the Knee of the Exponential Decay**
   The logarithmic mean of each generation reaches its peak when $p_s$ approaches 0.5. At this stage, the multiplicative terms in Eq. (47) are increasingly likely to fall below 0.5 due to biological noise. As a result, in Eq. (51), the terms $\{\mu_q\}_k$ following this knee point are more likely to contribute small negative values, reducing the overall mean. Consequently, in logarithmic space, the generation closest to the total tree size distribution is not the final layer but rather the one where the logarithmic mean peaks—which, in this case, occurs at Generation 39, regardless of the initial population size. This pattern is clearly observed in Fig. S29d, e, and f.
3. **Lognormal Fits the Number of Segments Across Generations and the Total Tree Size $T_n$**
   The number of segments at Generation 39, the final generation, and the total tree size all exhibit convincing lognormal distributions, confirming the predictions of our model. However, for $X_0$= 3, the small initial population leads to occasional outlier trees with very few segments, for which early tree extinction does not allow the continuous multiplicative central limit to emerge (Fig. S29g, h, and i).
4. **Lognormal Distribution Minimizes the KS distance Across All Scenarios**
   Lognormal consistently provides the best KS distance fit for Generation 39, the final generation, and the total tree size, regardless of the initial population size. However, for $X_0$ = 3, the presence of outliers in Panel g inflates the KS-distance of the lognormal distribution (Fig. S29j, k, l).
5. **Eq. (47) Accurately Predicts the Logarithmic Mean Across Generations**
   In all scenarios, Eq. (51) provides an accurate estimation of the logarithmic mean for segment distributions across generations, demonstrating the predictive power of our theoretical framework (Fig. S29m, n, and o).

6. **Eq. (48) Accurately Predicts the Logarithmic Standard Deviation Across Generations**
   Similarly, Eq. (52) effectively captures the logarithmic standard deviation across generations. As expected, when the initial population size is small ($X_0$ = 3), discrete stochastic effects increase the measured standard deviation, introducing additional variability (Panels p, q, and r).

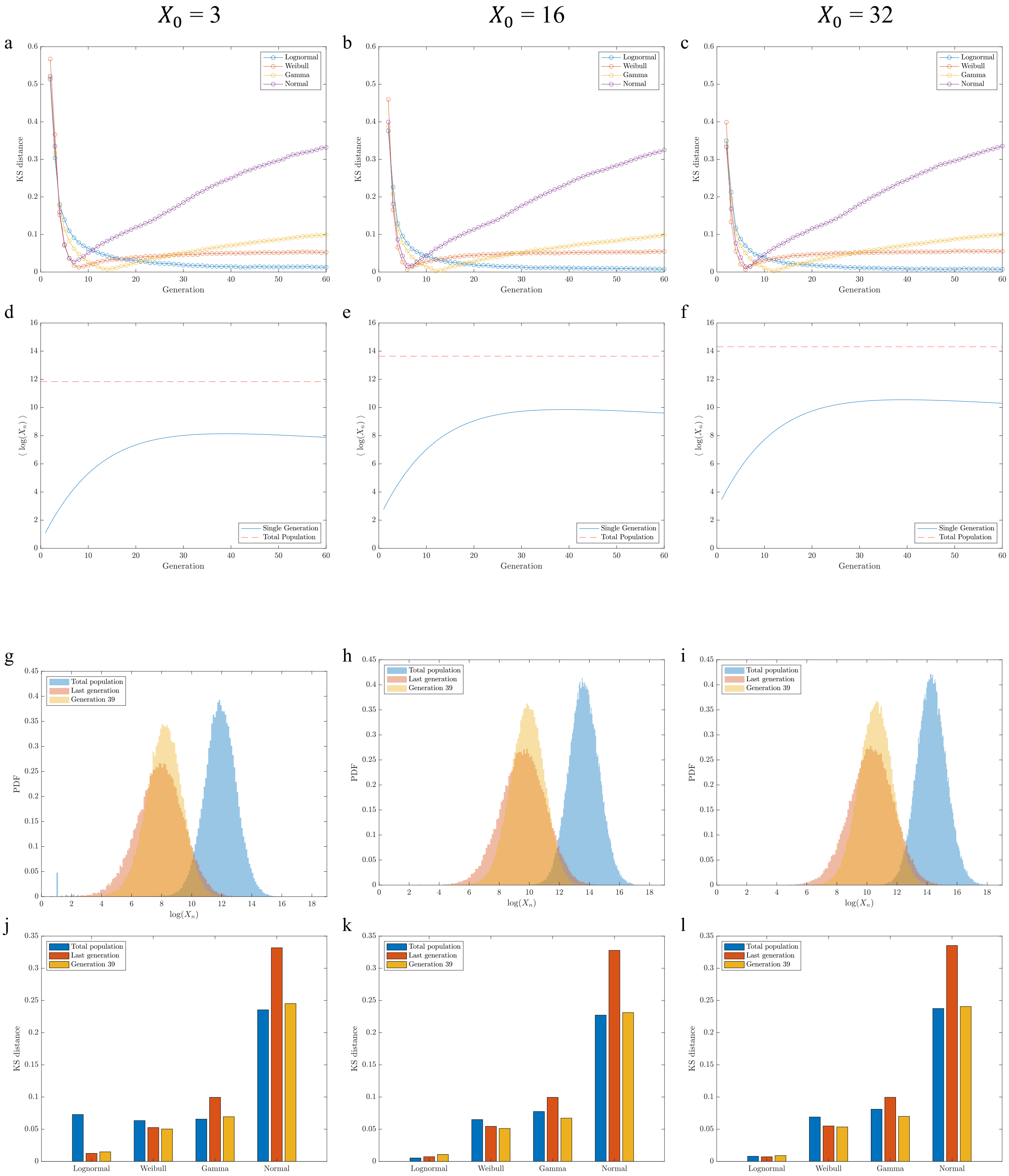

$X_0 = 3$
$X_0 = 16$
$X_0 = 32$
a
b
c
Lognormal
Weibull
Gamma
Normal
KS distance
Generation
d
e
f
Single Generation
Total Population
⟨ log($X_n$) ⟩
Generation
g
h
i
Total population
Last generation
Generation 39
PDF
log($X_n$)
j
k
l
Total population
Last generation
Generation 39
KS distance
Lognormal
Weibull
Gamma
Normal

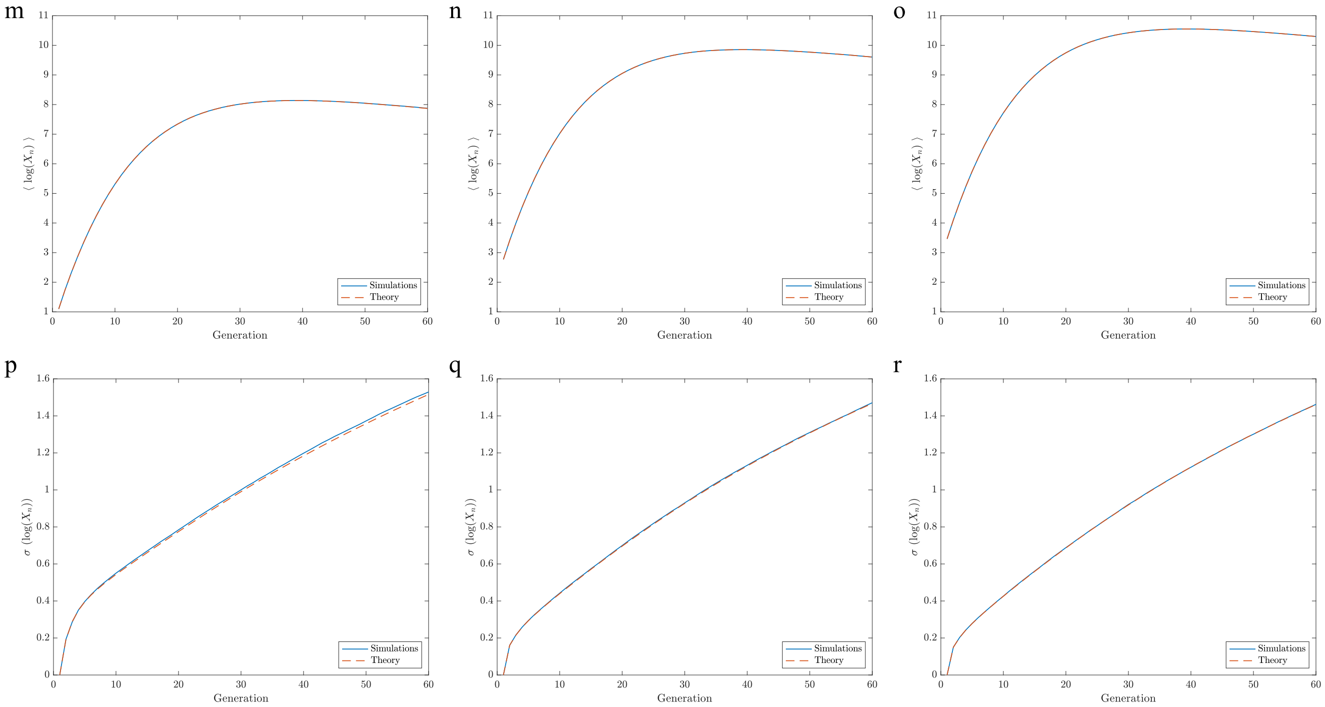

**Supplemental Figure 29: Tree simulations with exponential decay of $p_s$.** Galton-Watson simulations incorporating an exponentially decaying branching probability $p_s$, which starts at 0.95 and converges to 0.5 over 60 generations with a standard deviation of $\sigma[p_s]$ =0.1. The simulations analyze 100,000 neuronal trees, comparing cases with different initial populations: $X_0$3 (left), $X_0$ =16 (middle), and $X_0$=32 (right). **a)-c)** Across all cases, lognormality emerges as the best fit for segment distributions from generation 20 onward, aligning with Eq. (42). Larger initial populations ($X_0$) reduce stochastic fluctuations, leading to a better convergence to lognormality, as indicated by decreasing KS distances. **d)-f)** The logarithmic mean of segment distributions peaks when $p_s$ reaches 0.5. After this point, multiplicative noise causes terms in Eq. (45) to contribute small negative values, reducing the overall mean. This results in Generation 39 (rather than the final generation) being the closest to the logarithmic mean of total tree size distribution, independently of $X_0$. **g)-i)** The number of segments at Generation 39, the final generation, and the total tree size all follow a lognormal distribution, validating model predictions. For $X_0$=3, small population sizes introduce outliers where early tree extinction disrupts the expected lognormal behavior. **j)-l)** The lognormal distribution consistently provides the best KS distance fit across different generations and total tree size, with deviations for the total tree size for small $X_0$ (Panel j). **m)-o)** The theoretical prediction for the logarithmic mean (Eq. 49) closely matches simulation results, confirming its accuracy across all cases. **p)-r)** Eq. (50) correctly captures the logarithmic standard deviation across generations. For small $X_0$ (Panel p), discrete stochastic effects introduce additional variability, increasing the observed standard deviation.

#### Section 4.6 Weighted Stochastic Connectome Model

In this Section, we propose a mechanistic network model that replicates the four central empirical observations reported in our manuscript: (i) a lognormal strength distribution, (ii) a degree distribution approximating a lognormal, and (iii) a sublinear relationship between K and S.

In the previous sections, we showed empirically that the total length $L_i$ of a neuron $i$ is well approximated by a lognormal (Section 3), and that the lognormal distribution can be analytically derived from the properties of neuronal branching (Sections 4.1-4.5). Here we start from this result, in order to introduce a stochastic process that can account for the empirical observations in terms of the strengths and degrees. The key element of our approach is to interpret the neuronal length $L_i$ as the node fitness—representing a proxy of its ability to connect to other neurons[68,69]. In other words, we assume that each neuron has a synaptic budget proportional to its size. Consequently, the probability of establishing a connection with another neuron is a function of their combined finesses.

To formalize the model, let us start from the observation that the length of each neuron *i* follows a lognormal distribution, parametrized as

$$ln\, L_i \ \sim \mathcal{N}(\mu_l,\, \sigma_L^2).$$

For each pair of neurons, we write the latent weight (or potential synapse count between two neurons) as

$$W_{ij}^{(0)} = L_i L_j$$

In other words, we have $\ln\, W_{ij}^{(0)} = \ln\, L_i + \ln\, L_j \sim \mathcal{N}(2\mu_L, 2\sigma_L^2)$. This latent weight represents the capability of establishing synapses between neuron $i$ and $j$, based on their overall size. The observed network (connectome) can thus be generated from the full matrix of latent weights via probabilistic weight-based thinning. Accordingly, we define the probability of observing a link between neuron $i$ and $j$ as

$$p_{\text{keep}}(L_i L_j) \propto (L_i L_j)^{\alpha}.$$

When the link between neurons $i$ and $j$ is retained, its synaptic weight will be equal to its latent weight, i.e., $W_{ij} \ = W_{ij}^{(0)}$.

Given the measured set of neuronal lengths $\{L_i\}$, the probabilistic weight-based thinning treats each neuronal pair *(i,j)* as an independent Bernoulli variable $X_{ij}$, described by the overall probability distribution

$$P\big(X_{ij} = x \mid L_i, L_j\big) = p_{\text{keep}}(L_i, L_j)\, \delta_{x,1} + \Big(1 - p_{\text{keep}}(L_i, L_j)\Big)\, \delta_{x,0},$$

with $x \in \{0,1\}$. On the other hand, the weight of each link follows

$$P\big( W_{ij} = w \mid L_i, L_j \big) = p_{\text{keep}}\big(L_i, L_j\big)\delta_{w,W_{ij}^{(0)}(L_i,L_j)} + \Big(1 - p_{\text{keep}}\big(L_i, L_j\big)\Big)\ \delta_{w,0},$$

with $w \geq 0$.

In this model, the free parameter $\boldsymbol{\alpha}$ controls how strongly the probability of retaining a link is biased by its latent weight. The easy and logical choice is $\alpha = 1$, which implies a simple proportionality, with no additional assumptions. Deviations from $\alpha = 1$ can be rooted in physical constraints and geometry, as not all neurons can come physically close to each other—conditional for them to forming synapses—a limitations formally explored in the context of work focusing on physical networks[70]. In other words, physicality can offer systematic corrections, that could result in deviations from the straightforward approximation $\alpha = 1$. Corrections could also come from changes in the branching growth rate in response to synapse formation[64,71]. When α is small, link selection is relatively "egalitarian": even moderately strong candidate connections can be kept, which means high-degree nodes tend to accumulate a mix of link weights, and their *average link weight* rises disproportionately compared to low-degree nodes. As α increases, the rule becomes more sharply weight-biased: selection favors the heaviest latent weights almost deterministically. While α is a convenient model dial, and we can set $\alpha = 1$ without changing the overall implications of the model, to keep the model general, we will proceed with the general α case.

Next, we derive the statistical properties of degree and strength distributions, and the expected scaling between strength and degree.

**Degree Distribution P(k)**

The degree of neuron *i* represents the sum of the Bernoulli variables $X_{ij}$ with non-identical success probabilities, i.e.,

$$k_i = \sum_{\neq i} X_{ij}.$$

The probability of observing a link conditional on the fitness $L_i$ is

$$p_i = \Pr\big( X_{ij} = 1 \mid L_i \big) = \ \mathbb{E}\left[ p_{\text{keep}}\Big(W_{ij}^{(0)}\Big) \,\Big|\, L_i \right] \propto\ L_i^{\alpha}\ \mathbb{E}[L^{\alpha}] = L_i^{\alpha} \exp\big(\alpha\mu_L + \tfrac{1}{2}\alpha^2\sigma_L^2\big).$$

Therefore, conditional on $L_i$, $k_i$ behaves as a binomial distribution

$$\mathrm{P}(k_i \mid L_i) = \mathrm{Binomial}(n-1, p_i).$$

For large $n$ and small $p_i$(the "law of rare events"),

$$\mathrm{P}(k_i \mid L_i) \approx \text{ Poisson}(\lambda_i),$$

with

$$\lambda_i = \mathbb{E}[\, k_i \mid L_i \,] = \sum_{j\neq i} \mathbb{E}[p_{\mathrm{keep}}(W_{ij}^{(0)}) \mid L_i] \;\; \propto (n-1)p_i = (n-1)\mathbb{E}[L_j^{\alpha}]L_i^{\alpha} = C_\alpha L_i^{\alpha}.$$

Yet, $\lambda_i$ is not fixed but lognormally distributed ($\alpha$-power of a lognormal random variable), with

$$\ln\, \lambda_i \sim \mathcal{N}(\ln C_\alpha + \alpha\mu_L, \alpha^2\sigma_L^2),$$

leading to the degree distribution that formally follows a Poisson-Lognormal distribution (PLN),

$$\Pr(K = k) = \mathbb{E}\left[\frac{e^{-\Lambda}\Lambda^k}{k!}\right] = \int_0^\infty \frac{e^{-\lambda}\lambda^k}{k!} LN(\lambda\,, \ln C_\alpha + \alpha\mu_L, \alpha^2\sigma_L^2) d\lambda,$$

with $k \geq 0$. Multiplicative growth processes naturally yield lognormal distributions for continuous variables (e.g., axon length, synapse strength, link weight); however, when the observed outcome is discrete, such as node degrees, the appropriate analogue is the Poisson–lognormal, in which the latent lognormal rate generates discrete realizations (and what we observe in this case).

#### Node Strength Distribution P(S)

We can formally write the total node strength as

$$S_i = \sum_{j\neq i} W_{ij} = \sum_{j\neq i} X_{ij}\, W_{ij}^{(0)} \sum_{j\neq i} Y_{ij}\,.$$

Conditional on $L_i$, the term $Y_{ij}$, a Bernoulli weighted random variable, is expected to follow

$$\mathbb{E}\big[\, Y_{ij} \mid L_i \,\big] = \mathbb{E}\left[\, W_{ij}^{(0)}\, p_{\mathrm{keep}}\!\left(W_{ij}^{(0)}\right) \,\middle|\, L_i \,\right] \propto \; L_i^{1+\alpha}\; \mathbb{E}[L^{1+\alpha}]$$
$$= L_i^{1+\alpha} \exp\,((\alpha+1)\mu_L + \frac{1}{2}(\alpha+1)^2\sigma_L^2).$$

Therefore, the conditional expected strength is

$$\mathbb{E}[S_i \mid L_i] = \sum_{j\neq i} \mathbb{E}[p_{\mathrm{keep}}(W_{ij}^{(0)})W_{ij}^{(0)} \mid L_i] \;\; \propto \;\; (n-1)\mathbb{E}[L_j^{\alpha+1}]L_i^{\alpha+1} = c_\alpha L_i^{\alpha+1}.$$

The total distribution $\mathrm{P}(S_i \mid L_i)$ is a compound Poisson distribution with lognormal jump sizes (CP-LN), which can be approximated as a lognormal distribution (($\alpha + 1$)-power of a lognormal random variable), with

$$\ln\, S_i \sim \mathcal{N}(\ln c_\alpha + (\alpha+1)\mu_L, (\alpha+1)^2\sigma_L^2).$$

#### Scaling law S vs k (Eq. (5) in the manuscript)

Leveraging the conditional expectations $\mathbb{E}[k_i \mid L_i]$ and $\mathbb{E}[S_i \mid L_i]$, we eliminate the dependence from $L_i$, i.e.,

$$\mathbb{E}[k_i \mid L_i] \propto L_i^{\alpha} \;\Rightarrow\; L_i \propto k_i^{1/\alpha}, \alpha > 0$$

and substitute into $\mathbb{E}[\, S_i \mid L_i \,]$, finding the formal relationship between degrees and synapses,

$$\mathbb{E}[S_i \mid K_i] \propto k_i^{1+1/\alpha}.$$

If we invert this relation, we formally arrive at Eq. (5) in the paper, i.e.,

$$k_i \propto S_i^{\beta}$$

with

$$\beta = \frac{\alpha}{\alpha+1}.$$

In the simplest scenario of $\alpha = 1$, we obtain $\beta = 1/2$. However, for any $\alpha > 0$ we expect the degree to scale sub-linearly with the node strength. This analytical prediction is in excellent agreement with the empirical results of Figure 3g, where we document a very clear sublinear scaling in the range of $0.6 < \beta < 0.9$ for the different connectomes.

**Simulations**

Inspired by the parameters observed in the Hemibrain/MANC connectomes, we generated a brain network with the parameters

- $N = 20{,}000$ neurons
- $\mu_{\ln k} \approx 4.5$,
- $\sigma_{\ln k} \approx 1.0$,
- Mean degree $k \sim 130$.
- $\alpha = 1$

parameters chosen to test the potential agreement between the theoretical predictions and simulations for a system with features comparable to real brains. It is important to emphasize that the only lognormal distribution that the model uses as input is the fitness parameter $\ln L \sim \mathcal{N}(0,1)$, which in turn relates to the branching process. Degree and strength distributions are determined through the weight thinning generative process described in the earlier paragraphs.

Overall, Figure S30 documents an excellent agreement between the theoretical predictions and the numerical simulations. We start by analyzing the resulting degree distribution (Figure S30 a and b). As predicted in Section "Degree Distribution P(k)", the degree distribution is well fitted by a

Poisson-Lognormal (PLN), which, as the fits show, is well approximated with a Lognormal distribution (offering a better fit than Gamma or Weibull distributions, Figure S30b).

Figure S30 c and d confirm that the strength distribution has a lognormal nature (a derived about in the Section Node Strength Distribution P(S)), which once again fits better than Gamma and Weibull models.

Finally, Figure S30e corroborates the theoretical scaling prediction $S \propto k^2$, and compares it to a linear scaling with the same constant of proportionality.

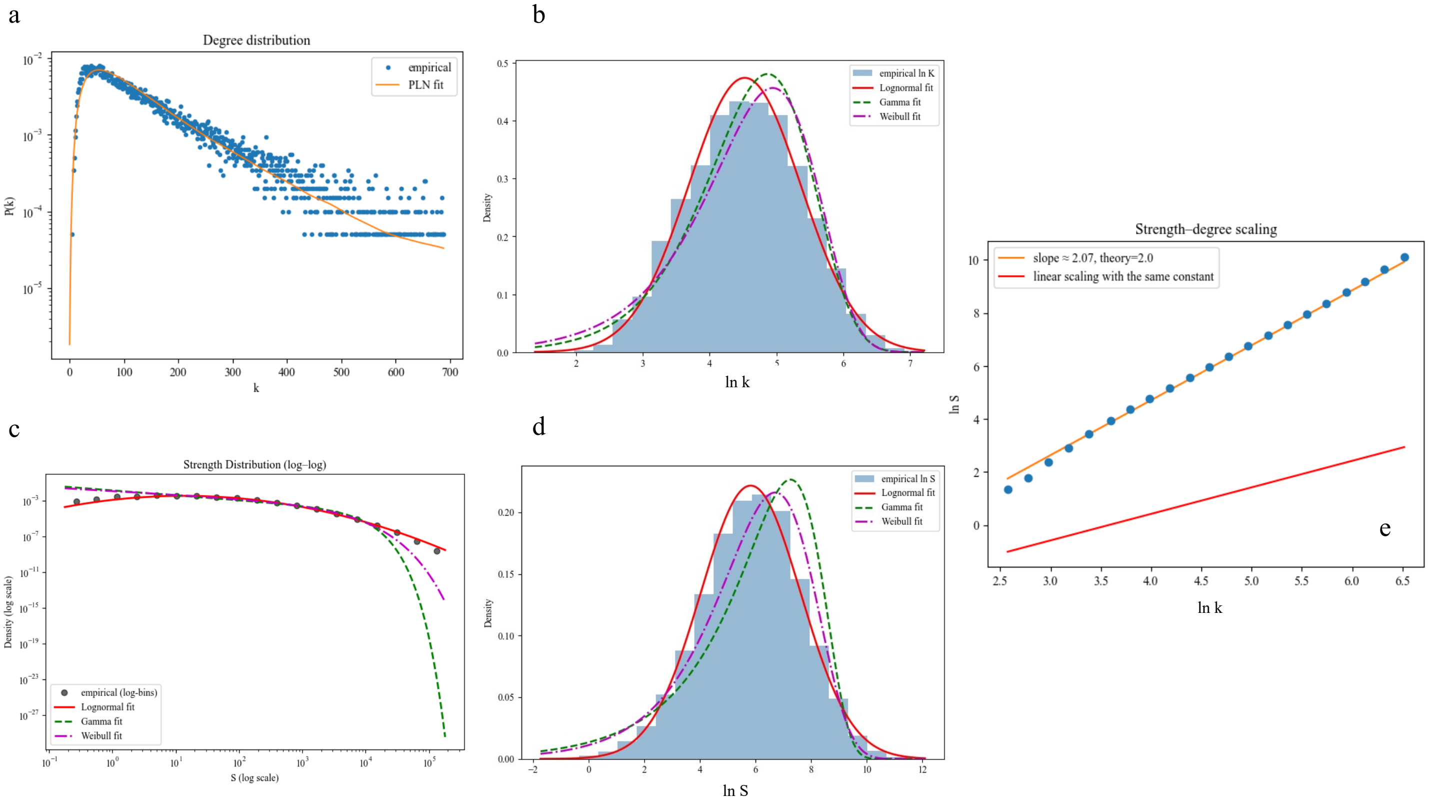

**Supplemental Figure 30: Weighted Thinning Model for Brain Networks with alpha=1, $N = 20,000$, $\mu_{ln\,k} \approx 4.5$, $\sigma_{ln\,k} \approx 1.0$, mean degree $k \sim 130$.** Panels a and b display the statistical behavior of the degree distribution compared to Poisson-Lognormal (PLN), and standard continuous candidates such as Lognormal, Gamma, and Weibull. Panels c and d display the statistical behavior of the strength distribution compared to Lognormal, Gamma, and Weibull. Panel e shows the observed superlinear scaling between S and k.

#### Section 4.7: Relation between power-law-distributed synaptic weights and synaptic strengths of neurons

Previous research has offered evidence that the synaptic weights $s_{ij}$ between neuron $i$ and $j$, representing the total number of synapses observed between the neurons, is well approximated by a power-law[72,73]. This finding has been reconfirmed in the recent data of the complete FlyWire connectome[73,74]. Specifically, Cirunay *et al.* recently reported that in several connectomes the link weight are predominantly scale-free distributed[73,75]. Interestingly, they also report the observation of a lognormal node strength distribution, which confirms our predictions, that the node strengths should follow a lognormal. Specifically, they write: "for the case of nonhuman connectomes [Figs. 4(e)–4(h)], we observe log-normal behaviors of source and sink node strengths." However, this lognormal observation is subsequently treated as an exception, as their modeling work predicts a power law source-sink distribution, ignoring the lognormal observation.

To test the main results of Lynn et al, and Cirunay *et al.*, we measured $P(s_{ij})$ in the eight datasets where the synaptic weights are identified, confirming its fat-tailed nature (Fig. S31-32). The strength of a neuron is the sum of its weights $S_i = \sum_{(i,j)} s_{ij}$ raising the question: could the sum of fat-tailed variables, describing the individual synaptic weights, lead to a lognormal strength distribution P(S), as we document in the paper? This is a particularly important question, as in some regimes, the sum of fat-tailed variables is expected to lead to another fat-tailed Levy distribution. As we show in this section, such fat-tailed strength distribution is expected to emerge only if the number of summands is constant. However, in our case, the number of summands is the degree $k_i$ of node $i$, which itself follows a lognormal distribution. Hence, the variable summand changes the nature of the strength distribution. Finally, we use simulations from the complete FlyWire data to support the range of validity of the obtained analytical results.

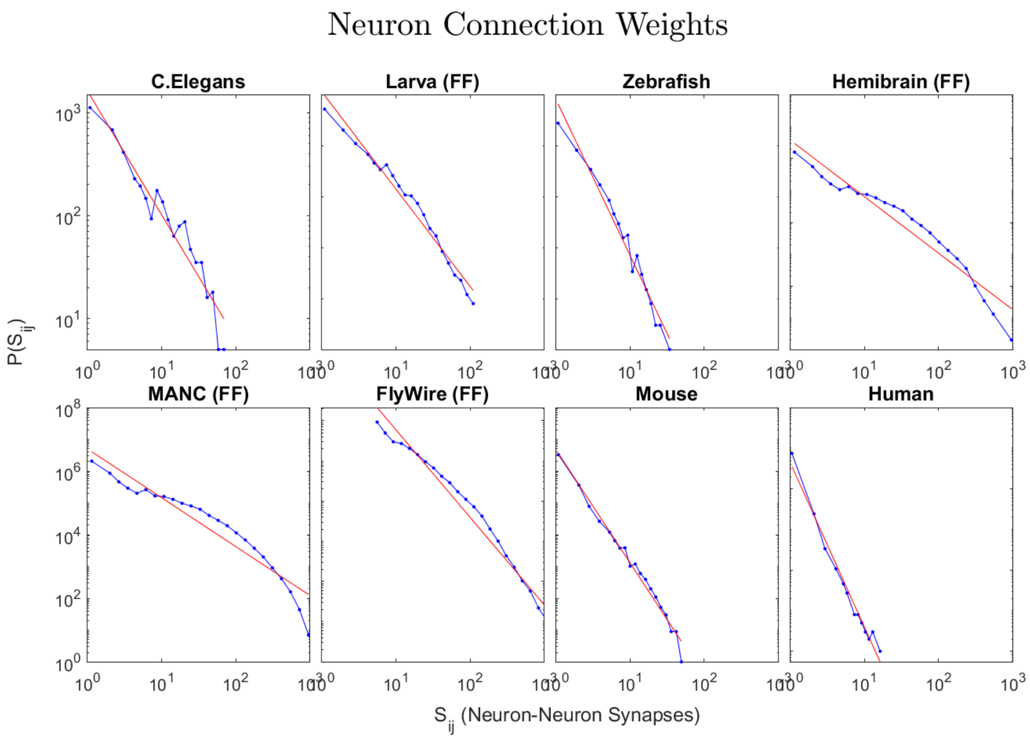

**Supplemental Figure 31: Connection weight distributions.** $s_{ij}$, the number of synapses between neuron pairs, and a single power-law fit (red line). As the data show, all $P(s_{ij})$ are fat-tailed, and some are well-approximated by a single power law, while others, like the FF dataset, show systematic deviations from it.

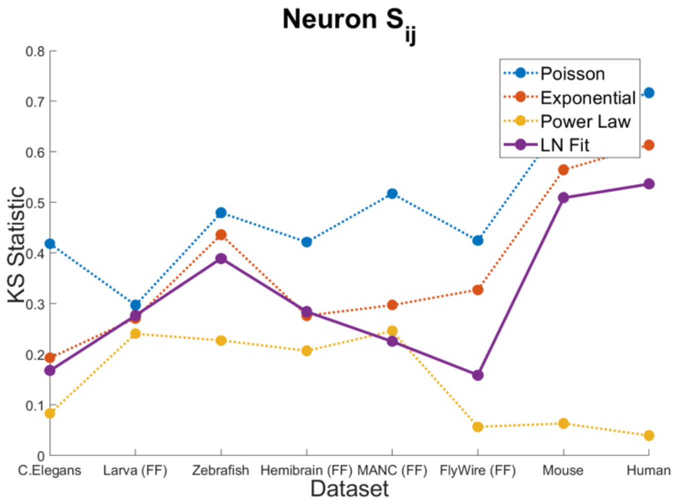

**Supplemental Figure 32: Connection weight KS statistics.** KS statistics calculated for $s_{ij}$ confirm that power-law is the most likely fit to the data.

The strength of a neuron, denoted by $S_i$, is calculated by summing the synaptic weights $\{s_{ij}\}$ associated with neuron $i$,

$$S_i = \sum_{(i,j)} s_{ij} \,. \tag{53}$$

Let us assume that the synaptic weights $\{s_{ij}\}$ are drawn from a power-law distribution defined by $P(s_{ij} = s) = (\gamma - 1) s_{min}{}^{\gamma-1} s^{-\gamma}$. The summation includes a number of terms $(i,j)$ which correspond to the degree $k_i$ of neuron *i*. When several random variables following a power-law distribution are summed, the Generalized Central Limit Theorem (GCLT) suggests that the resulting strength distribution *P(S)* converges to a stable, heavy-tailed distribution, such as a Lévy distribution. However, when the number of terms in the summation is itself a random variable, the resulting strength distribution is influenced by additional statistical noise. This compounding effect introduces variability that can dominate the overall behavior of the distribution across several regimes.

To illustrate the behavior of $P(S_i(k))$, let us rewrite Eq. (53) as

$$S_i(k) = s_{i1} + s_{i2} + \cdots + s_{ik}, \tag{54}$$

where the number of terms $k$ (degrees) follows a lognormal distribution $P(k) = LogNormal(\mu_k, \sigma_k^2)$. If $\{s_{ik}\}$ are independent and identically distributed (i.i.d.), the average and variance of $P(S_i(k))$ follow:

$$\langle S_i(k) \rangle = \langle k \rangle \langle s_{ik} \rangle \tag{55}$$

$$\sigma^2(S_i(k)) = \langle k \rangle \sigma^2(s_{ik}) + \sigma^2(k)\langle s_{ik} \rangle^2 , \tag{56}$$

describing a distribution with an expected value consistent with the sum of $\langle k \rangle$ i.i.d. $\{s_{ik}\}$ variables, but with inflated variance determined by the extra term $\sigma^2(k)\langle s_{ik} \rangle^2$ which arises from variability in *k*, the number of summands. By regrouping Eq. (56) as

$$\sigma^2(S_i(k)) = \sigma^2(k)\langle s_{ik} \rangle^2 \left(1 + \frac{\langle k \rangle \sigma^2(s_{ik})}{\sigma^2(k)\langle s_{ik} \rangle^2}\right)$$

$$= \sigma^2(k)\langle s_{ik} \rangle^2 \big(1 + f(\gamma)\big), \tag{57}$$

we can investigate under which conditions $S_i(k) \approx k\langle s_{ik} \rangle$, implying that the strength distribution is well approximated by a rescaled lognormal distribution driven by $P(k)$ rather than the power-law $P(s_{ik})$. We proceed by substituting the first and second moments of $P(k)$ and $P(s_{ik})$, particularly, $\sigma^2(s_{ik})$. In the regime $2 < \gamma < 3$, the variance of power-law distributions diverges, suggesting the overall dominance of $P(s_{ik})$. Yet, $\sigma^2(s_{ik})$ grows with the total number of weights sampled in the network, which can be approximated as the total number of links $L \approx N\langle k \rangle \approx Ne^{\left(\mu + \frac{\sigma^2}{2}\right)}$, with $N$ the network size.

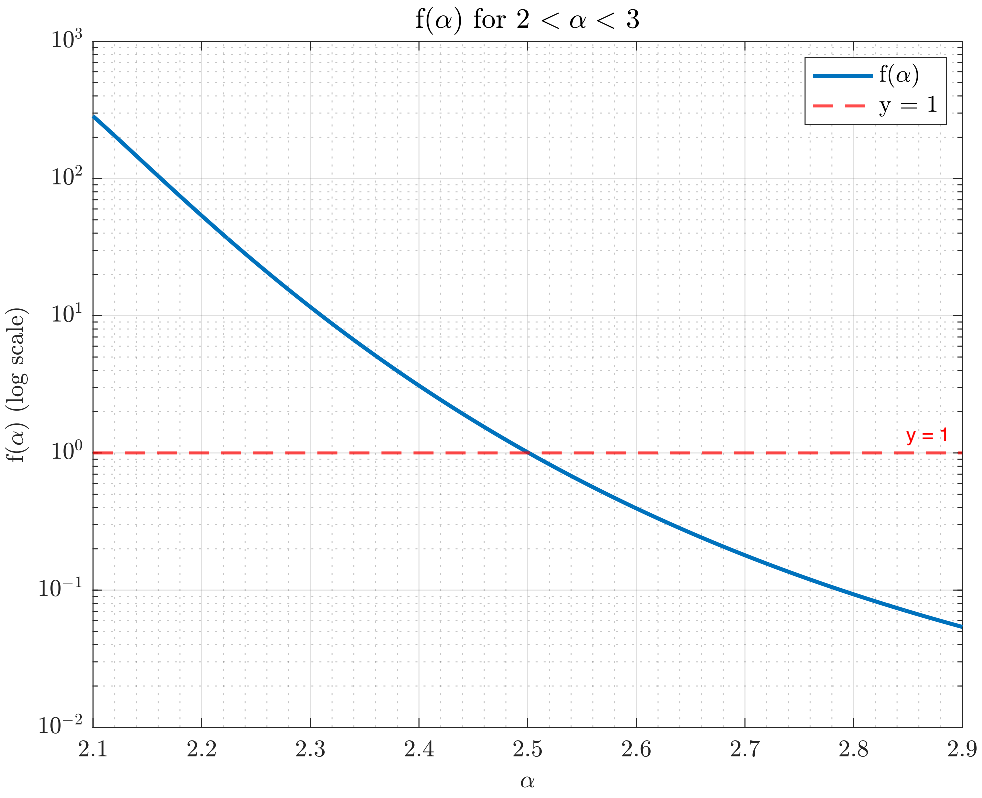

**Supplemental Figure 33: Overdispersion of $\sigma^2(s_i(k))$ compared to $\sigma^2(k)\langle s_{ik}\rangle^2$.** The correction factor in Eq. (58) progressively increases for $\gamma \to 2$. All parameters reflect the FlyWire data, except for $\gamma$, which varies between 2.1 and 2.9. The minimum estimation for $\gamma_{FlyWire}$ is 2.821.

The correction term $f(\gamma)$ in Eq. (57) is then approximated by

$$f(\gamma) \approx \frac{1}{e^{\mu+\frac{\sigma^2}{2}} \cdot \left(e^{\sigma^2}-1\right)} \cdot \left( \frac{(\gamma-2)^2}{(\gamma-1)(3-\gamma)} \cdot \left( \left( \Gamma\left(\frac{\gamma-2}{\gamma-1}\right) \right)^{3-\gamma} N^{\frac{3-\gamma}{\gamma-1}} \cdot e^{\left(\mu+\frac{\sigma^2}{2}\right) \cdot \frac{3-\gamma}{\gamma-1}} - 1 \right) - 1 \right). \quad (58)$$

Therefore, for specific values of $\mu$, $\sigma$, and $\gamma$, the lognormal distribution could outweigh the effects of the power-law tails, leading to a small $f(\gamma)$ and $S_i(k) \approx k\langle s_{ik}\rangle$.

By substituting the network features measured in the FlyWire dataset while keeping $\gamma$ as a free parameter, we can identify the range of $\gamma$ values for which $f(\gamma) \leq 1$. This range corresponds to a minimal contribution to the overdispersion of $\sigma^2(S_i(k))$. Analytically, this condition is satisfied for $\gamma^* \geq 2.501$ (as shown in Fig. S33). Indeed, the simulations show how the right tail of $P(S_i(k))$ progressively collapses on $P(k\langle s_{ik}\rangle)$ for $\gamma \to 3$, with limited contributions of the power-law tails from $\gamma = 2.5$ (Fig. S33). The behavior of $P(S_i(k))$ in the logarithmic space reveals how the body of the distribution still resembles a lognormal for $\gamma \to 2$ (Fig. S34a).

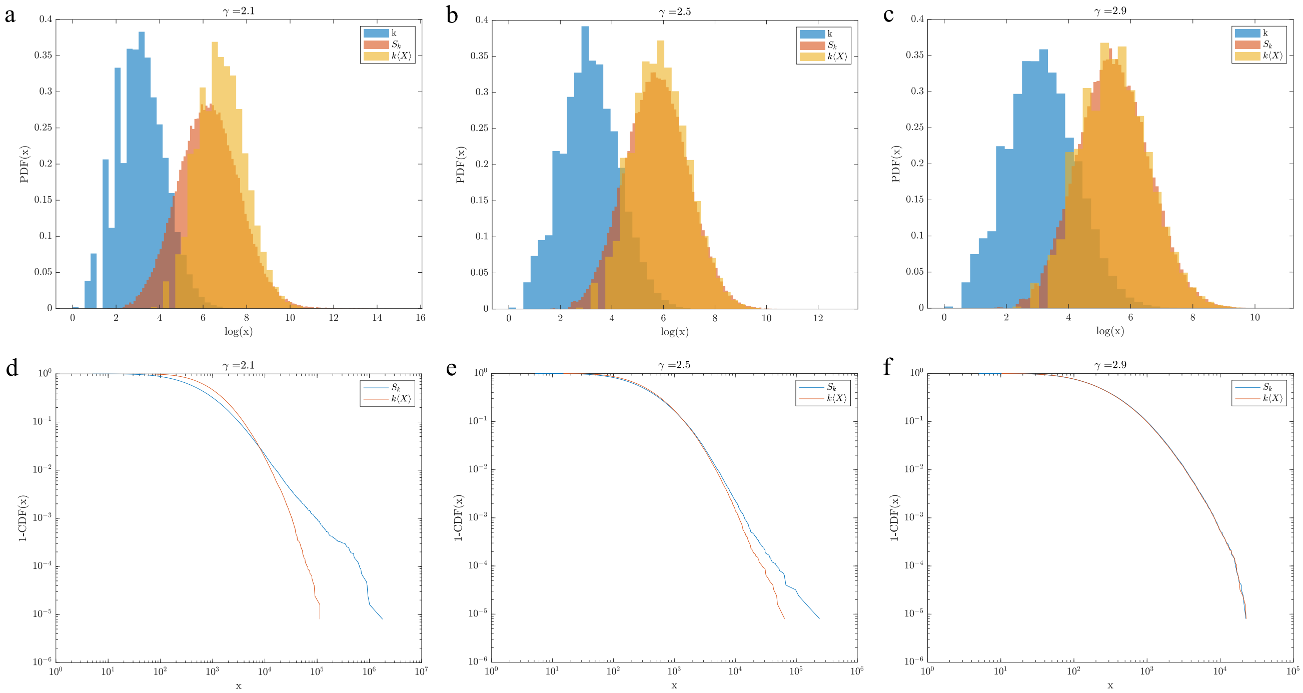

**Supplemental Figure 34: The probability distribution of $s_i(k)$ compared to $k\langle s_{ik}\rangle$.** While $P(k\langle s_{ik}\rangle)$ it behaves as a rescaled lognormal, $P(s_i(k))$ it displays a wider right tail, which progressively diverges $\gamma \rightarrow 2$. Given the FlyWire parameters, $P(s_i(k))$ is well approximated by $P(k\langle s_{ik}\rangle)$ for $\gamma \geq 2.5$. To simplify the notation, the random variable $s_{ik}$ is represented as $X$. **a)-c)** comparison of the probability density functions (PDF) in the logarithmic space. **d)-f)** comparison of the survival functions in the linear space (1-CDF).

The power-law fitting of synaptic weights in the FlyWire dataset revealed $2.821 \leq \gamma_{FlyWire} \leq 3.0571$. In this regime, $\sigma(S_i(k))$ is only minimally inflated compared to $\langle s_{ik}\rangle\sigma(k)$. This observation explains why a lognormal distribution provides an excellent approximation for $P(S_i(k))$ (Fig. S35).

While the previous estimations identified useful boundaries, synaptic weights are unlikely to be independent, prompting us to correct Eqs. (56)-(57) by considering $\{s_{ik}\}$ identically distributed and with the same pairwise covariance:

$$\sigma^2(S_i(k)) = \sigma^2(k)\langle s_{ik}\rangle^2 \left(1 + \frac{\langle k\rangle\sigma^2(s_{ik}) + \rho(\sigma^2(k) + \langle k\rangle^2 - \langle k\rangle)\sigma^2(s_{ik})}{\sigma^2(k)\langle s_{ik}\rangle^2}\right), \quad (59)$$

where $\rho$ is the correlation coefficient between different pairs of $\{s_{ik}\}$. When the total capacity of the system—represented by the feasible number of synapses—is constrained by factors such as geometry or energetic costs, $\rho$ is likely negative, further reducing the correction factor $f(\gamma)$.

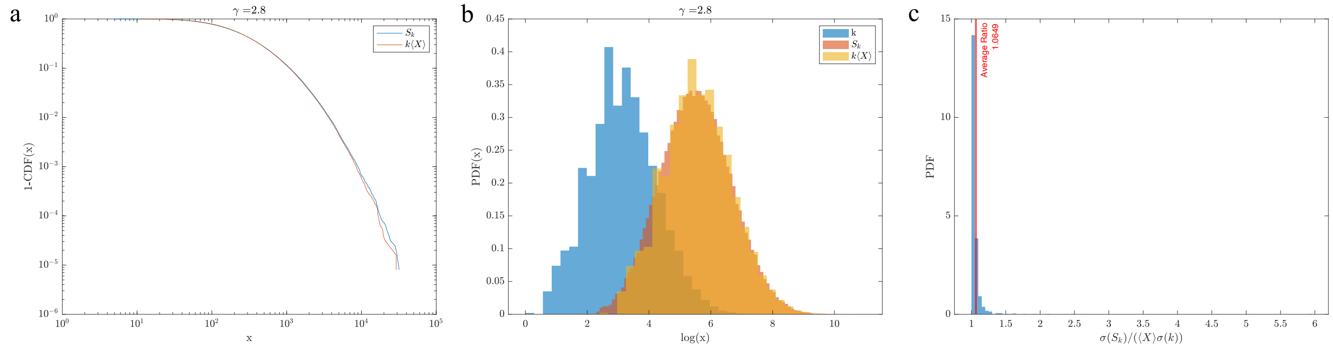

**Supplemental Figure 35: FlyWire regime.** At $\gamma = 2.8$ $s_i(k) \approx k\langle s_{ik}\rangle$ as we can appreciate by inspecting **a)** the survival function, and **b)** the probability density function in the logarithmic space. **c)** By generating 1,000 instances of datasets statistically equivalent to the FlyWire, we observe only limited inflation of the standard deviation $\sigma(s_i(k))$ compared to $\langle s_{ik}\rangle\sigma(k)$, as quantified by the average ratio over the ensemble, equal to 1.0649. To simplify the notation, the random variable $s_{ik}$ is represented as $X$.

### Section 4.8: Derivation of Rich-Club Organization from Sublinear Scaling

The rich-club phenomenon describes the tendency of high-degree nodes—often referred to as "rich nodes"—to form disproportionately more connections among themselves than would be expected by chance. Here, we show that the rich-club phenomenon is intrinsically linked to the sublinear scaling between degree and synaptic strength within the network (Eq (5) in the manuscript). This phenomenon has been observed in a variety of networks, including brain connectomes[76,77], scientific collaboration networks[78], transportation systems[79], and interbank networks[80]. While these studies reveal important structural properties of such systems, they often fall short by merely identifying the presence or absence of links among influential nodes, overlooking the essential information carried in the weights of these connections.

Here, we analyze the weighted network derived from the FlyWire dataset using the weighted rich-club measure originally introduced by Opsahl, Colizza, et al.[78] and later applied by van den Heuvel and Sporns[77] to structural brain networks generated via diffusion tensor imaging. In this framework, nodes are ranked according to their degree $k$ in increasing order. For each degree threshold $k$, we define a "club" comprising nodes whose degree exceeds this value. These $k$-clubs become progressively more selective as $k$ increases. Within each $k$-club, we count the total number of links $E_{>k}$ connecting its members and calculate the sum of the weights $W_{>k}$ associated with these links. The weighted rich-club coefficient, $\phi^w(k)$, is then computed as the ratio between $W_{>k}$ and the sum of the weights of the $E_{>k}$ strongest links across the entire network, namely,

$$\phi^w(k) = \frac{W_{>k}}{\sum_{l=1}^{E_{>k}} w_l^{rank}} \,, \tag{60}$$

where $w_l^{rank} \geq w_{l+1}^{rank}$ with $l=1, 2, \ldots E$ are the ranked weights on the links of the network, and $E$ is the total number of links. Eq. (60) provides a measure of the fraction of weights shared among rich nodes relative to the maximum possible weight they could share if all their connections represented the strongest links in the network. This quantifies the extent to which prominent nodes (i.e., high-degree neurons) dominate the flow of resources across the entire system.

To determine whether the weighted rich-club phenomenon is genuinely present or simply a product of random chance, $\phi^w(k)$ needs to be compared to a suitable benchmark. Indeed, even networks with random link and weight assignments can exhibit non-negligible $\phi^w(k)$, so a meaningful assessment requires the use of null models. For this purpose, we adopt the CSM Model (SI 2.12), a randomization procedure that preserves the network's topology while globally reshuffling the link weights. This approach offers a more stringent comparison than another commonly used method[81], which involves reshuffling both weights and the network topology while maintaining the original degree distribution *P(k)*.

The randomization disrupts the correlation between edge weights and node degrees, altering the observed relationship between strength $S$ and degree $k$ in the FlyWire dataset. Indeed, empirical observations indicate that the degree $k_i$ of a node $i$ scales with its total synaptic weight $S_i$ according to a sublinear relationship (Eq. (5) in the manuscript):

$$k_i = \alpha S_i^{\beta} \,. \tag{61}$$

Here, $\alpha$ is a proportionality constant, and $\beta<1$ signifies sublinear scaling. This implies that as the total synaptic weight of a neuron increases, its degree increases at a slower, sublinear rate. Conversely, when we can invert the relationship to express the total synaptic weight in terms of the degree:

$$S_i \sim k_i^{\frac{1}{\beta}} . \tag{62}$$

Given that $\beta<1$, the exponent $1/\beta$ exceeds unity, leading to a superlinear scaling of the synaptic strength with the degree. This superlinear relationship indicates that neurons with higher degrees tend to carry weights higher than those expected under a random weight assignment (Fig. 35a).

The superlinear scaling has profound implications for the network's topology, particularly in fostering rich-club organization. Specifically, the disproportionately high synaptic weights in high-degree nodes facilitate stronger and more numerous interconnections among these hubs. Consequently, high-degree nodes are not only numerous but also tightly interconnected, forming a robust and dense core within the network. This dense interconnectivity among hubs is the hallmark of rich-club behavior, wherein the most connected nodes collaborate to form an influential and cohesive subnetwork.

However, as we discussed earlier, a true assessment of the rich-club behavior needs to compare the result to a suitably randomized network. After randomization, the relationship between degrees and strengths becomes linear, indicating that strength and degree now convey equivalent information about the system (Fig. 37a). In this linear regime, the edge weights $\{s_{ij}\}$ are, on average, independent of the specific vertices $i$ and $j$. Consequently, the node strength $S_i$ simplifies to $S_i = \langle w \rangle k_i$, where $\langle w \rangle = \sum_{(i,j)} s_{ij}/E$ represents the average weight across the entire network (Fig. 37b). This significant disruption is reflected in the behavior of the ratio:

$$\rho^w(k) = \frac{\phi^w(k)}{\langle \phi_{null}^w(k) \rangle} \tag{63}$$

where $\langle \phi_{null}^w(k) \rangle$ represents the average value of $\phi^w(k)$ over 1,000 randomized instances. As shown in Fig. 37c, $\rho^w(k)$ is not only consistently greater than 1 but also increases across a wide range of degree values. This trend indicates the presence of a weighted rich-club effect in the FlyWire network.

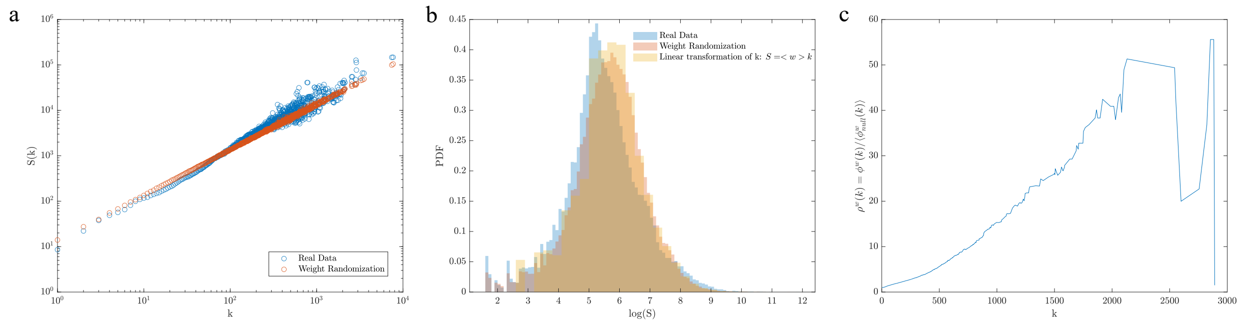

**Supplemental Figure 36: The Weighted Rich Club in the FlyWire data. a)** The relation between strength and degree is superlinear with exponent $\alpha = 1.0835$ (1.0669, 1.100), while reshuffling the weights leads to a linear relation with $\alpha = 0.99919$ (0.9966,1.0018). **b)** Weight randomization is approximately equivalent to the linear transformation $S_i = \langle w \rangle k_i$. **c)** Normalized weighted rich club $\rho^w(k)$ over 1,000 weight randomizations.

In summary, the derivation elucidates that the rich-club phenomenon is intrinsically linked to the sublinear scaling between degree and synaptic strength within the network (Eq (5) in the manuscript). The sublinear relationship ensures that as nodes become more connected, their total synaptic weight increases at a superlinear rate, thereby promoting dense interconnections among high-degree nodes. This intrinsic property of the network's scaling behavior fundamentally underpins the emergence of rich-club organization, highlighting the interplay between network topology and weighted interactions in shaping complex neural architectures.

Finally, we find that the empirically determined onset degree of the Rich Club correlates strongly with the mean parameter of the lognormal distribution, across the total, in-degree, and out-degree distribution. To find the Rich Club onset degree, we calculated the relative rich club coefficient using the methods of Lin et al[74], and found the degree at which it exceeds 1.01 (FlyWire, MANC) or 1.1 (all other datasets). We were not able to analyze the human connectome, as the core 104 neurons are weakly connected to each other, and mainly connect to fragments in the volume, thus the subgraph falls apart rapidly under the Rich Club testing process, and Rich Club regime cannot be determined.

We found that the agreement is particularly robust across all three measures (total, in-, and out-degree) for the complete datasets (*C. elegans*, Fly Larva, FlyWire), where the theory is expected to perform best. For MANC, Hemibrain, and Zebrafish, closer agreement could likely be achieved with a refined threshold heuristic, but here we adhered as closely as possible to the methodology of Lin et al.

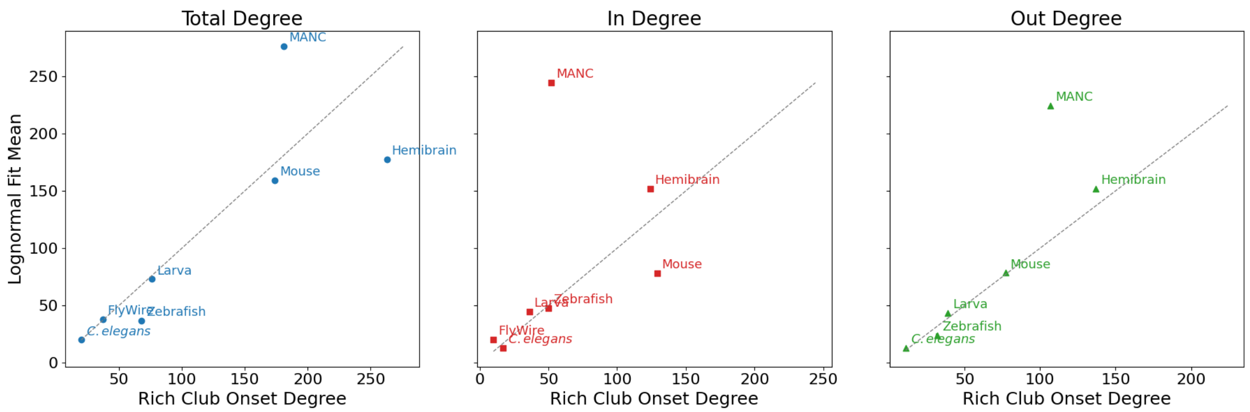

**Supplemental Figure 37: Agreement between the Rich Club Onset and Lognormal Fit Mean.** We plot the empirically calculated rich club onset degree against the mean degree of the fit lognormal. Across Total Degree, In Degree, and Out Degree, the Rich Club Onset degree has a high concordance with the lognormal fit (as highlighted by y = x dashed line).

#### Section 4.9: Related Works on Lognormality and Branching

The study of branching processes and lognormal statistics has a long history, and multiple often disjoint literatures have explored themes relevant to our work. Here we briefly summarize some key contributions and clarify how they relate to our work.

In mathematical biology, branching processes have been explored as models of population dynamics. *Branching Processes in Biology*[64] provides a thorough pedagogical treatment of this literature, and it also explores Galton–Watson processes and their applications, including discussions of random environments. However, it does not address how generation-to-generation variability alters moment dynamics, what we developed in Section 4.3 above. The emergence of lognormal distributions from a branching process is rooted in this variability, hance the text does not discuss lognormal distribution as a potential outcome.

Other classical treatments extend branching theory to population-dependent growth[71] or parametric estimation under extinction[82], respectively, but again they do not derive the generation-indexed probability generating functions (PGFs) nor do they discover the connection to lognormality. More recent works considered moment-based estimators for random graph models[83], a framework valuable for network science but not directly relevant to connectome structure or branching morphology. To put in context, our main finding, discussed in Section 4.3, 4.4 and numerically validates in Section 4.5, is that developmental noise induces stochastic variability in branching probabilities across generations, converting the classical exponential-growth framework with Gaussian fluctuations into a multiplicative stochastic process. This shift yields lognormal distributions of neuronal segments with distance from the soma, fundamentally shaping tree architecture in ways not addressed by prior literature.

Lognormality has also been repeatedly observed in neuroscience as well. For instance, lognormal statistics have been reported in neural dynamics, including the firing rates of neurons[84,85] and pairwise information transfer between neuron pairs[86]. These discoveries concern neuronal activity rather than reflecting the connectivity patterns of the underlying connectome. More recent modeling efforts propose that spatial growth can give rise to lognormal *connectivity weights*, though these models mainly predict scale-free degree distributions, omit morphological branching, and were not yet tested against connectome data[87].

Lognormal distributions have appeared in sporadically in network science as well. One interesting study assumed lognormal inputs, in the form of a lognormal fitness parameter, which then yielded a lognormal outcome. The paper has made no attempt to explain the mechanistic origin of the lognormal input[88]. Another work has provided a broad statistical survey showing that lognormal can fit the degree distribution of many social and technological networks, however, it did not attempt to offer an underlying mechanisms for the origin of this distribtion[23]. At this moment we are lacking network science models that could predict a lognormal distribution for relevant network measures—like degrees or node strengths. This is where our work breaks new ground --- it offers a comprehensive empirical analysis across all available connectomes, demonstrating that neurite length, synapse density, degree, and strength all follow the same lognormal law. Most important, it offers a mechanistic explanation based on neuronal branching that can account for the emergence of the lognormal distribution in the context of the connectome—as such it also represents the first network generative model that offers a lognormal outcome.

#### Bibliography

1. White, J. G., Southgate, E., Thomson, J. N. & Brenner, S. The Structure of the Nervous System of the Nematode Caenorhabditis elegans. *Philos. Trans. R. Soc. B Biol. Sci.* **314**, 1–340 (1986).
2. Szigeti, B., Gleeson, P., Vella, M., Khayrulin, S., Palyanov, A., Hokanson, J., Currie, M., Cantarelli, M., Idili, G. & Larson, S. OpenWorm: an open-science approach to modeling Caenorhabditis elegans. *Front. Comput. Neurosci.* **8**, (2014).
3. Varshney, L. R., Chen, B. L., Paniagua, E., Hall, D. H. & Chklovskii, D. B. Structural Properties of the Caenorhabditis elegans Neuronal Network. *PLoS Comput. Biol.* **7**, e1001066 (2011).
4. Cook, S. J., Jarrell, T. A., Brittin, C. A., Wang, Y., Bloniarz, A. E., Yakovlev, M. A., Nguyen, K. C. Q., Tang, L. T.-H., Bayer, E. A., Duerr, J. S., & others. Whole-animal connectomes of both Caenorhabditis elegans sexes. *Nature* **571**, 63 (2019).
5. Winding, M., Pedigo, B. D., Barnes, C. L., Patsolic, H. G., Park, Y., Kazimiers, T., Fushiki, A., Andrade, I. V., Khandelwal, A., Valdes-Aleman, J., Li, F., Randel, N., Barsotti, E., Correia, A., Fetter, R. D., Hartenstein, V., Priebe, C. E., Vogelstein, J. T., Cardona, A. & Zlatic, M. The connectome of an insect brain. *Science* **379**, eadd9330 (2023).
6. Xu, C. S., Januszewski, M., Lu, Z., Takemura, S., Hayworth, K. J., Huang, G., Shinomiya, K., Maitin-Shepard, J., Ackerman, D., Berg, S., Blakely, T., Bogovic, J., Clements, J., Dolafi, T., Hubbard, P., Kainmueller, D., Katz, W., Kawase, T., Khairy, K. A., Leavitt, L., Li, P. H., Lindsey, L., Neubarth, N., Olbris, D. J., Otsuna, H., Troutman, E. T., Umayam, L., Zhao, T., Ito, M., Goldammer, J., Wolff, T., Svirskas, R., Schlegel, P., Neace, E. R., Knecht, C. J., Alvarado, C. X., Bailey, D. A., Ballinger, S., Borycz, J. A., Canino, B. S., Cheatham, N., Cook, M., Dreher, M., Duclos, O., Eubanks, B., Fairbanks, K., Finley, S., Forknall, N., Francis, A., Hopkins, G. P., Joyce, E. M., Kim, S., Kirk, N. A., Kovalyak, J., Lauchie, S. A., Lohff, A., Maldonado, C., Manley, E. A., McLin, S., Mooney, C., Ndama, M., Ogundeyi, O., Okeoma, N., Ordish, C., Padilla, N., Patrick, C., Paterson, T., Phillips, E. E., Phillips, E. M., Rampally, N., Ribeiro, C., Robertson, M. K., Rymer, J. T., Ryan, S. M., Sammons, M., Scott, A. K., Scott, A. L., Shinomiya, A., Smith, C., Smith, K., Smith, N. L., Sobeski, M. A., Suleiman, A., Swift, J., Takemura, S., Talebi, I., Tarnogorska, D., Tenshaw, E., Tokhi, T., Walsh, J. J., Yang, T., Horne, J. A., Li, F., Parekh, R., Rivlin, P. K., Jayaraman, V., Ito, K., Saalfeld, S., George, R., Meinertzhagen, I., Rubin, G. M., Hess, H. F., Scheffer, L. K., Jain, V. & Plaza, S. M. A Connectome of the Adult *Drosophila* Central Brain. Preprint at https://doi.org/10.1101/2020.01.21.911859 (2020).
7. Takemura, S., Hayworth, K. J., Huang, G. B., Januszewski, M., Lu, Z., Marin, E. C., Preibisch, S., Xu, C. S., Bogovic, J., Champion, A. S., Cheong, H. S., Costa, M., Eichler, K., Katz, W., Knecht, C., Li, F., Morris, B. J., Ordish, C., Rivlin, P. K., Schlegel, P., Shinomiya, K., Stürner, T., Zhao, T., Badalamente, G., Bailey, D., Brooks, P., Canino, B. S., Clements, J., Cook, M., Duclos, O., Dunne, C. R., Fairbanks, K., Fang, S., Finley-May, S., Francis, A., George, R., Gkantia, M., Harrington, K., Hopkins, G. P., Hsu, J., Hubbard, P. M., Javier, A., Kainmueller, D., Korff, W., Kovalyak, J., Krzemiński, D., Lauchie, S. A., Lohff, A., Maldonado, C., Manley, E. A., Mooney, C., Neace, E., Nichols, M., Ogundeyi, O., Okeoma, N., Paterson, T., Phillips, E., Phillips, E. M., Ribeiro, C., Ryan, S. M., Rymer, J. T., Scott, A. K., Scott, A. L., Shepherd, D., Shinomiya, A., Smith, C., Smith, N., Suleiman, A., Takemura, S., Talebi, I., Tamimi, I. F., Trautman, E. T., Umayam, L., Walsh, J. J., Yang, T., Rubin, G. M., Scheffer, L. K., Funke, J., Saalfeld, S., Hess, H. F., Plaza, S. M., Card, G. M., Jefferis, G.

S. & Berg, S. A Connectome of the Male *Drosophila* Ventral Nerve Cord. Preprint at https://doi.org/10.1101/2023.06.05.543757 (2023).

8. Zheng, Z., Lauritzen, J. S., Perlman, E., Robinson, C. G., Nichols, M., Milkie, D., Torrens, O., Price, J., Fisher, C. B., Sharifi, N., Calle-Schuler, S. A., Kmecova, L., Ali, I. J., Karsh, B., Trautman, E. T., Bogovic, J. A., Hanslovsky, P., Jefferis, G. S. X. E., Kazhdan, M., Khairy, K., Saalfeld, S., Fetter, R. D. & Bock, D. D. A Complete Electron Microscopy Volume of the Brain of Adult Drosophila melanogaster. *Cell* **174**, 730-743.e22 (2018).
9. Dorkenwald, S., Matsliah, A., Sterling, A. R., Schlegel, P., Yu, S., McKellar, C. E., Lin, A., Costa, M., Eichler, K., Yin, Y., Silversmith, W., Schneider-Mizell, C., Jordan, C. S., Brittain, D., Halageri, A., Kuehner, K., Ogedengbe, O., Morey, R., Gager, J., Kruk, K., Perlman, E., Yang, R., Deutsch, D., Bland, D., Sorek, M., Lu, R., Macrina, T., Lee, K., Bae, J. A., Mu, S., Nehoran, B., Mitchell, E., Popovych, S., Wu, J., Jia, Z., Castro, M. A., Kemnitz, N., Ih, D., Bates, A. S., Eckstein, N., Funke, J., Collman, F., Bock, D. D., Jefferis, G. S. X. E., Seung, H. S., Murthy, M., The FlyWire Consortium, Lenizo, Z., Burke, A. T., Willie, K. P., Serafetinidis, N., Hadjerol, N., Willie, R., Silverman, B., Ocho, J. A., Bañez, J., Candilada, R. A., Kristiansen, A., Panes, N., Yadav, A., Tancontian, R., Serona, S., Dolorosa, J. I., Vinson, K. J., Garner, D., Salem, R., Dagohoy, A., Skelton, J., Lopez, M., Capdevila, L. S., Badalamente, G., Stocks, T., Pandey, A., Akiatan, D. J., Hebditch, J., David, C., Sapkal, D., Monungolh, S. M., Sane, V., Pielago, M. L., Albero, M., Laude, J., Dos Santos, M., Vohra, Z., Wang, K., Gogo, A. M., Kind, E., Mandahay, A. J., Martinez, C., Asis, J. D., Nair, C., Patel, D., Manaytay, M., Tamimi, I. F. M., Lim, C. A., Ampo, P. L., Pantujan, M. D., Javier, A., Bautista, D., Rana, R., Seguido, J., Parmar, B., Saguimpa, J. C., Moore, M., Pleijzier, M. W., Larson, M., Hsu, J., Joshi, I., Kakadiya, D., Braun, A., Pilapil, C., Gkantia, M., Parmar, K., Vanderbeck, Q., Salgarella, I., Dunne, C., Munnelly, E., Kang, C. H., Lörsch, L., Lee, J., Kmecova, L., Sancer, G., Baker, C., Joroff, J., Calle, S., Patel, Y., Sato, O., Fang, S., Salocot, J., Salman, F., Molina-Obando, S., Brooks, P., Bui, M., Lichtenberger, M., Tamboboy, E., Molloy, K., Santana-Cruz, A. E., Hernandez, A., Yu, S., Diwan, A., Patel, M., Aiken, T. R., Morejohn, S., Koskela, S., Yang, T., Lehmann, D., Chojetzki, J., Sisodiya, S., Koolman, S., Shiu, P. K., Cho, S., Bast, A., Reicher, B., Blanquart, M., Houghton, L., Choi, H., Ioannidou, M., Collie, M., Eckhardt, J., Gorko, B., Guo, L., Zheng, Z., Poh, A., Lin, M., Taisz, I., Murfin, W., Díez, Á. S., Reinhard, N., Gibb, P., Patel, N., Kumar, S., Yun, M., Wang, M., Jones, D., Encarnacion-Rivera, L., Oswald, A., Jadia, A., Erginkaya, M., Drummond, N., Walter, L., Tastekin, I., Zhong, X., Mabuchi, Y., Figueroa Santiago, F. J., Verma, U., Byrne, N., Kunze, E., Crahan, T., Margossian, R., Kim, H., Georgiev, I., Szorenyi, F., Adachi, A., Bargeron, B., Stürner, T., Demarest, D., Gür, B., Becker, A. N., Turnbull, R., Morren, A., Sandoval, A., Moreno-Sanchez, A., Pacheco, D. A., Samara, E., Croke, H., Thomson, A., Laughland, C., Dutta, S. B., De Antón, P. G. A., Huang, B., Pujols, P., Haber, I., González-Segarra, A., Choe, D. T., Lukyanova, V., Mancini, N., Liu, Z., Okubo, T., Flynn, M. A., Vitelli, G., Laturney, M., Li, F., Cao, S., Manyari-Diaz, C., Yim, H., Duc Le, A., Maier, K., Yu, S., Nam, Y., Bąba, D., Abusaif, A., Francis, A., Gayk, J., Huntress, S. S., Barajas, R., Kim, M., Cui, X., Sterne, G. R., Li, A., Park, K., Dempsey, G., Mathew, A., Kim, J., Kim, T., Wu, G., Dhawan, S., Brotas, M., Zhang, C., Bailey, S., Del Toro, A., Yang, R., Gerhard, S., Champion, A., Anderson, D. J., Behnia, R., Bidaye, S. S., Borst, A., Chiappe, E., Colodner, K. J., Dacks, A., Dickson, B., Garcia, D., Hampel, S., Hartenstein, V., Hassan, B., Helfrich-Forster, C., Huetteroth, W., Kim, J., Kim, S. S., Kim, Y.-J., Kwon, J. Y., Lee, W.-C., Linneweber, G. A., Maimon, G., Mann, R., Noselli, S.,

Pankratz, M., Prieto-Godino, L., Read, J., Reiser, M., Von Reyn, K., Ribeiro, C., Scott, K., Seeds, A. M., Selcho, M., Silies, M., Simpson, J., Waddell, S., Wernet, M. F., Wilson, R. I., Wolf, F. W., Yao, Z., Yapici, N. & Zandawala, M. Neuronal wiring diagram of an adult brain. *Nature* **634**, 124–138 (2024).

10. Schlegel, P., Yin, Y., Bates, A. S., Dorkenwald, S., Eichler, K., Brooks, P., Han, D. S., Gkantia, M., Dos Santos, M., Munnelly, E. J., Badalamente, G., Serratosa Capdevila, L., Sane, V. A., Fragniere, A. M. C., Kiassat, L., Pleijzier, M. W., Stürner, T., Tamimi, I. F. M., Dunne, C. R., Salgarella, I., Javier, A., Fang, S., Perlman, E., Kazimiers, T., Jagannathan, S. R., Matsliah, A., Sterling, A. R., Yu, S., McKellar, C. E., FlyWire Consortium, Kruk, K., Bland, D., Lenizo, Z., Burke, A. T., Willie, K. P., Bates, A. S., Serafetinidis, N., Hadjerol, N., Willie, R., Silverman, B., Ocho, J. A., Bañez, J., Candilada, R. A., Gager, J., Kristiansen, A., Panes, N., Yadav, A., Tancontian, R., Serona, S., Dolorosa, J. I., Vinson, K. J., Garner, D., Salem, R., Dagohoy, A., Skelton, J., Lopez, M., Stocks, T., Pandey, A., Akiatan, D. J., Hebditch, J., David, C., Sapkal, D., Monungolh, S. M., Sane, V., Pielago, M. L., Albero, M., Laude, J., Dos Santos, M., Deutsch, D., Vohra, Z., Wang, K., Gogo, A. M., Kind, E., Mandahay, A. J., Martinez, C., Asis, J. D., Nair, C., Patel, D., Manaytay, M., Lim, C. A., Ampo, P. L., Pantujan, M. D., Bautista, D., Rana, R., Seguido, J., Parmar, B., Saguimpa, J. C., Moore, M., Pleijzier, M. W., Larson, M., Hsu, J., Joshi, I., Kakadiya, D., Braun, A., Pilapil, C., Parmar, K., Vanderbeck, Q., Dunne, C., Munnelly, E., Kang, C. H., Lörsch, L., Lee, J., Kmecova, L., Sancer, G., Baker, C., Joroff, J., Calle, S., Patel, Y., Sato, O., Salocot, J., Salman, F., Molina-Obando, S., Bui, M., Lichtenberger, M., Tamboboy, E., Molloy, K., Santana-Cruz, A. E., Hernandez, A., Yu, S., Sorek, M., Diwan, A., Patel, M., Aiken, T. R., Morejohn, S., Koskela, S., Yang, T., Lehmann, D., Chojetzki, J., Sisodiya, S., Koolman, S., Shiu, P. K., Cho, S., Bast, A., Reicher, B., Blanquart, M., Houghton, L., Choi, H., Ioannidou, M., Collie, M., Eckhardt, J., Gorko, B., Guo, L., Zheng, Z., Poh, A., Lin, M., Taisz, I., Murfin, W., Díez, Á. S., Reinhard, N., Gibb, P., Patel, N., Kumar, S., Yun, M., Wang, M., Jones, D., Encarnacion-Rivera, L., Oswald, A., Jadia, A., Erginkaya, M., Drummond, N., Walter, L., Tastekin, I., Zhong, X., Mabuchi, Y., Figueroa Santiago, F. J., Verma, U., Byrne, N., Kunze, E., Crahan, T., Margossian, R., Kim, H., Georgiev, I., Szorenyi, F., Adachi, A., Bargeron, B., Stürner, T., Demarest, D., Gür, B., Becker, A. N., Turnbull, R., Morren, A., Sandoval, A., Moreno-Sanchez, A., Pacheco, D. A., Samara, E., Croke, H., Thomson, A., Laughland, C., Dutta, S. B., De Antón, P. G. A., Huang, B., Pujols, P., Haber, I., González-Segarra, A., Lin, A., Choe, D. T., Lukyanova, V., Mancini, N., Liu, Z., Okubo, T., Flynn, M. A., Vitelli, G., Laturney, M., Li, F., Cao, S., Manyari-Diaz, C., Yim, H., Duc Le, A., Maier, K., Yu, S., Nam, Y., Bąba, D., Abusaif, A., Francis, A., Gayk, J., Huntress, S. S., Barajas, R., Kim, M., Cui, X., Sterling, A. R., Sterne, G. R., Li, A., Park, K., Dempsey, G., Mathew, A., Kim, J., Kim, T., Wu, G., Dhawan, S., Brotas, M., Zhang, C., Bailey, S., Del Toro, A., Lee, K., Macrina, T., Schneider-Mizell, C., Popovych, S., Ogedengbe, O., Yang, R., Halageri, A., Silversmith, W., Gerhard, S., Champion, A., Eckstein, N., Ih, D., Kemnitz, N., Castro, M., Jia, Z., Wu, J., Mitchell, E., Nehoran, B., Mu, S., Bae, J. A., Lu, R., Morey, R., Kuehner, K., Brittain, D., Jordan, C. S., Anderson, D. J., Behnia, R., Bidaye, S. S., Borst, A., Chiappe, E., Collman, F., Colodner, K. J., Dacks, A., Dickson, B., Funke, J., Garcia, D., Hampel, S., Hartenstein, V., Hassan, B., Helfrich-Forster, C., Huetteroth, W., Kim, J., Kim, S. S., Kim, Y.-J., Kwon, J. Y., Lee, W.-C., Linneweber, G. A., Maimon, G., Mann, R., Noselli, S., Pankratz, M., Prieto-Godino, L., Read, J., Reiser, M., Von Reyn, K., Ribeiro, C., Scott, K., Seeds, A. M., Selcho, M., Silies, M., Simpson, J., Waddell, S., Wernet, M. F., Wilson, R. I.,

Wolf, F. W., Yao, Z., Yapici, N., Zandawala, M., Costa, M., Seung, H. S., Murthy, M., Hartenstein, V., Bock, D. D. & Jefferis, G. S. X. E. Whole-brain annotation and multi-connectome cell typing of Drosophila. *Nature* **634**, 139–152 (2024).
11. Vishwanathan, A., Daie, K., Ramirez, A. D., Lichtman, J. W., Aksay, E. R. F. & Seung, H. S. Electron Microscopic Reconstruction of Functionally Identified Cells in a Neural Integrator. *Curr. Biol.* **27**, 2137-2147.e3 (2017).
12. Vishwanathan, A., Ramirez, A. D., Wu, J., Sood, A., Yang, R., Kemnitz, N., Ih, D., Turner, N., Lee, K., Tartavull, I., Silversmith, W. M., Jordan, C. S., David, C., Bland, D., Goldman, M. S., Aksay, E. R. F., Seung, H. S., & the Eyewirers. Predicting modular functions and neural coding of behavior from a synaptic wiring diagram. Preprint at https://doi.org/10.1101/2020.10.28.359620 (2020).
13. Schneider-Mizell, C. M., Bodor, A. L., Brittain, D., Buchanan, J., Bumbarger, D. J., Elabbady, L., Gamlin, C., Kapner, D., Kinn, S., Mahalingam, G., Seshamani, S., Suckow, S., Takeno, M., Torres, R., Yin, W., Dorkenwald, S., Bae, J. A., Castro, M. A., Halageri, A., Jia, Z., Jordan, C., Kemnitz, N., Lee, K., Li, K., Lu, R., Macrina, T., Mitchell, E., Mondal, S. S., Mu, S., Nehoran, B., Popovych, S., Silversmith, W., Turner, N. L., Wong, W., Wu, J., The MICrONS Consortium, Reimer, J., Tolias, A. S., Seung, H. S., Reid, R. C., Collman, F. & Maçarico Da Costa, N. Cell-type-specific inhibitory circuitry from a connectomic census of mouse visual cortex. Preprint at https://doi.org/10.1101/2023.01.23.525290 (2023).
14. Liu, L., Yun, Z., Manubens-Gil, L., Chen, H., Xiong, F., Dong, H., Zeng, H., Hawrylycz, M., Ascoli, G. A. & Peng, H. Connectivity of single neurons classifies cell subtypes in mouse brains. *Nat. Methods* **22**, 861–873 (2025).
15. Peng, H., Xie, P., Liu, L., Kuang, X., Wang, Y., Qu, L., Gong, H., Jiang, S., Li, A., Ruan, Z., Ding, L., Yao, Z., Chen, C., Chen, M., Daigle, T. L., Dalley, R., Ding, Z., Duan, Y., Feiner, A., He, P., Hill, C., Hirokawa, K. E., Hong, G., Huang, L., Kebede, S., Kuo, H.-C., Larsen, R., Lesnar, P., Li, L., Li, Q., Li, X., Li, Y., Li, Y., Liu, A., Lu, D., Mok, S., Ng, L., Nguyen, T. N., Ouyang, Q., Pan, J., Shen, E., Song, Y., Sunkin, S. M., Tasic, B., Veldman, M. B., Wakeman, W., Wan, W., Wang, P., Wang, Q., Wang, T., Wang, Y., Xiong, F., Xiong, W., Xu, W., Ye, M., Yin, L., Yu, Y., Yuan, J., Yuan, J., Yun, Z., Zeng, S., Zhang, S., Zhao, S., Zhao, Z., Zhou, Z., Huang, Z. J., Esposito, L., Hawrylycz, M. J., Sorensen, S. A., Yang, X. W., Zheng, Y., Gu, Z., Xie, W., Koch, C., Luo, Q., Harris, J. A., Wang, Y. & Zeng, H. Morphological diversity of single neurons in molecularly defined cell types. *Nature* **598**, 174–181 (2021).
16. Liu, Y., Jiang, S., Li, Y., Zhao, S., Yun, Z., Zhao, Z.-H., Zhang, L., Wang, G., Chen, X., Manubens-Gil, L., Hang, Y., Gong, Q., Li, Y., Qian, P., Qu, L., Garcia-Forn, M., Wang, W., De Rubeis, S., Wu, Z., Osten, P., Gong, H., Hawrylycz, M., Mitra, P., Dong, H., Luo, Q., Ascoli, G. A., Zeng, H., Liu, L. & Peng, H. Neuronal diversity and stereotypy at multiple scales through whole brain morphometry. *Nat. Commun.* **15**, 10269 (2024).
17. Gao, L., Liu, S., Gou, L., Hu, Y., Liu, Y., Deng, L., Ma, D., Wang, H., Yang, Q., Chen, Z., Liu, D., Qiu, S., Wang, X., Wang, D., Wang, X., Ren, B., Liu, Q., Chen, T., Shi, X., Yao, H., Xu, C., Li, C. T., Sun, Y., Li, A., Luo, Q., Gong, H., Xu, N. & Yan, J. Single-neuron projectome of mouse prefrontal cortex. *Nat. Neurosci.* **25**, 515–529 (2022).
18. Shapson-Coe, A., Januszewski, M., Berger, D. R., Pope, A., Wu, Y., Blakely, T., Schalek, R. L., Li, P. H., Wang, S., Maitin-Shepard, J., Karlupia, N., Dorkenwald, S., Sjostedt, E., Leavitt, L., Lee, D., Troidl, J., Collman, F., Bailey, L., Fitzmaurice, A., Kar, R., Field, B., Wu, H., Wagner-Carena, J., Aley, D., Lau, J., Lin, Z., Wei, D., Pfister, H., Peleg, A., Jain, V.

& Lichtman, J. W. A petavoxel fragment of human cerebral cortex reconstructed at nanoscale resolution. *Science* **384**, eadk4858 (2024).
19. Krapivsky, P. L., Redner, S. & Leyvraz, F. Connectivity of Growing Random Networks. *Phys. Rev. Lett.* **85**, 4629–4632 (2000).
20. Mitzenmacher, M. A Brief History of Generative Models for Power Law and Lognormal Distributions. *Internet Math.* **1**, 226–251 (2004).
21. Voitalov, I., Van Der Hoorn, P., Van Der Hofstad, R. & Krioukov, D. Scale-free networks well done. *Phys. Rev. Res.* **1**, 033034 (2019).
22. Serafino, M., Cimini, G., Maritan, A., Rinaldo, A., Suweis, S., Banavar, J. R. & Caldarelli, G. True scale-free networks hidden by finite size effects. *Proc. Natl. Acad. Sci.* **118**, e2013825118 (2021).
23. Broido, A. D. & Clauset, A. Scale-free networks are rare. *Nat. Commun.* **10**, 1017 (2019).
24. Caldarelli, G. *Scale-Free Networks: Complex Webs in Nature and Technology*. (Oxford Univ. Press, Oxford, 2010).
25. Vuong, Q. H. Likelihood Ratio Tests for Model Selection and Non-Nested Hypotheses. *Econometrica* **57**, 307 (1989).
26. Clauset, A., Shalizi, C. R. & Newman, M. E. J. Power-Law Distributions in Empirical Data. *SIAM Rev.* **51**, 661–703 (2009).
27. Erdös, P. & Rényi, A. On evolution of random graphs. *Publ. Math. Inst. Hung. Acad. Sci.* 17–61 (1960) doi:10.1.1.153.5943.
28. Barabasi, A.-L. & Albert, R. Emergence of Scaling in Random Networks. *Science* **286**, 509–512 (1999).
29. Barabási, A. L. *Network Science*. (Cambridge University Press, 2016).
30. Barabási, A.-L. & Albert, R. Emergence of scaling in random networks. *Science* **286**, 509–512 (1999).
31. Cohen, R. & Havlin, S. Scale-Free Networks are Ultrasmall. *Phys. Rev. Lett.* **90**, 058701 (2003).
32. Bollobás, B., Riordan, O., Spencer, J. & Tusnády, G. The degree sequence of a scale-free random graph process. *Random Struct. Algorithms* **18**, 279–290 (2001).
33. Dorogovtsev, S. N. & Mendes, J. F. F. *The Nature of Complex Networks*. (Oxford university press, Oxford, United Kingdom, 2022).
34. Albert, R. & Barabási, A.-L. Statistical mechanics of complex networks. *Rev. Mod. Phys.* **74**, 47–97 (2002).
35. Sheridan, P. & Onodera, T. A Preferential Attachment Paradox: How Preferential Attachment Combines with Growth to Produce Networks with Log-normal In-degree Distributions. *Sci. Rep.* **8**, 2811 (2018).
36. Redner, S. Citation Statistics from 110 Years of *Physical Review*. *Phys. Today* **58**, 49–54 (2005).
37. Kim, M. & Leskovec, J. Multiplicative Attribute Graph Model of Real-World Networks. *Internet Math.* **8**, 113–160 (2012).
38. Ferreira Castro, A. & Cardona, A. Invariant synaptic density links neuronal activity stability and wiring optimisation principles across species. Preprint at https://doi.org/10.1101/2024.07.18.604056 (2024).
39. Otopalik, A. G., Goeritz, M. L., Sutton, A. C., Brookings, T., Guerini, C. & Marder, E. Sloppy morphological tuning in identified neurons of the crustacean stomatogastric ganglion. *eLife* **6**, (2017).

40. Scorcioni, R., Polavaram, S. & Ascoli, G. L-Measure: a web-accessible tool for the analysis, comparison and search of digital reconstructions of neuronal morphologies. *Nat Protoc* **3**, 866–876 (2008).
41. Kanari, L., Dłotko, P., Scolamiero, M., Levi, R., Shillcock, J., Hess, K. & Markram, H. A topological representation of branching neuronal morphologies. *Neuroinformatics* **16**, 3–13 (2018).
42. Kanari, L., Ramaswamy, S., Shi, Y., Morand, S., Meystre, J., Perin, R., Abdellah, M., Wang, Y., Hess, K. & Markram, H. Objective morphological classification of neocortical pyramidal cells. *Cereb. Cortex* **29**, 1719–1735 (2019).
43. Liao, M., Bird, A. D., Cuntz, H. & Howard, J. Topology recapitulates morphogenesis of neuronal dendrites. *Cell Rep.* **42**, (2023).
44. Bird, A. D. & Cuntz, H. Dissecting sholl analysis into its functional components. *Cell Rep.* **27**, 3081–3096 (2019).
45. Snider, J., Pillai, A. & Stevens, C. F. A universal property of axonal and dendritic arbors. *Neuron* **66**, 45–56 (2010).
46. Meng, X., Piazza, B., Both, C., Barzel, B. & Barabási, A.-L. A Manifold Minimisation Principle for Physical Networks. (2025).
47. Van Pelt, J., Uylings, H., Verwer, R., Pentney, R. & Woldenberg, M. Tree asymmetry—a sensitive and practical measure for binary topological trees. *Bull Math Biol* **54**, 759–784 (1992).
48. van Pelt, J., Schierwagen, A. & Uylings, H. B. Modeling dendritic morphological complexity of deep layer cat superior colliculus neurons. *Neurocomputing* **38**, 403–408 (2001).
49. Zeng, H. & Sanes, J. R. Neuronal cell-type classification: challenges, opportunities and the path forward. *Nat. Rev. Neurosci.* **18**, 530–546 (2017).
50. Cuntz, H., Forstner, F., Borst, A., Häusser, M., Graham, R. L. & Hell, P. On the history of the minimum spanning tree problem. *PLOS Comput Biol* **7**, 43–57 (1985).
51. Ferreira Castro, A., Baltruschat, L., Stürner, T., Bahrami, A., Jedlicka, P., Tavosanis, G. & Cuntz, H. Achieving functional neuronal dendrite structure through sequential stochastic growth and retraction. *Elife* **9**, e60920 (2020).
52. Van Ooyen, A. & Willshaw, D. J. Competition for neurotrophic factor in the development of nerve connections. *Proc. R. Soc. B Biol. Sci.* **266**, 883–892 (1999).
53. Uylings, H. B. & Van Pelt, J. Measures for quantifying dendritic arborizations. *Netw. Comput. Neural Syst.* **13**, 397 (2002).
54. Ascoli, G. A. & Krichmar, J. L. L-neuron: A modeling tool for the efficient generation and parsimonious description of dendritic morphology. *Neurocomputing* **32–33**, 1003–1011 (2000).
55. Nanda, S., Das, R., Bhattacharjee, S., Cox, D. N. & Ascoli, G. A. Morphological determinants of dendritic arborization neurons in Drosophila larva. *Brain Struct. Funct.* **223**, 1–14 (2017).
56. Shree, S., Sutradhar, S., Trottier, O., Tu, Y., Liang, X. & Howard, J. Dynamic instability of dendrite tips generates the highly branched morphologies of sensory neurons. *Sci. Adv.* **8**, eabn0080 (2022).
57. Memelli, H., Torben-Nielsen, B. & Kozloskiy, J. Self-referential forces are suffcient to explain different dendritic morphologies. *Front. Neuroinformatics* **6**, (2013).
58. Luczak, A. Spatial embedding of neuronal trees modeled by diffusive growth. *J. Neurosci. Methods* **157**, 132–141 (2006).

59. Stepanyants, A. & Chklovskii, D. B. Neurogeometry and potential synaptic connectivity. *Trends Neurosci.* **28**, 387–94 (2005).
60. Kuznetsov, A. V. Comparison of active transport in neuronal axons and dendrites. **228**, 195–202 (2010).
61. Kobayashi, T., Terajima, K., Nozumi, M., Igarashi, M. & Akazawa, K. A stochastic model of neuronal growth cone guidance regulated by multiple sensors. *J. Theor. Biol.* **266**, 712–722 (2010).
62. Suter, D. M. & Miller, K. E. The emerging role of forces in axonal elongation. *Prog. Neurobiol.* **94**, 91–101 (2011).
63. Athreya, K. B. & Ney, P. E. *Branching Processes*. (Springer Berlin Heidelberg, Berlin, Heidelberg, 1972). doi:10.1007/978-3-642-65371-1.
64. Kimmel, M. & Axelrod, D. E. *Branching Processes in Biology*. (Springer, New York).
65. Kossovsky, A. E. *Benford's Law: Theory, the General Law of Relative Quantities, and Forensic Fraud Detection Applications*. (World Scientific, Singapore, 2015).
66. Mitchell, R. L. Permanence of the Log-Normal Distribution*. *J. Opt. Soc. Am.* **58**, 1267 (1968).
67. Dufresne, D. Sums of lognormals. in *Actuarial Research Conference* 1–6 (2008).
68. Bianconi, G. & Barabási, A.-L. Bose-Einstein Condensation in Complex Networks. *Phys. Rev. Lett.* **86**, 5632–5635 (2001).
69. Bianconi, G. & Barabási, A.-L. Competition and multiscaling in evolving networks. *Europhys. Lett. EPL* **54**, 436–442 (2001).
70. Pósfai, M., Szegedy, B., Bačić, I., Blagojević, L., Abért, M., Kertész, J., Lovász, L. & Barabási, A.-L. Impact of physicality on network structure. *Nat. Phys.* **20**, 142–149 (2024).
71. Jagers, P. Branching processes with dependence but homogeneous growth. *Ann. Appl. Probab.* **9**, (1999).
72. Lynn, C. W., Holmes, C. M. & Palmer, S. E. Heavy-tailed neuronal connectivity arises from Hebbian self-organization. *Nat. Phys.* **20**, 484–491 (2024).
73. Cirunay, M., Ódor, G., Papp, I. & Deco, G. Scale-free behavior of weight distributions of connectomes. https://doi.org/10.48550/ARXIV.2407.17220 (2024) doi:10.48550/ARXIV.2407.17220.
74. Lin, A., Yang, R., Dorkenwald, S., Matsliah, A., Sterling, A. R., Schlegel, P., Yu, S., McKellar, C. E., Costa, M., Eichler, K., Bates, A. S., Eckstein, N., Funke, J., Jefferis, G. S. X. E. & Murthy, M. Network statistics of the whole-brain connectome of Drosophila. *Nature* **634**, 153–165 (2024).
75. Cirunay, M. T., Batac, R. C. & Ódor, G. Learning and criticality in a self-organizing model of connectome growth. *Sci. Rep.* **15**, 31890 (2025).
76. Towlson, E. K., Vértes, P. E., Ahnert, S. E., Schafer, W. R. & Bullmore, E. T. The Rich Club of the *C. elegans* Neuronal Connectome. *J. Neurosci.* **33**, 6380–6387 (2013).
77. Van Den Heuvel, M. P. & Sporns, O. Rich-Club Organization of the Human Connectome. *J. Neurosci.* **31**, 15775–15786 (2011).
78. Colizza, V., Flammini, A., Serrano, M. A. & Vespignani, A. Detecting rich-club ordering in complex networks. *Nat. Phys.* **2**, 110–115 (2006).
79. Zhang, Y. & Ng, S. T. Unveiling the rich-club phenomenon in urban mobility networks through the spatiotemporal characteristics of passenger flow. *Phys. Stat. Mech. Its Appl.* **584**, 126377 (2021).

80. In 'T Veld, D. & Van Lelyveld, I. Finding the core: Network structure in interbank markets. *J. Bank. Finance* **49**, 27–40 (2014).
81. Van Den Heuvel, M. P. & Sporns, O. Network hubs in the human brain. *Trends Cogn. Sci.* **17**, 683–696 (2013).
82. Becker, N. On parametric estimation for mortal branching processes. *Biometrika* **61**, 393–399 (1974).
83. Bickel, P. J., Chen, A. & Levina, E. The method of moments and degree distributions for network models. https://doi.org/10.48550/ARXIV.1202.5101 (2012) doi:10.48550/ARXIV.1202.5101.
84. Buzsáki, G. & Mizuseki, K. The log-dynamic brain: how skewed distributions affect network operations. *Nat. Rev. Neurosci.* **15**, 264–278 (2014).
85. Buzsáki, G. *The Brain from inside Out*. (Oxford University Press, New York, NY, 2021).
86. Nigam, S., Shimono, M., Ito, S., Yeh, F.-C., Timme, N., Myroshnychenko, M., Lapish, C. C., Tosi, Z., Hottowy, P., Smith, W. C., Masmanidis, S. C., Litke, A. M., Sporns, O. & Beggs, J. M. Rich-Club Organization in Effective Connectivity among Cortical Neurons. *J. Neurosci.* **36**, 670–684 (2016).
87. Liu, Y., Seguin, C., Betzel, R. F., Han, D., Akarca, D., Di Biase, M. A. & Zalesky, A. A generative model of the connectome with dynamic axon growth. *Netw. Neurosci.* **8**, 1192–1211 (2024).
88. Smith, K. M. Explaining the emergence of complex networks through log-normal fitness in a Euclidean node similarity space. *Sci. Rep.* **11**, 1976 (2021).
